# Structural basis of end processing in the nucleosome by polynucleotide kinase-phosphatase

**DOI:** 10.64898/2026.08.12.743989

**Authors:** Daniel J. Boesch, Daria K. Majewska, Nadia I. Martin, John J. Evans, Tyler M. Weaver

## Abstract

Genomic DNA is packaged into chromatin through a fundamental repeating unit known as the nucleosome core particle. Chromatinized DNA is under constant assault from endogenous and exogenous sources of damage, which must be effectively repaired to preserve genome stability. Single-strand breaks (SSBs) with chemically heterogeneous DNA ends are one of the most prevalent forms of genomic DNA damage. These SSBs must be enzymatically processed prior to downstream gap-filling DNA synthesis and/or nick ligation during single-strand break repair (SSBR). Polynucleotide kinase-phosphatase (PNKP) is a multifunctional end-processing enzyme that possesses two catalytic activities important for converting non-ligatable SSBs into ligatable SSBs. To date, a mechanistic description for how PNKP processes non-ligatable SSBs in the context of chromatin to initiate SSBR remains undefined. Here, we utilize a combination of biochemical assays and cryogenic electron microscopy (cryo-EM) to define the structural basis of end processing in the nucleosome by PNKP. Cryo-EM structures of PNKP engaged with non-ligatable SSBs at three unique positions within the nucleosome reveal that PNKP locally deforms nucleosomal DNA to reposition the SSBs into the kinase and phosphatase active sites, providing a structural basis for the efficient processing of SSBs throughout the nucleosome. Additional cryo-EM structures reveal the PNKP FHA domain also engages the nucleosome acidic patch during non-ligatable SSB recognition, which accelerates the processing of non-ligatable SSBs in the nucleosome. Together, these findings provide important mechanistic insight into the initial end processing step of chromatin-based SSBR.

## Introduction

Genomic DNA is packaged into chromatin through a fundamental repeating structural unit called the nucleosome core particle. The nucleosome core particle consists of approximately 147 base-pairs of DNA wrapped around an octameric histone assembly containing two copies each of histones H2A, H2B, H3, and H4^1^, which functions as a structural scaffold and regulatory barrier governing access to the underlying genomic DNA. Chromatinized genomic DNA is continuously exposed to endogenous and exogenous stress that generates DNA damage^2–12^, which must be faithfully repaired to preserve genomic integrity.

Among the most prevalent forms of genomic damage are single-strand breaks (SSBs), which arise from reactive oxygen species, ionizing radiation, and as intermediates during DNA excision repair^13,14^. Failure to repair SSBs results in genome instability as well as neurodevelopmental and/or neurodegenerative disorders^15–18^. SSBs commonly form with chemically diverse DNA ends that lack the canonical 5′-phosphate and 3′-hydroxyl termini required for gap-filling DNA synthesis and/or nick ligation^19–21^. Collectively, these chemically diverse SSBs are repaired through the single-strand break repair (SSBR) pathway^19–21^. SSBR is initiated by one of several end-processing enzymes that convert non-canonical SSBs into SSBs bearing canonical 5′-phosphate and 3′-hydroxyl termini. Depending on the nature of the break (i.e., gap or nick), the canonical 5′-phosphate and 3′-hydroxyl termini then undergo gap-filling DNA synthesis by DNA Polymerase β and/or nick ligation by DNA Ligase I/IIIα to restore continuity of the DNA backbone. In addition to the core enzymatic machinery, SSBR is facilitated by poly(ADP-ribose) polymerases (PARP1/2/3), which act as single-strand break sensors^22–24^ that mediate the ADP-ribosylation-dependent recruitment of the scaffolding protein X-ray repair cross complementing protein 1 (XRCC1)^25–29^. XRCC1 directly interacts with multiple end processing enzymes^30–34^, DNA Polymerase β^35,36^, and DNA Ligase IIIα^37,38^, which enables XRCC1 to coordinate the assembly of SSBR enzymes at cellular SSBs and facilitate timely repair^30,39–41^. Importantly, SSBR is a genome surveillance pathway that must deal with SSBs that arise throughout the chromatinized genome. While prior work has unraveled the molecular basis of gap-filling DNA synthesis^42^ and nick ligation^43^ in the nucleosome during chromatin-based SSBR, the molecular details of initial end processing in the nucleosome remain undefined.

Polynucleotide kinase-phosphatase (PNKP) is a bifunctional end processing enzyme that converts non-canonical 3′-phosphate and 5′-hydroxyl termini to canonical 5′-phosphate and 3′-hydroxyl termini^44–47^. PNKP adopts a modular three-domain architecture comprised of an N-terminal Forkhead-associated (FHA) domain that mediates an interaction with XRCC1^34,48^, and a C-terminal catalytic core containing a phosphatase and kinase domain that dephosphorylates 3′-phosphates and phosphorylates 5′-hydroxyls^45–51^, respectively. Prior biochemical and structural studies have provided extensive molecular insight into how PNKP processes non-ligatable SSBs in the context of non-chromatinized DNA^45–53^. These studies revealed that recognition and processing of the non-ligatable SSBs in DNA is mediated by a helical wedge in the PNKP kinase domain that inserts into the duplex at the nick site to deform the DNA and reposition the 3′-phosphate and 5′-hydroxyl into the phosphatase and kinase active sites, respectively. Despite this detailed mechanistic description of end processing in non-chromatinized DNA, how PNKP recognizes and processes non-ligatable SSBs in the nucleosome remains unknown. Here, we combine biochemical assays and cryogenic electron microscopy (cryo-EM) to define the structural basis of end processing in the nucleosome by PNKP, providing important mechanistic insight into the first enzymatic step of chromatin-based SSBR

## Results

### PNKP efficiently binds and processes non-ligatable nicks in the nucleosome

To obtain insight into the mechanism used by PNKP to catalyze end processing in the nucleosome, we initially designed five unique Widom 601 strong positioning DNA sequences with a nick containing a 3′-phosphate and 5′-hydroxyl (i.e., non-ligatable nick) (Fig. 1A). Each non-ligatable nick is located in a solvent-exposed rotational orientation at a unique translational position in the nucleosomal DNA at SHL−5, SHL−4, SHL−3, SHL−2, and SHL−1 (Fig. 1B). Herein, these nucleosome (NCP) substrates are referred to as Nick-NCP−5, Nick-NCP−4, Nick-NCP−3, Nick-NCP−2, and Nick-NCP−1. These positions were chosen as solvent-exposed locations in the nucleosome are highly prone to direct single-strand breaks formed via the oxidation-induced disintegration of the sugar phosphate backbone^54,55^. Each Nick-NCP was then reconstituted using recombinant human histones (SI Appendix, Fig. S1), and the nucleosome substrates were used to determine how efficiently PNKP binds and processes non-ligatable nicks in the nucleosome.

**Fig. 1:**
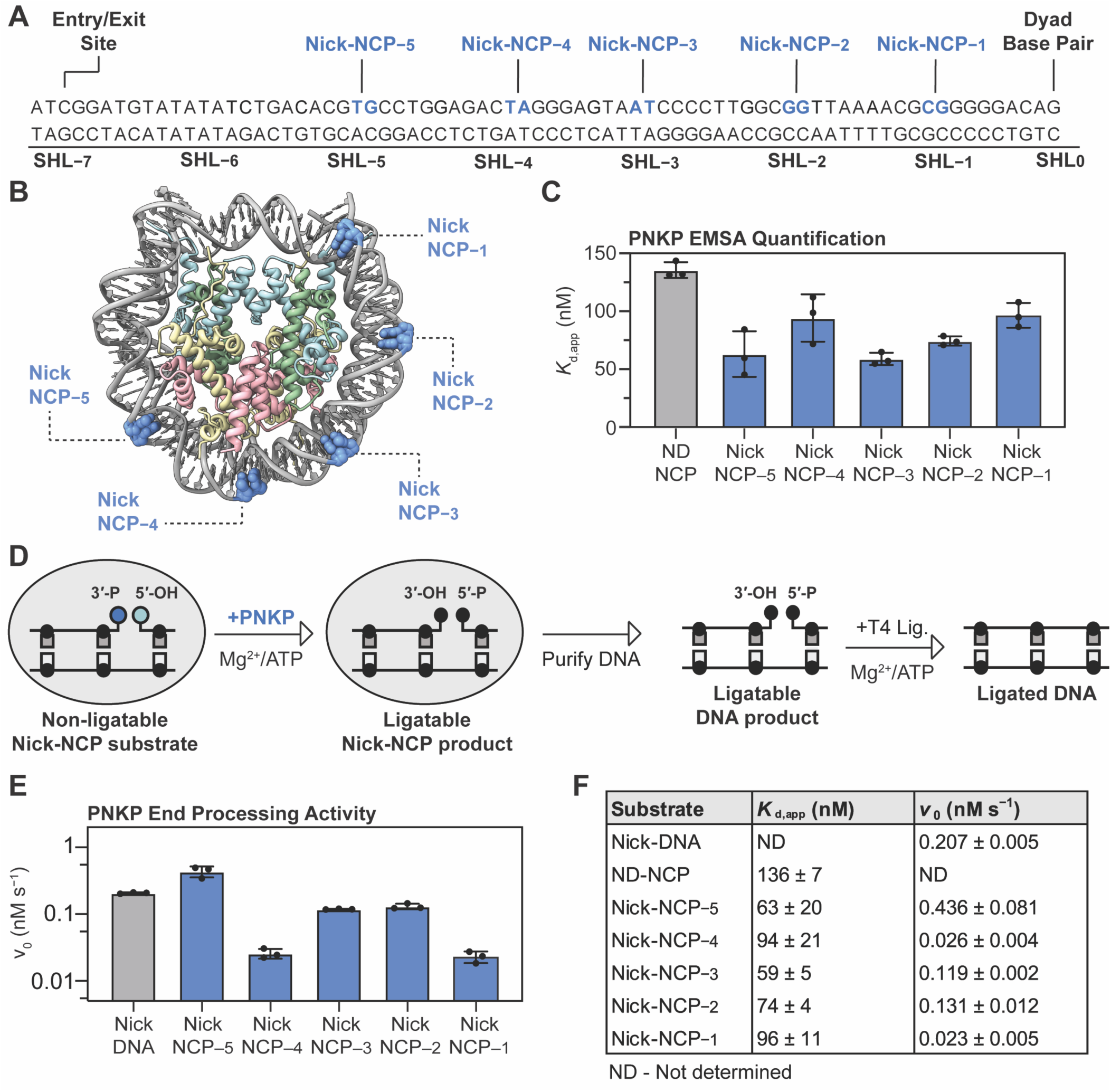
PNKP efficiently binds and processes non-ligatable nicks in the nucleosome. **(A)** Diagram of the 601 strong positioning sequence highlighting the position of each individual non-ligatable nick in blue. **(B)** Nucleosome core particle (adapted from PDB:9DWF) highlighting nicked positions at SHL−5, SHL−4, SHL−3, SHL−2, and SHL−1. **(C)** Apparent binding affinities (*K*_d, app_) of PNKP for the non-damaged NCP (ND-NCP), Nick-NCP−5, Nick-NCP−4, Nick-NCP−3, Nick-NCP−2, and Nick-NCP−1 obtained from the electrophoretic mobility shift assays. The data points represent the mean ± standard deviation from three independent replicate experiments (see SI Appendix, Fig. S2 for associated data). **(D)** Diagram outlining the ligase-coupled PNKP product formation assay. **(E)** Quantification of the the initial velocities (*v*_0_) obtained for PNKP end processing on Nick-DNA, Nick-NCP−5, Nick-NCP−4, Nick-NCP−3, Nick-NCP−2, and Nick-NCP−1. The data points represent the mean ± standard deviation from three independent replicate experiments (see SI Appendix, Fig. S3 for associated data). **(F)** Summary of apparent binding affinities (*K*_d, app_) and initial velocities (*v*_0_) obtained for PNKP and each respective substrate, from the data in **(C)** and **(E)**. ND represents not determined.

To assess the ability of PNKP to recognize non-ligatable nicks in the nucleosome, we utilized electrophoretic mobility shift assays (EMSAs) to determine apparent binding affinities (*K*_d,app_) of PNKP for a nucleosome without DNA damage (ND-NCP) and the five non-ligatable Nick-NCPs (Fig. 1C and SI Appendix, Fig. S2). Importantly, these experiments were performed using a PNKP D171A phosphatase dead mutant^48,50,53^ in the absence of Mg^2+^ and ATP to prevent catalysis. To establish a baseline, we initially performed the EMSAs with PNKP and a nucleosome without DNA damage (ND-NCP). This EMSA analysis revealed PNKP binds the ND-NCP with a *K*_d,app_ of 136 ± 7 nM (Fig. 1C and SI Appendix, Fig. S2), consistent with robust nucleosome binding even in the absence of a non-ligatable nick. We then performed EMSAs with the five Nick-NCPs to determine whether PNKP engages non-ligatable nicks at different translational positions throughout the nucleosome. PNKP bound Nick-NCP−5, Nick-NCP−4, Nick-NCP−3, Nick-NCP−2, and Nick-NCP−1 with apparent binding affinities of 63 ± 20 nM, 94 ± 21 nM, 59 ± 5 nM, 74 ± 4 nM, and 96 ± 11 nM, respectively (Fig. 1C and SI Appendix, Fig. S2). The apparent binding affinities for the Nick-NCPs are 1.4- to 2.4-fold tighter than that observed for the ND-NCP, indicating that PNKP has a modest specificity for non-ligatable nicks in the nucleosome. Of note, PNKP bound all five Nick-NCPs with similar apparent affinities, indicating that PNKP can also readily bind non-ligatable nicks at each translational position in the nucleosome.

To define how efficiently PNKP processes non-ligatable nicks in the nucleosome, we determined initial reaction velocities for PNKP on the five non-ligatable Nick-NCPs under steady-state conditions. To overcome challenges associated with the simultaneous detection of the phosphatase and kinase activities, we developed a ligase-coupled product formation assay for combined detection of the PNKP phosphatase and kinase activities in the nucleosome (Fig. 1D, see methods). As a control, we first measured the initial velocity of PNKP on a 147 bp DNA containing a non-ligatable nick (Nick-DNA). PNKP rapidly converted the non-ligatable Nick-DNA into a ligatable Nick-DNA with a *v*_0_ of 0.207 ± 0.005 nM s^−1^(Fig. 1 E and F, and SI Appendix, Fig. S3), consistent with prior observations of robust phosphatase and kinase activities^45–47,49–51^. To determine how the nucleosome impacts PNKP enzymatic activity, we then performed the enzyme kinetics on the five non-ligatable Nick-NCPs. PNKP converted the non-ligatable Nick-NCP into a ligatable Nick-NCP with a *v*_0_ of 0.436 ± 0.081 nM s^−1^ for Nick-NCP−5, 0.026 ± 0.004 nM s^−1^ for Nick-NCP−4, 0.119 ± 0.002 nM s^−1^ for Nick-NCP−3, 0.131 ± 0.012 nM s^−1^ for Nick-NCP−2, and 0.023 ± 0.005 nM s^−1^ for Nick-NCP−1 (Fig. 1 E and F, and SI Appendix, Fig. S3). The initial velocities of the reaction at SHL−5, SHL−3 and SHL−2 were 2.1-fold faster, 1.7-fold slower, and 1.6-fold slower relative to the non-ligatable Nick-DNA, respectively, indicating that PNKP can catalyze end processing in the nucleosome at these locations with comparable rates to the non-ligatable Nick-DNA. In contrast, the initial velocities of the reaction at SHL−4 and SHL−1 were 8-fold and 9-fold slower relative to the non-ligatable Nick-DNA. Together, the enzyme kinetics indicate that PNKP can process non-ligatable nicks throughout the nucleosome, though the efficiency of non-ligatable nick processing has a subtle dependence on the translational position of the solvent-exposed non-ligatable nick in the nucleosome. Importantly, the position-dependent differences in PNKP end processing in the nucleosome are substantially smaller than that observed for downstream SSBR enzymes DNA Polymerase β and DNA Ligase IIIα^42,43,56–58^.

### Structural basis of non-ligatable nick recognition in the nucleosome by PNKP

To understand the structural mechanism used by PNKP to recognize and process a non-ligatable nick in the nucleosome, we generated a pre-catalytic PNKP-Nick-NCP−3 complex stabilized by mild glutaraldehyde crosslinking and subjected the complex to single particle analysis (SI Appendix, Fig. S4). Importantly, the complex was generated using a PNKP D171A phosphatase dead mutant^48,50,53^ in the absence of Mg^2+^ and ATP to ensure PNKP was captured in a pre-catalytic conformation. The cryo-EM dataset resulted in a 2.6 Å reconstruction of the Nick-NCP−3 in the absence of PNKP and a 2.9 Å reconstruction of the PNKP-Nick-NCP−3 complex with PNKP in a pre-catalytic conformation poised for 3′-phosphate hydrolysis and 5′-OH phosphorylation (Fig. 2A and SI Appendix, Figs. S5 and S6, and Table S1), referred to as the PNKP-Nick-NCP−3 (catalytic) complex. The local resolution of PNKP in the PNKP-Nick-NCP−3 (catalytic) structure was 3 - 5 Å (SI Appendix, Fig. S6), which was sufficient to rigid-body dock a previously determined high-resolution X-ray crystal structure of murine PNKP (PDB: 3ZVN)^49^, and manually build protein side chains throughout most of the DNA-binding interface and the PNKP active sites.

**Fig. 2:**
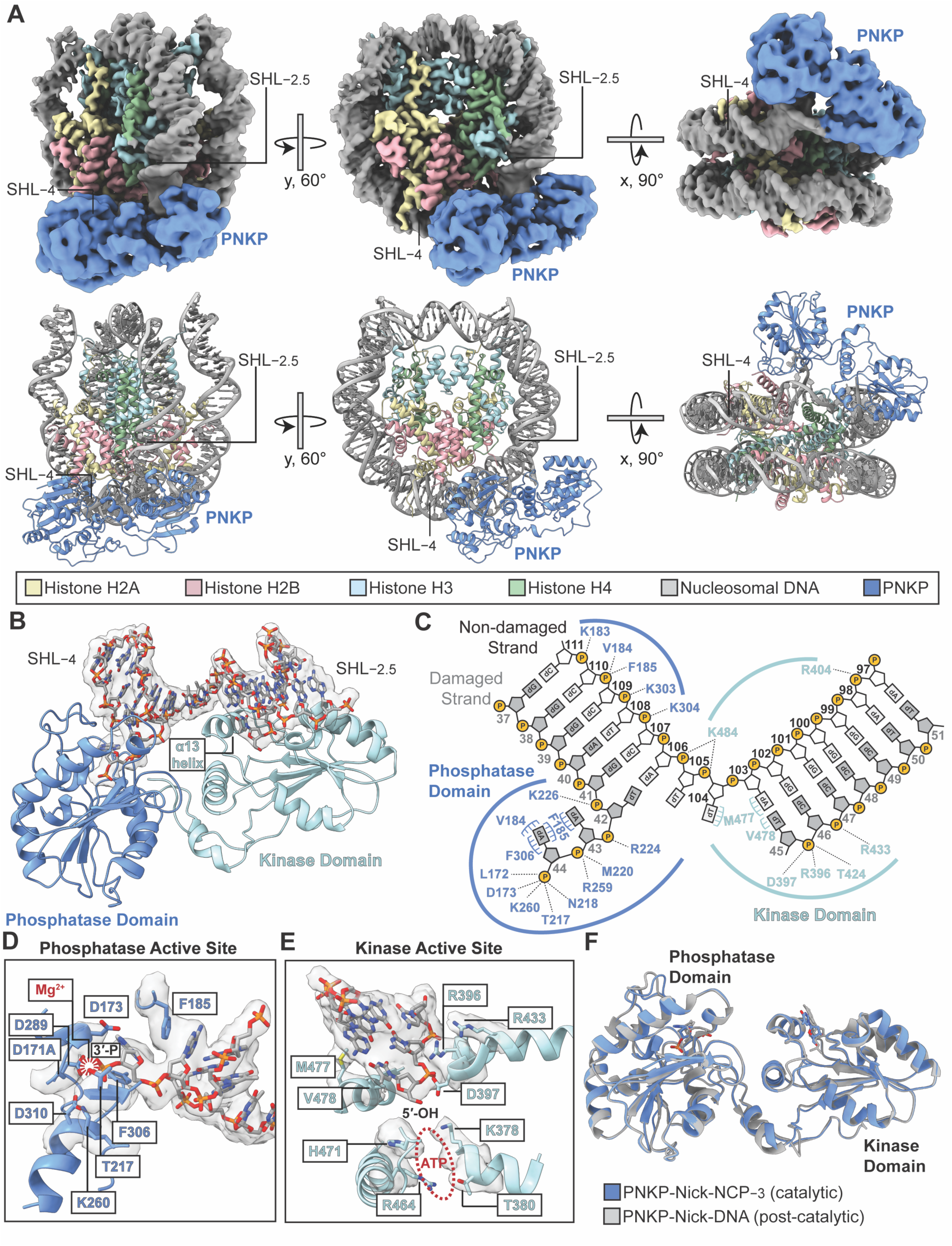
Structural basis of non-ligatable nick recognition in the nucleosome by PNKP. **(A)** Composite cryo-EM map and model of the PNKP-Nick-NCP−3 (catalytic) complex shown in three different orientations. **(B)** Focused view of the PNKP-Nick-NCP−3 (catalytic) complex highlighting the PNKP-nucleosomal DNA binding interface from SHL−2.5 to SHL−4. Segmented cryo-EM density for the nucleosomal DNA is shown as a transparent gray surface. The PNKP phosphatase domain and kinase domain are shown in blue and light blue, respectively. **(C)** Contact map from the PNKP-Nick-NCP−3 (catalytic) complex highlighting the interaction between PNKP and the nucleosomal DNA from SHL−2.5 to SHL−4. The PNKP phosphatase and kinase domain residues that interact with the nucleosomal DNA are labeled in blue and light blue, respectively. **(D-E)** Focused view of the PNKP phosphatase **(D)** and kinase active sites **(E)**. PNKP active site residues are shown as sticks. The segmented cryo-EM density for the nucleosomal DNA and PNKP active site residues is shown as a transparent gray surface. **(F)** Structural comparison of the PNKP-Nick-NCP−3 (catalytic) complex and the post-catalytic PNKP-Nick-DNA complex (PDB: 3ZVN). The PNKP-Nick-NCP−3 (catalytic) complex and post-catalytic PNKP-Nick-DNA complex are shown in blue and gray, respectively. For clarity, only the 3′- and 5′-termini are shown from both structures.

The PNKP-Nick-NCP−3 (catalytic) structure provides structural insight into non-ligatable nick recognition, 3′-phosphate hydrolysis, and 5′-OH phosphorylation by PNKP. During non-ligatable nick recognition, PNKP directly engages the nucleosomal DNA with a core footprint spanning ∼10 bp from SHL−3 to SHL−4 (Fig. 2 A–C). The PNKP-nucleosome binding interface buries ∼ 1,400 Å^2^ surrounding the non-ligatable nick and the interaction is mediated by an extensive contact network between the phosphatase and kinase domains and both non-damaged and damaged strands of the nucleosomal DNA (Fig. 2 B and C). The interaction with the non-damaged strand is mediated by backbone electrostatic contacts involving residues K183, V184, F185, K303, and K304 in the phosphatase domain, and residues R404 and K484 in the kinase domain (Fig. 2C). On the 3′-end of the damaged strand, the α13 helix separates the double-stranded DNA (Fig. 2B), which disrupts the base-pairing interactions between the terminal two nucleotides and repositions the 3′-phosphate into the phosphatase active site. The two unpaired bases on the 3′ end of the nick are stabilized by hydrophobic and aromatic stacking interactions mediated by V184, F185, and F306 (Fig. 2C). Within the phosphatase active site, the 3′-phosphate is stabilized by L172, T217, N218, and K260, and is positioned near the phosphatase catalytic residues D171A, D173 and D289 (Fig. 2D). Importantly, these catalytic residues are critical for nucleophilic attack on the 3′-phosphate, stabilization of the phosphoaspartate intermediate, and 3′-phosphate hydrolysis^49,50,59^. On the 5′-end of the damaged strand, the 5′-OH is repositioned into the kinase active site and stabilized through electrostatic and hydrophobic interactions mediated by R396, D397, T424, M477 and V478 (Fig. 2 C and E). Though ATP and Mg^2+^ are not present in the active site, the 5′-OH is in close proximity to catalytic residue D397 (Fig. 2E), which is essential for deprotonation of the 5′-OH and its subsequent nucleophilic attack on ATP^48,49^. Although we do not observe clear density for the D397 side chain, the adjacent R396 is clearly resolved and in position to stabilize the conformation of D397 for catalysis (Fig. 2E). Ultimately, the overall conformation of the phosphatase and kinase active sites in the PNKP-Nick-NCP−3 structure closely resemble those observed in the previously determined post-catalytic structure of PNKP bound to a pseudo Nick-DNA (Fig. 2F, PDB: 3ZVN)^49^, which strongly suggests that PNKP is poised for catalyzing 3′-phosphate hydrolysis and 5′-OH phosphorylation in the nucleosome.

To adopt the pre-catalytic conformation, PNKP deforms the nucleosomal DNA to simultaneously engage the 3′-phosphate and 5′-OH termini (Fig. 3 A –C). Structural comparison of the PNKP-Nick-NCP−3 (catalytic) and Nick-NCP−3 revealed the 3′-phosphate and 5′-OH termini undergo substantial ∼26 Å and ∼9 Å movements during non-ligatable nick recognition (Fig. 3C), which repositions the 3′-phosphate and 5′-OH into the phosphatase and kinase active sites, respectively (Fig. 2 D and E). The uncoupling and repositioning of the 3′-phosphate and 5′-OH is mediated by the insertion of the α13 helix of the kinase domain into the path of the duplex DNA at the site of the non-ligatable nick, which acts to simultaneously support the exposed base-pair surfaces on both sides of the nick and blocks base stacking across the strand discontinuity (Fig. 3C). On the 3′-phosphate side of the non-ligatable nick, insertion of the α13 helix displaces the terminal two nucleotides and disrupts their base pairing. The displacement of the terminal two nucleotides is accompanied by a ∼70° rotation that directs the 3′-phosphate away from the histone octamer and into the narrow phosphatase active site (Fig. 3 B and C). Importantly, disrupting the base pairing of the terminal two nucleotides is critical for dephosphorylation of the 3′-phosphate, as the phosphatase active site is only large enough to accommodate single-stranded DNA^48–50^. On the 5′-OH side of the non-ligatable nick, insertion of the α13 helix redirects the 5′-OH away from the histone octamer and into the kinase active site (Fig. 3 B and C). Unlike the 3′-end of the non-ligatable nick, the 5′-end remains base paired as the kinase active site is large enough to accommodate double-stranded DNA. Together, the combined deformations required to position the ends of the non-ligatable nick into the PNKP active sites results in bending of the nucleosomal DNA and a subtle displacement of the nucleosomal DNA away from the histone octamer with minimal disruption of histone-DNA contacts (Fig. 3B). The ability of PNKP to readily deform the nucleosomal DNA and engage the non-ligatable nick with minimal disruption of histone-DNA contacts likely explains why PNKP can efficiently process solvent-exposed non-ligatable nicks in the nucleosome (Fig. 1E).

**Fig. 3:**
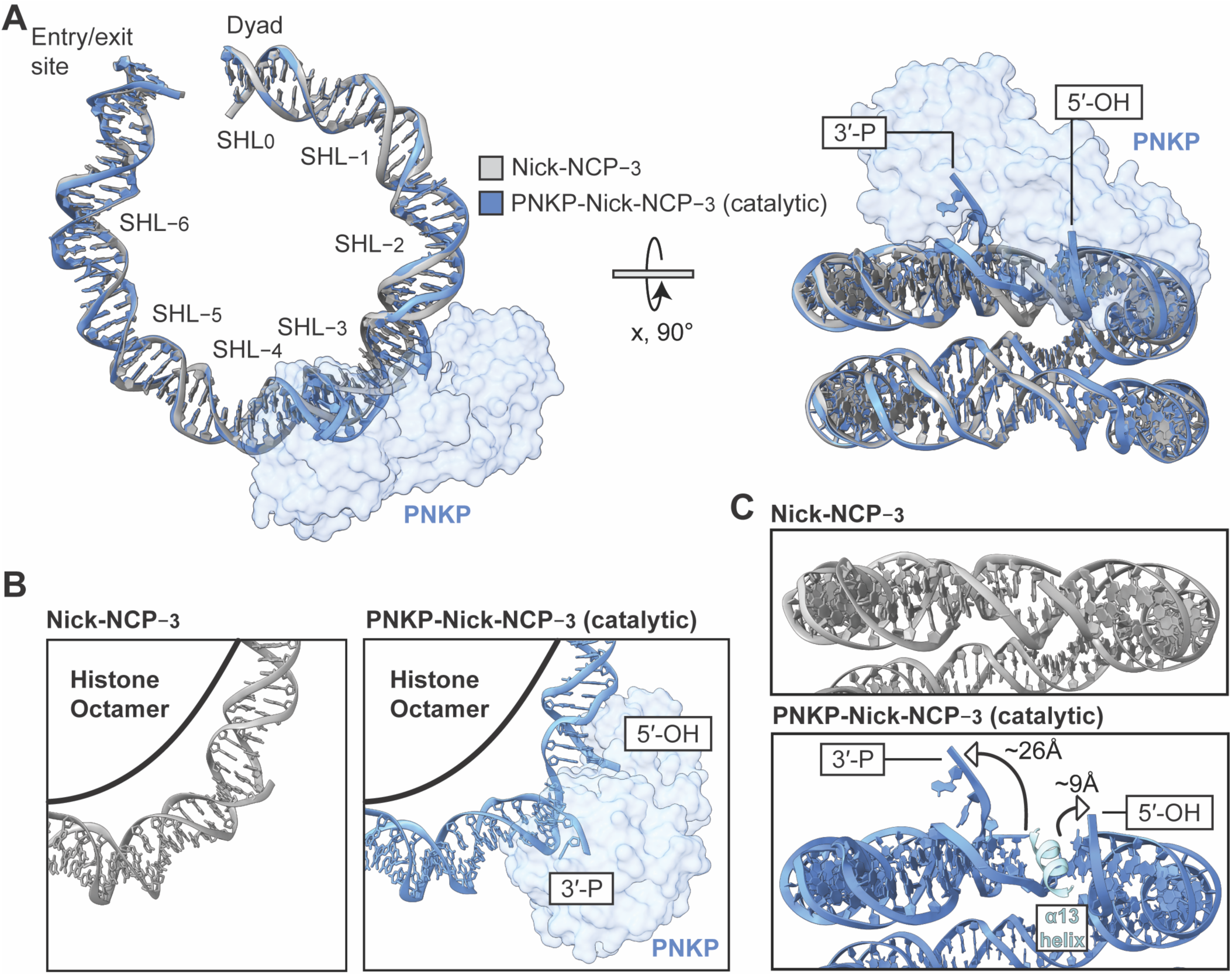
PNKP deforms the nucleosomal DNA during non-ligatable nick recognition. **(A)** Structural comparison of the nucleosomal DNA in the Nick-NCP−3 and the PNKP-Nick-NCP−3 (catalytic) complex shown in two different orientations. The Nick-NCP−3 and PNKP-Nick-NCP−3 (catalytic) complex are shown in gray and blue, respectively. PNKP is shown as a transparent blue surface for improved clarity. **(B)** Focused views of the nucleosomal DNA from SHL−2 to SHL−4.5 in the Nick-NCP−3 (left) and PNKP-Nick-NCP−3 (catalytic) complex (right). **(C)** Focused views of the nucleosomal DNA from the inter-gyres perspective in the Nick-NCP−3 (top) and PNKP-Nick-NCP−3 (catalytic) complex (bottom). The α13 helix of the PNKP kinase domain is shown as a cartoon in light blue.

### PNKP uses a conserved mechanism for non-ligatable nick recognition throughout the nucleosome

To determine whether PNKP uses unique mechanisms to recognize and processes non-ligatable nicks at different translational positions in the nucleosome, we generated pre-catalytic PNKP-Nick-NCP−4 and PNKP-Nick-NCP−5 complexes stabilized by mild glutaraldehyde crosslinking and subjected the complexes to single particle analysis (SI Appendix, Figs. S7 and S8). Similar to the PNKP-Nick-NCP−3 complex, these complexes were generated using the PNKP D171A phosphatase dead mutant^48,50,53^ in the absence of Mg^2+^ and ATP to ensure PNKP was captured in a pre-catalytic conformation. The cryo-EM datasets resulted in a 3.5 Å and 3.2 Å reconstruction of the PNKP-Nick-NCP−4 and PNKP-Nick-NCP−5 complexes (Fig. 4 A and B, and SI Appendix, Figs. S10 and S12), respectively, with PNKP in a pre-catalytic conformation poised for 3′-phosphate hydrolysis and 5′-OH phosphorylation. These structures are referred to as the PNKP-Nick-NCP−4 (catalytic) and PNKP-Nick-NCP−5 (catalytic) complexes. In addition, we also obtained a 2.6 Å and 2.8 Å reconstruction of Nick-NCP−4 and Nick-NCP−5, respectively (SI Appendix, Figs. S9 and S11). Of note, the local resolution of PNKP in the PNKP-Nick-NCP−4 (catalytic) and PNKP-Nick-NCP−5 (catalytic) reconstructions ranged from 5 - 7 Å, which was sufficient to observe general secondary structural features and unambiguously dock the PNKP catalytic domains. However, the local resolution of PNKP was not sufficient to readily model side chains conformations at the DNA binding interface or within the PNKP active site (SI Appendix, Figs. S10 and S12). Therefore, the PNKP side chains within these structures generally reflect their positions in the higher-resolution PNKP-Nick-NCP−3 structure.

**Fig. 4:**
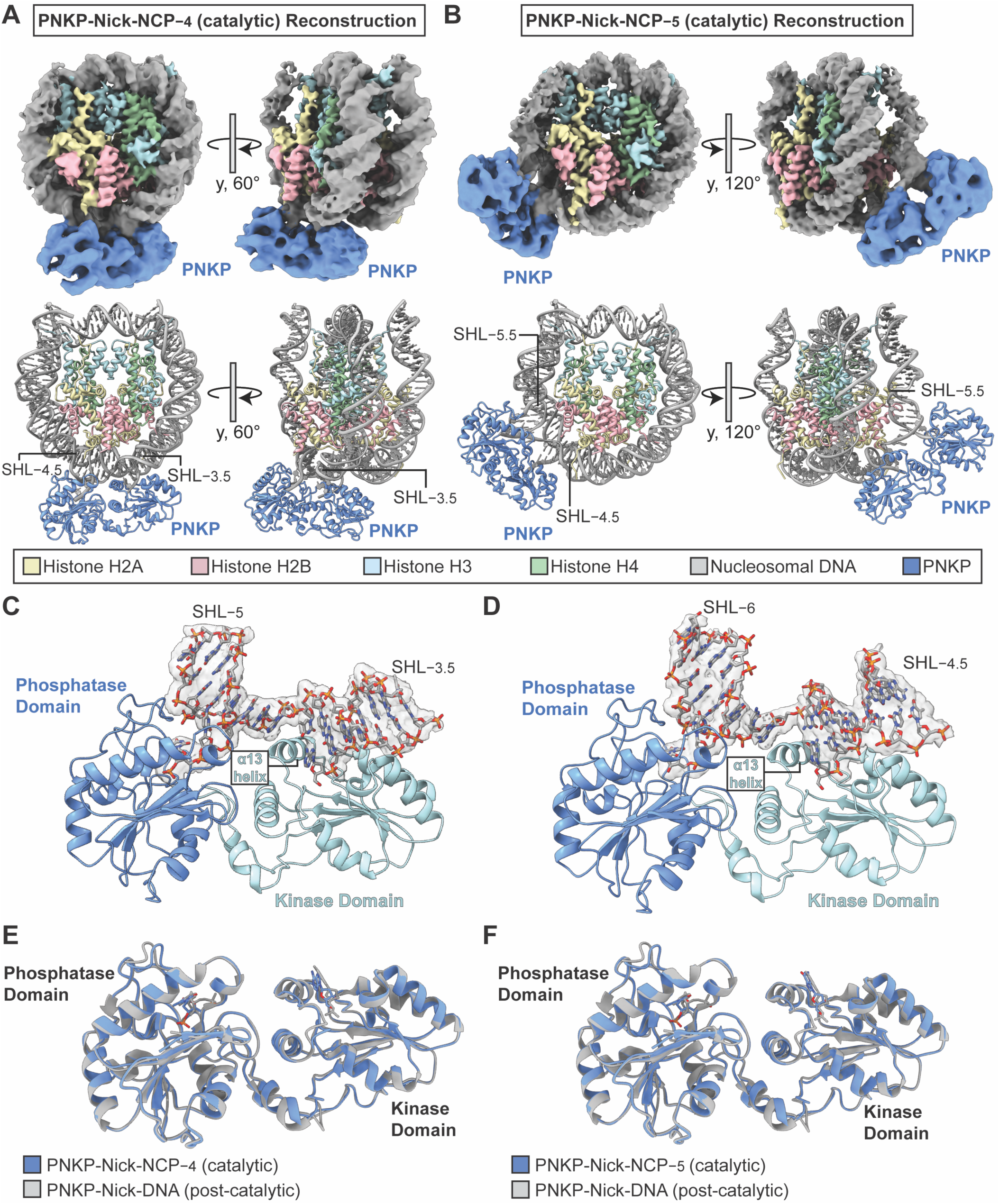
A conserved mechanism for non-ligatable nick recognition throughout the nucleosome. **(A)** Composite cryo-EM map and model of the PNKP-Nick-NCP−4 (catalytic) complex shown in two different orientations. **(B)** Composite cryo-EM map and model of the PNKP-Nick-NCP−5 (catalytic) complex shown in two different orientations. **(C)** Focused view of the PNKP-Nick-NCP−4 (catalytic) complex highlighting the PNKP-nucleosomal DNA binding interface from SHL−3.5 to SHL−5. **(D)** Focused view of the PNKP-Nick-NCP−5 (catalytic) complex highlighting the PNKP-nucleosomal DNA binding interface from SHL−4.5 to SHL−6. In **C** and **D**, the segmented cryo-EM density for the nucleosomal DNA is shown as a transparent gray surface and the PNKP phosphatase domain and kinase domain are shown in blue and light blue, respectively. **(E)** Structural comparison of the pre-catalytic PNKP-Nick-NCP−4 complex and the post-catalytic PNKP-Nick-DNA complex (PDB: 3ZVN) **(F)** Structural comparison of the pre-catalytic PNKP-Nick-NCP−5 complex and the post-catalytic PNKP-Nick-DNA complex (PDB: 3ZVN). In **E** and **F**, the PNKP-Nick-NCP (catalytic) complexes and post-catalytic PNKP-Nick-DNA complex are shown in blue and gray, respectively. For clarity, only the 3′- and 5′-termini are shown from both structures.

The PNKP-Nick-NCP−4 (catalytic) and PNKP-Nick-NCP−5 (catalytic) structures reveal that PNKP engages the non-ligatable nick using a mechanism strikingly similar to that observed for PNKP at SHL −3 (Fig. 4 A and B). During non-ligatable nick recognition at SHL−4 and SHL−5, PNKP directly engages ∼10 bp of nucleosomal DNA surrounding the non-ligatable nick, with the phosphatase and kinase domains forming extensive interactions with the non-damaged and damaged strands (Fig. 4 C and D). In both structures, the phosphatase domain engages the terminal 3′-phosphate and the kinase domain engages the terminal 5′-OH terminus, positioning the two ends of the non-ligatable nick within the two active sites (Fig. 4 C and D). The ability of PNKP to reposition the 3′-phosphate and 5′-OH into the phosphatase and kinase active sites is mediated by displacement of the nucleosomal DNA away from the histone octamer at the PNKP binding site with minimal disruption of histone-DNA contacts (SI Appendix, Figs. S13 and S14). Although the local resolution of PNKP precludes the ability to assign the exact conformations of active site side chains in both structures (SI Appendix, Figs. S10 and S12), the phosphatase and kinase domains adopt a similar conformation to that of the post-catalytic structure of PNKP bound to a pseudo Nick-DNA (Fig. 4 E and F, PDB: 3ZVN)^49^, strongly suggesting these structures represent PNKP in a conformation poised for 3′-phosphate hydrolysis and 5′-OH phosphorylation. Further comparison of the PNKP-Nick-NCP−4 and PNKP-Nick-NCP−5 structures with the PNKP-Nick-NCP−3 structure also revealed a similar mode of non-ligatable nick recognition at each translational position in the nucleosome (SI Appendix, Fig. S15), suggesting that PNKP engages and processes solvent-exposed non-ligatable nicks throughout the nucleosome using a similar general mechanism. Ultimately, the ability of PNKP to readily adopt a catalytic conformation at SHL−3, SHL−4, and SHL−5 without substantial disruptions to overall nucleosome structure provides a strong rationale for the robust end processing observed at multiple solvent-exposed translational positions throughout the nucleosome (Fig. 1E).

### The PNKP FHA domain engages the nucleosome acidic patch during end processing

During 3D-classification of the PNKP-Nick-NCP−3, PNKP-Nick-NCP−4, and PNKP-Nick-NCP−5 cryo-EM datasets, we identified two distinct classes of PNKP-bound particles containing additional density adjacent to the nucleosome acidic patch (Fig. 5 A–C and SI Appendix, Figs. S4, S7, and S8). Importantly, the density adjacent to the acidic patch matches the general dimensions and secondary structural features of the PNKP FHA domain (Fig. 5 A–C and SI Appendix, Figs. S16–S21). Additional refinement of the first unique class resulted in 3.1 Å, 3.2 Å, and 3.4 Å reconstructions of the PNKP-Nick-NCP complex with the FHA domain bound to the nucleosome acidic patch and the catalytic domains bound to the non-ligatable nick at SHL−3, SHL−4, and SHL−5, respectively (Fig. 5 A–C, SI Appendix, Figs. S16–S18, and Tables S1–S3). We refer to these structures as the PNKP-Nick-NCP−3 (catalytic-FHA), PNKP-Nick-NCP−4 (catalytic-FHA), and PNKP-Nick-NCP−5 (catalytic-FHA) complexes. Additional refinement of the second unique class resulted in 3.1 Å, 3.1 Å, and 3.0 Å reconstructions of the PNKP-Nick-NCP complex with the FHA domain bound to the acidic patch and the catalytic domains of PNKP disengaged from the non-ligatable nick at SHL−3, SHL−4, and SHL−5, respectively (SI Appendix, Figs. S18–S21 and Tables S1–S3). We refer to these structures as the PNKP-Nick-NCP−3 (FHA), PNKP-Nick-NCP−4 (FHA), and PNKP-Nick-NCP−5 (FHA) complexes, respectively. In all six reconstructions, the local resolution of the PNKP FHA domain ranged from 5 - 9 Å (SI Appendix, Figs. S16–S21), which was sufficient to dock a previously determined high-resolution X-ray crystal structure of the human PNKP FHA domain (PDB: 2W3O)^34^, but not sufficient to model FHA side chain residues that mediate the acidic patch interaction.

**Fig. 5:**
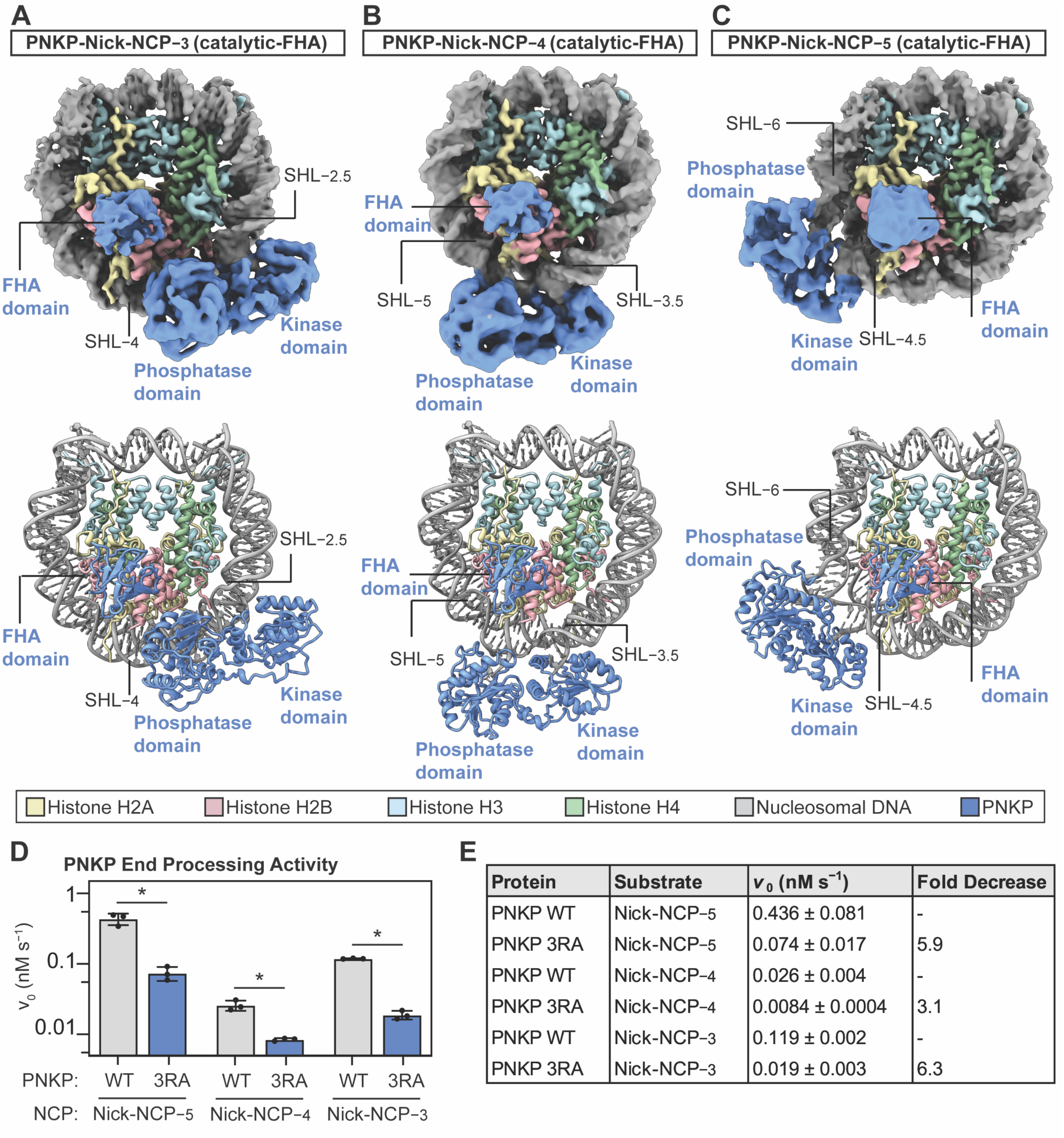
The PNKP FHA domain engages the nucleosome acidic patch during end processing. **(A)** Composite cryo-EM map and model of the PNKP-Nick-NCP−3 (catalytic-FHA) complex**. (B)** Composite cryo-EM map and model of the PNKP-Nick-NCP−4 (catalytic-FHA) complex**. (C)** Composite cryo-EM map and model of the PNKP-Nick-NCP−5 (catalytic-FHA) complex**. (D)** Quantification of the initial velocities (*v*_0_) obtained for PNKP WT (gray) and PNKP 3RA (blue) end processing on Nick-NCP−5, Nick-NCP−4, and Nick-NCP−3. The data points represent the mean ± standard deviation from three independent replicate experiments, and the p-values were obtained using a two-tailed Student’s T-test (* p < 0.01) (see SI Appendix, Fig. S26 for associated data). The PNKP WT data (gray bars) presented in **(D)** are the same data as presented in Fig. 1E. Summary of the initial velocities (v_0_) and the fold decrease in end processing for PNKP 3RA relative to PNKP WT on Nick-NCP−5, Nick-NCP−4, and Nick-NCP−3, from the data in **(D)** and Fig. 1E.

In the PNKP-Nick-NCP (catalytic-FHA) and PNKP-Nick-NCP (FHA) structures at SHL−3, SHL−4, and SHL−5, the PNKP FHA domain directly engages the nucleosome acidic patch (Fig. 5 A–C and SI Appendix, Fig. S22). Though the modest resolution of the FHA domain precludes accurate assignment of side-chain residues that mediate this interaction (SI Appendix, Figs. S16–S21), the general conformation observed for the FHA domain orients two unstructured loops containing arginine residues R35, R44, and R48 towards the nucleosome acidic patch (SI Appendix, Fig. S23). Importantly, the arginine residues in the PNKP FHA domain are analogous to the arginine anchors used by many chromatin-associated proteins to engage the nucleosome acidic patch^60^. Further structural comparison of the PNKP-Nick-NCP (catalytic-FHA) and PNKP-Nick-NCP (FHA) structures at SHL−3, SHL−4, and SHL−5 revealed a similar mode of nucleosome acidic patch binding by the FHA domain regardless of whether the catalytic domains were engaged with the non-ligatable nick (SI Appendix, Fig. S24). This indicates that the PNKP FHA domain can readily engage the nucleosome acidic patch independent of non-ligatable nick recognition by the catalytic domains, and that non-ligatable nick recognition by the catalytic domains has minimal impact on the mode of acidic patch binding by the FHA domain.

To determine whether nucleosome acidic patch binding alters the mechanism used by PNKP for non-ligatable nick recognition, we compared the structures of the PNKP-Nick-NCP (catalytic) and the PNKP-Nick-NCP (catalytic-FHA) complexes at SHL−3, SHL−4, and SHL−5 (SI Appendix, Fig. S25). This structural analysis revealed a similar mechanism for non-ligatable nick recognition in the PNKP-Nick-NCP (catalytic) and PNKP-Nick-NCP (catalytic-FHA) structures, suggesting that PNKP can readily adopt a catalytically active conformation at SHL−3, SHL−4, and SHL−5 while simultaneously engaged with the nucleosome acidic patch. To determine if the interaction with the nucleosome acidic patch is important for PNKP to efficiently process non-ligatable nicks in the nucleosome, we generated a PNKP R35A/R44A/R48A mutant (PNKP 3RA) and tested the impact of this mutant on the ability of PNKP to convert a non-ligatable nick to a ligatable nick in the nucleosome at SHL−5, SHL−4, and SHL−3 under identical steady-state conditions as PNKP WT. These experiments revealed PNKP 3RA catalyzes end processing with a *v*_0_ of 0.074 ± 0.017 nM s^−1^ for Nick-NCP−5, 0.0084 ± 0.0004 nM s^−1^ for Nick-NCP−4, and 0.019 ± 0.003 nM s^−1^ for Nick-NCP−3, which are 5.9-fold, 3.1-fold, and 6.3-fold slower compared to WT PNKP, respectively (Fig. 5 D and E, and SI Appendix, Fig. S26). This suggests that the interaction of the FHA domain with the nucleosome acidic patch accelerates the ability of PNKP to catalyze end processing in the nucleosome.

## Discussion

The enzymatic processing of non-ligatable nicks with chemically heterogeneous DNA ends is a critical step for initiating SSBR. How end-processing enzymes process these chemically heterogeneous DNA ends to initiate SSBR within the context of chromatin has been largely unexplored. Our work has provided mechanistic details into how PNKP processes non-ligatable nicks in the nucleosome. Our biochemical assays revealed that PNKP can readily engage solvent exposed non-ligatable nicks at multiple translational positions in the nucleosome and rapidly convert them to ligatable nicks. The cryo-EM structures of PNKP engaged with a non-ligatable nick at SHL−3, SHL−4, and SHL−5 provide a structural rationale for this robust activity in the nucleosome. These structures identified that PNKP displaces the nucleosomal DNA from the histone octamer and bends the nucleosomal DNA during non-ligatable nick recognition. Ultimately, these deformations in the nucleosomal DNA reposition 3′-phosphate and 5′-OH into the phosphatase and kinase active sites for catalysis, with minimal disruption of histone-DNA contacts and overall nucleosome structure. The ability of PNKP to adopt a catalytic conformation without clashing with the histone octamer likely explains the ability of PNKP to efficiently process solvent-exposed non-ligatable nicks throughout the nucleosome. Of note, PNKP also processes non-ligatable nicks and gaps containing an individual 3′-phosphate or 5′-OH^45–51^. While our biochemical and structural analysis focused exclusively on non-ligatable nicks containing a 3′-phosphate and 5′-OH, we hypothesize that PNKP likely employs a similar strategy to engage and process non-ligatable nicks and gaps containing an individual 3′-phosphate or 5′-OH^49,50^.

In addition to defining how PNKP engages non-ligatable nicks in the nucleosome, our structural analysis revealed a second binding interface between PNKP and the nucleosome. This second binding interface is mediated by the PNKP FHA domain, which interacts with the nucleosome acidic patch through two loops containing a series of arginine residues (R35, R44, and R48). Though the interaction of the FHA domain with the nucleosome acidic patch is clearly not required for non-ligatable nick recognition by the phosphatase and kinase domains, our biochemical assays with a PNKP R35A/R44A/R48A mutant indicate this interaction can accelerate end processing in the nucleosome. Notably, a PNKP R35A/R48A mutant was previously shown to severely impair the accumulation of PNKP at sites of cellular DNA damage and delay SSBR^61,62^. However, interpreting the importance of the nucleosome acidic patch interaction from these experiments is challenging, as these two arginine residues within the PNKP FHA domain are also critical for the interaction with phosphorylated XRCC1^34,48,63^ and the phosphorylated XRCC1-dependent accumulation of PNKP at cellular SSBs^62,64–66^. Consequently, the overlapping FHA binding site would suggest that PNKP likely forms a mutually exclusive interaction with phosphorylated XRCC1 or the nucleosome acidic patch. We hypothesize that nucleosome acidic patch binding may be important for stabilizing PNKP on the nucleosome following the phosphorylated XRCC1-dependent recruitment of PNKP to cellular SSBs. Tethering PNKP to the nucleosome acidic patch within this context may support the phosphatase and kinase domains as they attempt to engage the non-ligatable nick, deform the nucleosomal DNA, and reposition the non-ligatable nick within the active sites for catalysis. This intriguing model would not require the complete dissociation of PNKP from XRCC1, as XRCC1 also interacts with PNKP via a secondary, low-affinity interaction with the catalytic domains^67–69^. Finally, nucleosome acidic patch binding may also contribute to the XRCC1-independent recruitment of PNKP to SSBs previously described^61,65,70,71^. Future work will be needed to dissect the functional importance of nucleosome acidic patch binding by PNKP during chromatin-based SSBR.

SSBR is a multi-step DNA repair pathway with three major enzymatic steps including initial end processing, gap-filling DNA synthesis, and/or nick ligation. Prior work established that DNA Polymerase β and DNA Ligase IIIα can process solvent-exposed DNA damage in the nucleosome^42,43,56–58^. However, the enzymatic efficiencies of DNA Polymerase β and DNA Ligase IIIα are highly dependent on the translational position of the DNA damage in the nucleosome, where solvent-exposed gaps and nicks outside of the nucleosome entry/exit site are refractory towards repair^42,43,56–58^. In contrast to DNA Polymerase β and DNA Ligase IIIα, our biochemical assays indicate that PNKP can readily process non-ligatable nicks at solvent-exposed positions throughout the nucleosome, with the translational position of the non-ligatable nick only subtly impacting the enzymatic activity of PNKP. This suggests that the nucleosome differentially impacts the three enzymatic steps of chromatin-based SSBR, and that gap-filling DNA synthesis and nick ligation are more sensitive to nucleosome structure. The differential impact of nucleosome structure on SSBR likely stems from intrinsic differences in the enzyme-induced conformational changes in the nucleosomal DNA required for each enzyme to adopt a catalytic conformation during SSB recognition. In contrast to PNKP, DNA Polymerase β and DNA Ligase IIIα require substantially larger deformations in the nucleosomal DNA to position the 1-nt gap and nick within their respective active sites^42,43,72^. Ultimately, the impaired activities of DNA Polymerase β and DNA Ligase IIIα at many solvent-exposed positions in the nucleosome implies the terminal enzymatic steps of SSBR may frequently require active chromatin remodeling for efficient repair. This active chromatin remodeling may be mediated by Poly-(ADP-ribose) Polymerases (PARPs), which are critical for efficient SSBR^25–29^, and/or downstream ATP-dependent chromatin remodeling enzymes that have been implicated in SSBR^73–81^. However, future work will be needed to dissect the impact of PARPs and the various downstream ATP-dependent chromatin remodeling enzymes on chromatin-based SSBR.

## Methods

### Purification of recombinant human PNKP

Codon optimized *H. sapiens* full-length (FL) PNKP WT, PNKP D171A, and PNKP R35A/R44A/R48A (3RA) in a pET28a vector were purchased from GenScript. All PNKP proteins were expressed and purified using the same purification scheme. In brief, the pET28a-PNKP WT and mutant plasmids were transformed and expressed in BL21(DE3) RIPL *E. coli* cells (Agilent). The transformed cells were grown at 37 °C to an OD_600_ of 1.0, and induced with 0.5 mM IPTG for 20 – 24 hours at 25 °C. The cells were subsequently harvested and resuspended in a buffer containing 25 mM HEPES (pH 8.0), 250 mM NaCl, 0.25 mM TCEP, 10 mM Imidazole, and a protease inhibitor cocktail (AEBSF, leupeptin, benzamidine, pepstatin A). The cells were then lysed via sonication, and the cell lysate clarified via centrifugation. The supernatant containing PNKP protein was purified under gravity flow using a HisPur Ni-NTA resin (Thermo Fisher Scientific) equilibrated with a buffer containing 25 mM HEPES (pH 8.0), 250 mM NaCl, 0.25 mM TCEP, 10 mM Imidazole, and eluted with a buffer containing 25 mM HEPES (pH 8.0), 250 mM NaCl, 0.25 mM TCEP, 400 mM Imidazole. The PNKP protein was further purified by cation-exchange chromatography using a HiTrap SP HP (Cytiva) equilibrated in a buffer containing 25 mM HEPES (pH 7.5), 200 mM NaCl, and 0.1 mM TCEP, and eluted from the column in a buffer containing 25 mM HEPES (pH 7.5), 1 M NaCl, and 0.1 mM TCEP (0 – 100% gradient). The PNKP protein was then polished by size-exclusion chromatography using a HiPrep 26/60 Sephacryl S-200 HR (Cytiva) equilibrated in a buffer containing 25 mM HEPES (pH 7.5), 150 mM NaCl, and 0.1 mM TCEP. The purity of the PNKP proteins was confirmed by SDS-PAGE and the purified PNKP proteins stored long term at −80 °C. An SDS-PAGE gel of the purified PNKP proteins can be found in SI Appendix, Fig. S1C. Of note, the PNKP WT and mutant proteins contain a C-terminal 6xHis tag that was not removed during the purification.

### Preparation of oligonucleotides

The oligonucleotides used to generate recombinant nucleosomes containing nicks with 3′-phosphate and 5′-OH were synthesized by Integrated DNA Technologies (Coralville, IA). Each individual oligonucleotide was resuspended in a buffer containing 10 mM Tris (pH 7.5) and 1 mM EDTA to a final concentration of 100 µM. Equimolar amounts of complementary oligonucleotides (SI Appendix, Tables S4 and S5) were mixed and diluted with a buffer containing 10 mM Tris (pH 7.5) and 1 mM EDTA to a final concentration of 10 µM. The complementary oligonucleotides were then annealed by heating to 95°C for 2 minutes and cooling to 10 °C at a rate of −5 °C/min. The annealed oligonucleotides were generated immediately prior to nucleosome reconstitution. A complete list of the oligonucleotides used to generate recombinant nucleosomes containing non-ligatable nicks with a 3′-phosphate and 5′-OH can be found in (SI Appendix, Tables S4 and S5).

### Purification of recombinant human histones

Recombinant *H. sapiens* histone H2A (UniProt identifier: P0C0S8), histone H2B (UniProt identifier: P62807), histone H3 C110A (UniProt identifier: Q71DI3), and histone H4 (UniProt identifier: P62805) proteins were generated from a pET3a expression vector. The vector containing histone H2A, H3, and H4 were transformed and expressed in T7 Express lysY/I^q^ competent *E. coli* cells (New England BioLabs), and the vector containing histone H2B was transformed and expressed in BL21-CodonPlus (DE3)-RIPL *E. coli* cells (Agilent). All histones were expressed in M9 minimal media supplemented with 0.4% (w/v) glucose. For histone H2A and H2B, the cells were grown at 37 °C to an OD_600_ of 0.4 and expression was induced with 0.4 mM IPTG for 4 hours. For histone H3, the cells were grown at 37 °C to an OD_600_ of 0.4 and expression was induced with 0.4 mM IPTG for 3 hours. For histone H4, the cells were grown at 37 °C to an OD_600_ of 0.4 and expression was induced with 0.3 mM IPTG for 3 hours. The cells were harvested via centrifugation and resuspended in a buffer containing 50 mM Tris (pH 7.5), 100 mM NaCl, 1 mM benzamidine, 1 mM DTT, and 1 mM EDTA. The purification of each individual histone was carried out using established methods^82,83^. The cell pellets containing histones were lysed via sonication in a buffer containing 50 mM Tris (pH 7.5), 100 mM NaCl, 1 mM benzamidine, 1 mM DTT, and 1 mM EDTA, and the cell lysate clarified via centrifugation. The resulting pellet containing histones was then washed two times with a buffer containing 50 mM Tris (pH 7.5), 100 mM NaCl, 1 mM benzamidine, 1 mM DTT, 1 mM EDTA, and 1% Triton X-100, and washed an additional time with a buffer containing 50 mM Tris (pH 7.5), 100 mM NaCl, 1 mM benzamidine, 1 mM DTT, and 1 mM EDTA. The histones were then extracted from the pellet under denaturing condition using a buffer containing 10 mM Tris (pH 7.5), 6 M Guanidinium-HCl, and 10 mM DTT. The extracted histones were then purified using subtractive anion-exchange chromatography and cation-exchange chromatography. The purified histones were then dialyzed five times against H_2_O, lyophilized, and stored long term at −20 °C.

### Generation of H2A/H2B dimer and H3/H4 tetramer

H2A/H2B dimers and H3/H4 tetramers were prepared prior to nucleosome reconstitution using established methods^82,83^. The H2A/H2B dimer was generated by resuspending lyophilized histone H2A and H2B in a buffer containing 20 mM Tris (pH 7.5), 6 M Guanidinium-HCl, and 10 mM DTT. The resuspended histone H2A and H2B were then mixed in a 1:1 molar ratio and dialyzed three times against a buffer containing 20 mM Tris (pH 7.5), 2 M NaCl, and 1 mM EDTA at 4 °C. The H3/H4 tetramer was generated by resuspending lyophilized histone H3 and H4 in a buffer containing 20 mM Tris (pH 7.5), 6 M Guanidinium-HCl, and 10 mM DTT. The resuspended histone H3 and H4 were then mixed in a 1:1 molar ratio and dialyzed three times against a buffer containing 20 mM Tris (pH 7.5), 2 M NaCl, and 1 mM EDTA at 4 °C. The refolded H2A/H2B dimer and H3/H4 tetramer were subsequently purified by size-exclusion chromatography using a HiPrep Sephacryl S-200 16/60 HR equilibrated in a buffer containing 20 mM Tris (pH 7.5), 2 M NaCl, and 1 mM EDTA. The purity of the H2A/H2B dimer and H3/H4 tetramer was confirmed by SDS-PAGE, and the purified H2A/H2B dimer and H3/H4 tetramer stored long term at −20 °C in 50% glycerol.

### Nucleosome reconstitution

All nucleosomes were prepared using an established salt-dialysis method with minor modifications^82,83^. H2A/H2B dimer, H3/H4 tetramer, and annealed Nick-DNA were mixed at a 2.2:1:1 molar ratio, respectively, and the components equilibrated in a buffer containing 20 mM Tris (pH 7.5), 2 M NaCl, and 1 mM EDTA for one hour. The nucleosomes were then reconstituted by sequentially decreasing the concentration of NaCl from 2 M NaCl to 1.5 M NaCl, to 1.0 M NaCl, to 0.75 M NaCl, to 0.5 M NaCl, to 0.25 M NaCl, and to 0 M NaCl over 24 - 28 hours. The reconstituted nucleosomes were then purified by sucrose gradient ultracentrifugation (10–40% gradient). The final nucleosome purity and homogeneity was assessed via native PAGE (5%, 59:1 acrylamide:bis-acrylamide ratio) and the nucleosomes stored in a buffer containing 10 mM Tris (pH 7.5) and 1 mM EDTA at 4 °C. Native PAGE gels of the final nucleosomes can be found in SI Appendix, Figs. S1A and B.

### Electrophoretic mobility shift assays

Electrophoretic mobility shift assay reactions were carried out by mixing each respective NCP (20 nM) with increasing concentrations of PNKP D171A in a buffer containing 50 mM HEPES (pH 7.5), 50 mM NaCl, 1 mM TCEP, 0.1 mg/ml bovine serum albumin, 5% sucrose, and 0.25 % (w/v) bromophenol blue. The reactions were incubated for 10 mins at 4 °C prior to separation of the free NCP and PNKP-NCP species by native polyacrylamide gel electrophoresis (5%, 59:1 acrylamide:bis-acrylamide). The disappearance of the free NCP band was quantified using ImageJ^84^ and fit to a modified Hill equation:

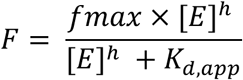

Where *fmax* is the maximal nucleosome bound, [*E*] is the total PNKP concentration, ℎ is the Hill coefficient, and *K_d_*_,*app*_ is the apparent dissociation constant. Each experimental point represents the average of three independent replicate experiments, and the error bars represent the standard deviation of the three independent replicate experiments. The *K*_d,app_ is reported as the mean of the three independent replicate experiments ± the standard deviation of the three independent replicate experiments.

### PNKP enzyme kinetics

PNKP enzyme kinetics assays were carried out under steady-state conditions to extract initial velocities of the reaction. The assays were performed by mixing the Nick-DNA (50 nM) or each individual Nick-NCP (50 nM) with WT PNKP (0.5 nM) or PNKP 3RA (0.5 nM) in a buffer containing 50 mM HEPES (pH 7.5), 100 mM NaCl, 1 mM TCEP, and 0.2 mg/mL BSA. The reactions were incubated at 37 °C for 2 minutes and the reactions initiated through the addition of 2 mM MgCl_2_ and 1 mM ATP. The reactions were quenched at various time points with 10 mM EDTA. The substrate and product DNA was subsequently purified from the histone proteins and PNKP via an oligonucleotide cleanup kit (Zymo Research). Of note, this method routinely recovered >70% of the initial FAM-labeled DNAs, with minimal difference in recovery between oligos of different DNA lengths (SI Appendix, Fig. S27). The recovered substrate and product DNA were then treated with T4 DNA ligase for 1 hour at 37 °C to convert the PNKP product bearing a 3′-OH and 5′-phosphate terminus into a ligated DNA product. Of note, PNKP substrate DNA bearing a 3′-phosphate and 5′-OH, or individual phosphatase and kinase products bearing a 3′-OH and 5′-OH or 3′-phosphate and 5′-phosphate, respectively, are not converted into a ligated product by T4 DNA ligase. The substrate and product bands were then separated by 8% denaturing urea polyacrylamide gel electrophoresis (29:1 acrylamide:bis-acrylamide). The amount of substrate and ligated product at each time point was quantified using ImageJ^84^. Background subtraction was performed for each individual lane, where each lane was plotted as a density profile with a fixed selection area and the straight-line tool was used to draw a baseline under each peak. The resulting substrate and product values were then converted to a fraction product, and subsequently converted to a concentration of product (nM). The initial velocity for each reaction was determined by linear regression of product concentration as a function of time. All progress curves were linear (R^2^ > 0.9) over the time course for each reaction. The product concentration (nM) at each time point and the calculated initial velocities represent the mean of the three independent replicate experiments ± the standard deviation of the three independent replicate experiments.

### Cryo-EM sample and grid preparation

PNKP-Nick-NCP complexes were generated by mixing PNKP D171A (2.0 μM) with each individual Nick-NCP (1.0 μM) in a buffer containing 25 mM HEPES (pH 7.5), 25 mM NaCl, 1 mM TCEP, and 1 mM EDTA. The complexes were formed with the PNKP D171A mutant in the absence of Mg^2+^ and ATP to generate a pre-catalytic complex prior to 3′-phosphate hydrolysis and 5′-OH phosphorylation. The PNKP-Nick-NCP complexes were incubated for 5 mins at 4 °C prior to stabilization of the complex via glutaraldehyde crosslinking (0.2 %) for 15 min at 4 °C. The PNKP-Nick-NCP complexes were then immediately purified via a Superdex S200 Increase 10/300 GL (Cytiva) equilibrated in a buffer containing 25 mM HEPES (pH 7.5), 25 mM NaCl, 1 mM TCEP, and 1 mM EDTA. Fractions containing the PNKP-Nick-NCP complexes were combined and concentrated to 1.2 μM for PNKP-Nick-NCP−5, 1.2 μM for PNKP-Nick-NCP−4, 1.2 μM for PNKP-Nick-NCP−3 complexes. The final cryo-EM sample quality was assessed by native PAGE (5%, 59:1 acrylamide:bis-acrylamide ratio), and the native PAGE gels of the final cryo-EM samples can be found in SI Appendix, Figs. S4A, S7A, and S8A. Cryo-EM grids were generated by applying each PNKP-Nick-NCP complex (3 μL) to a glow discharged Quantifoil R2/1 300 mesh copper cryo-EM grid. The cryo-EM grids were blotted for 3.5 – 4s at 4 °C and 95% humidity and plunge-frozen in liquid ethane using a Vitrobot Mark IV (Thermo Fisher Scientific).

### Cryo-EM data collection and processing

All cryo-EM data collections were performed at the University of Virginia Molecular Electron Microscopy Facility on a 300 kV TFS Titan Krios cryo-TEM equipped with a Gatan K3/GIF direct electron detector using EPU. All data collection parameters can be found in SI Appendix, Tables S1-S3. Each cryo-EM dataset was processed using cryoSPARC^85^ following a similar general processing workflow (see SI Appendix, Figs. S4, S7, and S8). The micrographs for each dataset were initially imported into cryoSPARC and subjected to pre-processing using patch motion correction and patch CTF-estimation. An initial cycle of blob particle picking was then performed to generate a training set for Topaz particle picking^86^. Topaz was then used to generate the final particle stack for downstream classification. An ab initio model was generated using the final particle stack for each dataset, and heterogeneous refinement(s) and/or 3D classification(s) were performed to separate the Nick-NCPs and PNKP-Nick-NCP complexes. Following classification, the Nick-NCPs and PNKP-Nick-NCP reconstructions were subjected to local CTF refinement and reference-based motion correction prior to a final non-uniform refinement. To improve interpretability of PNKP-Nick-NCP reconstructions, the maps were further subjected to local refinement without particle subtraction using a mask for the PNKP catalytic domains and the adjacent nucleosomal DNA, and/or the PNKP FHA domain. A composite map was then generated by combining the consensus and local refinement maps using PHENIX combine focus maps^87^. The local resolution estimation for each composite map was generated using cryoSPARC local resolution estimation^85^ after combining the half maps from the consensus and local refinements using PHENIX combine focus maps^87^.

### Model building and refinement

An initial nucleosome model was generated from a previously determined cryo-EM structure of a non-damaged nucleosome (PDB: 10XZ)^43^. The initial PNKP model was generated from a previously determined high-resolution X-ray crystal structure of murine PNKP catalytic domains (PDB: 3ZVN)^49^where the murine side chain residues were manually mutated to the human side chain residues, or a high-resolution X-ray crystal structure of human PNKP FHA domain (PDB: 2W3O)^34^. For the Nick-NCP structures, the initial nucleosome model was rigid-body docked into each respective map using ChimeraX^88^, and the models further refined using PHENIX^87^ and Coot^89^ with protein and DNA secondary structure restraints. For the PNKP-Nick-NCP structures, the initial nucleosome and PNKP models were rigid-body docked into the map using ChimeraX^88^, and the models further refined using PHENIX^87^ and Coot^89^ with protein and DNA secondary structure restraints. Each model was validated using MolProbity^90^ and PHENIX comprehensive cryo-EM validation^87^ prior to deposition into the Protein Data Bank (PDB).

## Acknowledgements

This research was supported by startup funds from the University of Virginia School of Medicine and the University of Virginia Comprehensive Cancer Center (T.M.W.), the National Institute of General Medical Sciences R35GM165658 (T.M.W.), as well as Farrow Fellowship funding within the University of Virginia Comprehensive Cancer Center (D.J.B.). This work used the 200 kV TFS Glacios cryo-TEM and the 300 kV TFS Titan Krios cryo-TEM within the University of Virginia Molecular Electron Microscopy Core, which is supported by the University of Virginia School of Medicine (RRID:SCR_019031) and the University of Virginia Comprehensive Cancer Center NCI Cancer Center Support Grant P30-CA044579. The Titan Krios (S10-RR025067) and K3/GIF (U24-GM116790) within the Molecular Electron Microscopy Core were purchased in part or in full using designated NIH grants. We also acknowledge Michael Purdy, Ph.D. and David Cooper, Ph.D. from the Molecular Electron Microscopy Core at the University of Virginia School of Medicine for their generous help with cryo-EM screening and data collection.

## Author Contributions

D.J.B., N.I.M., and T.M.W. conceptualized the experiments and established the general research goals. D.J.B., D.K.M., and N.I.M. generated recombinant nucleosomes and recombinant proteins for biochemical assays and cryo-EM. D.J.B., D.K.M., N.I.M., and T.M.W. performed and analyzed the biochemical experiments. D.J.B. performed cryo-EM sample preparation and validation. D.J.B., J.J.E., and T.M.W. processed and analyzed the cryo-EM datasets. D.J.B., J.J.E., and T.M.W. performed model building and refinement. D.J.B. and T.M.W. wrote the manuscript, with input from D.K.M., N.I.M. and J.J.E.

## Data, Material, and Software Availability

The model coordinates for all Nick-NCP and PNKP-Nick-NCP structures were deposited into the PDB under accession number 37XU for Nick-NCP−5, 37XP for Nick-NCP−4, 37XL for Nick-NCP−3, 37XV for PNKP-Nick-NCP−5 (catalytic), 37XQ for PNKP-Nick-NCP−4 (catalytic), 37XM for PNKP-Nick-NCP−3 (catalytic), 37XW for PNKP-Nick-NCP−5 (catalytic-FHA), 37XS for PNKP-Nick-NCP−4 (catalytic-FHA), 37XO for PNKP-Nick-NCP−3 (catalytic-FHA), 37XX for PNKP-Nick-NCP−5 (FHA), 37XT for PNKP-Nick-NCP−4 (FHA), and 37XN for PNKP-Nick-NCP−3 (FHA). The consensus cryo-EM maps for all Nick-NCP and PNKP-Nick-NCP structures were deposited into the Electron Microscopy Data Bank (EMDB) under accession number EMD-78608 for Nick-NCP−5, EMD-78604 for Nick-NCP−4, EMD-78600 for Nick-NCP−3, EMD-78626 for PNKP-Nick-NCP−5 (catalytic), EMD-78619 for PNKP-Nick-NCP−4 (catalytic), EMD-78612 for PNKP-Nick-NCP−3 (catalytic), EMD-78628 for PNKP-Nick-NCP−5 (catalytic-FHA), EMD-78621 for PNKP-Nick-NCP−4 (catalytic-FHA), EMD-78614 for PNKP-Nick-NCP−3 (catalytic-FHA), EMD-78631 for PNKP-Nick-NCP−5 (FHA), EMD-78624 for PNKP-Nick-NCP−4 (FHA), and EMD-78617 for PNKP-Nick-NCP−3 (FHA). The composite cryo-EM maps for PNKP-Nick-NCP structures were deposited into the EMDB under accession number EMD-78609 for PNKP-Nick-NCP−5 (catalytic), EMD-78605 for PNKP-Nick-NCP−4 (catalytic), EMD-78601 for PNKP-Nick-NCP−3 (catalytic), EMD-78610 for PNKP-Nick-NCP−5 (catalytic-FHA), EMD-78606 for PNKP-Nick-NCP−4 (catalytic-FHA), EMD-78603 for PNKP-Nick-NCP−3 (catalytic-FHA), EMD-78611 for PNKP-Nick-NCP−5 (FHA), EMD-78607 for PNKP-Nick-NCP−4 (FHA), and EMD-78602 for PNKP-Nick-NCP−3 (FHA). The PNKP local refine cryo-EM maps for PNKP-Nick-NCP structures were deposited into the EMDB under accession number EMD-78627 for PNKP-Nick-NCP−5 (catalytic, PNKP/DNA local refine), EMD-78620 for PNKP-Nick-NCP−4 (catalytic, PNKP/DNA local refine), EMD-78613 for PNKP-Nick-NCP−3 (catalytic, PNKP/DNA local refine), EMD-78629 for PNKP-Nick-NCP−5 (catalytic-FHA, PNKP/DNA local refine), EMD-78630 for PNKP-Nick-NCP−5 (catalytic-FHA, FHA local refine), EMD-78623 for PNKP-Nick-NCP−4 (catalytic-FHA, PNKP/DNA local refine), EMD-78622 for PNKP-Nick-NCP−4 (catalytic-FHA, FHA local refine), EMD-78615 for PNKP-Nick-NCP−3 (catalytic-FHA, PNKP/DNA local refine), EMD-78616 for PNKP-Nick-NCP−3 (catalytic-FHA, FHA local refine), EMD-78632 for PNKP-Nick-NCP−5 (FHA, FHA local refine), EMD-78625 for PNKP-Nick-NCP−4 (FHA, FHA local refine), and EMD-78618 for PNKP-Nick-NCP−3 (FHA, FHA local refine). All other data are included in the article and/or Supporting Information file.

