## Supporting Information File for "Structural basis of end processing in the nucleosome by polynucleotide kinase-phosphatase"

**This PDF file includes:**

Figures S1 – S27
Tables S1 – S5

**Fig. S1: Validation of recombinant NCPs and recombinant PNKP proteins**

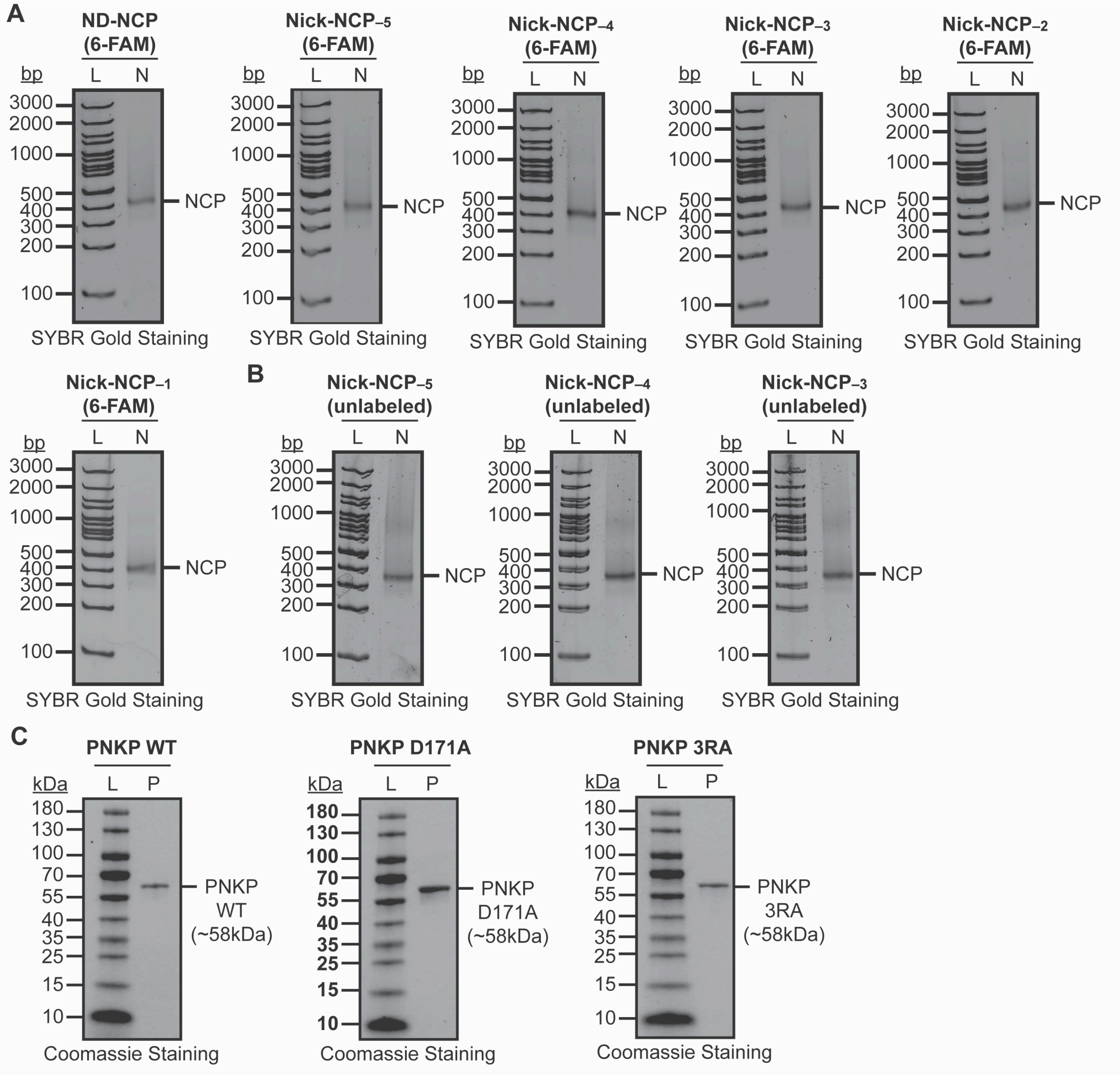

**Fig. S1: Validation of recombinant NCPs and recombinant PNKP proteins**

**(A)** Native PAGE gels confirming nucleosome formation and purity for the fluorescein (FAM)-labeled non-damaged NCP (ND-NCP), Nick-NCP$_{-5}$, Nick-NCP$_{-4}$, Nick-NCP$_{-3}$, Nick-NCP$_{-2}$, and Nick-NCP$_{-1}$ samples used for biochemical assays. The ND-NCP and Nick-NCPs were detected via SYBR Gold staining. The lanes corresponding to 100 bp DNA ladder (L) and the ND-NCP or Nick-NCP sample (N) are labeled. **(B)** Native PAGE gels confirming nucleosome formation and purity for the Nick-NCP$_{-5}$, Nick-NCP$_{-4}$, Nick-NCP$_{-3}$ samples used for cryo-EM. The Nick-NCPs were detected via SYBR Gold staining. The lanes corresponding to 100 bp DNA ladder (L) and each Nick-NCP sample (N) are labeled. **(C)** SDS-PAGE gel confirming the purity of the PNKP WT (left), PNKP D171A (middle), and PNKP 3RA (right) recombinant proteins. The recombinant proteins were detected with Coomassie blue staining. The lanes corresponding to the protein ladder (L) and PNKP protein (P) are labeled.

**Fig. S2: EMSA analysis of PNKP nucleosome binding**

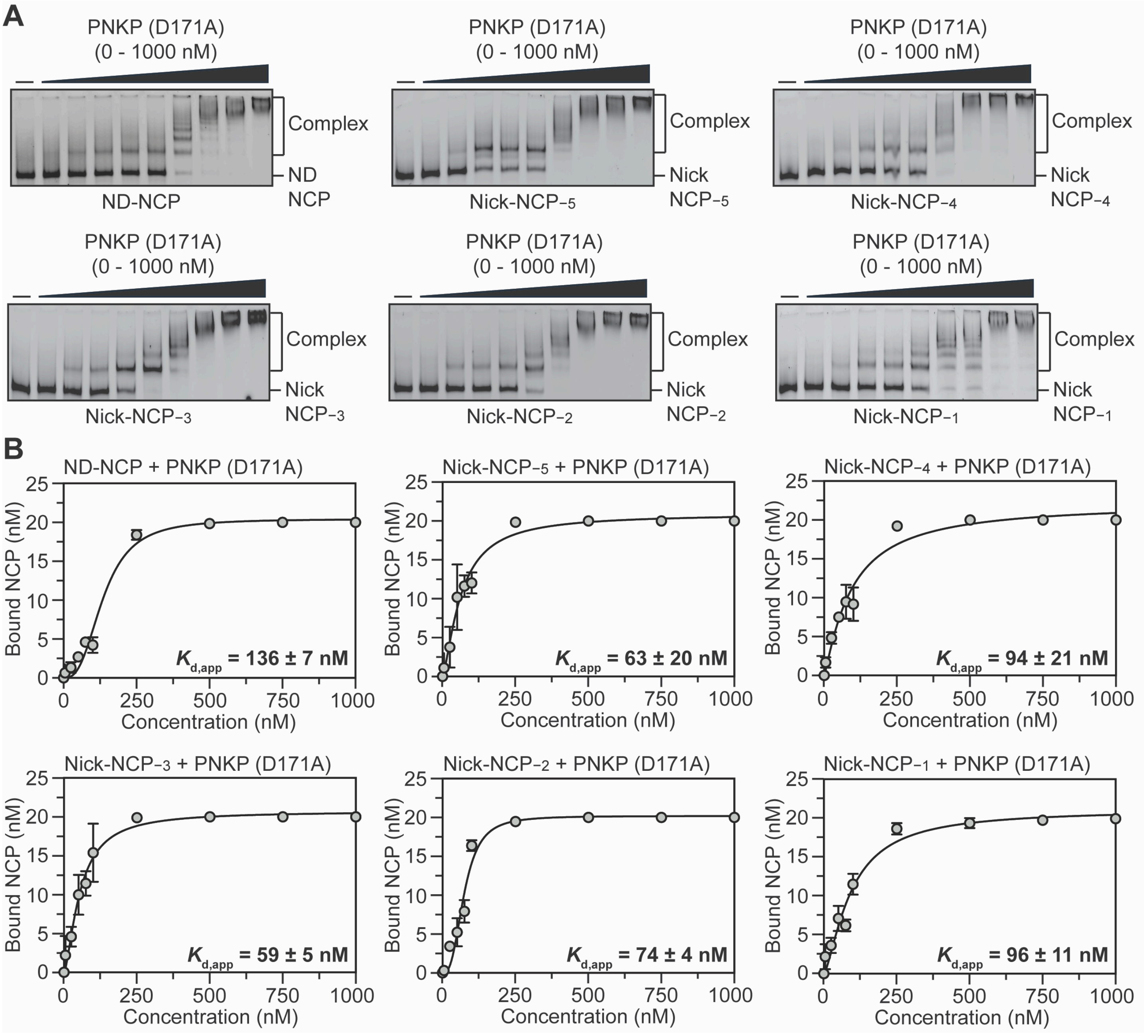

**Fig. S2: EMSA analysis of PNKP nucleosome binding**

**(A)** Representative native PAGE gels from the electrophoretic mobility shift assays (EMSAs) for PNKP D171A and the ND-NCP, Nick-NCP$_{-5}$, Nick-NCP$_{-4}$, Nick-NCP$_{-3}$, Nick-NCP$_{-2}$, and Nick-NCP$_{-1}$. The unbound NCP and PNKP-bound NCP complexes were detected using the 6-FAM label on the nucleosomal DNA. The gels are representative of three independent EMSA experiments performed for PNKP and the ND-NCP, Nick-NCP$_{-5}$, Nick-NCP$_{-4}$, Nick-NCP$_{-3}$, Nick-NCP$_{-2}$, and Nick-NCP$_{-1}$. **(B)** Quantification of the EMSA experiments for PNKP D171A and ND-NCP, Nick-NCP$_{-5}$, Nick-NCP$_{-4}$, Nick-NCP$_{-3}$, Nick-NCP$_{-2}$, and Nick-NCP$_{-1}$. The data points represent the mean ± standard deviation from the three independent replicate experiments. The error bars are included for all experimental data points, but some error bars are smaller than the circles used to represent data points. The apparent binding affinity ($K_{d,app}$) is shown as an inset for each experiment and represents the mean ± standard deviation from the three independent replicate experiments.

**Fig. S3: Kinetic analysis of PNKP end processing in the nucleosome**

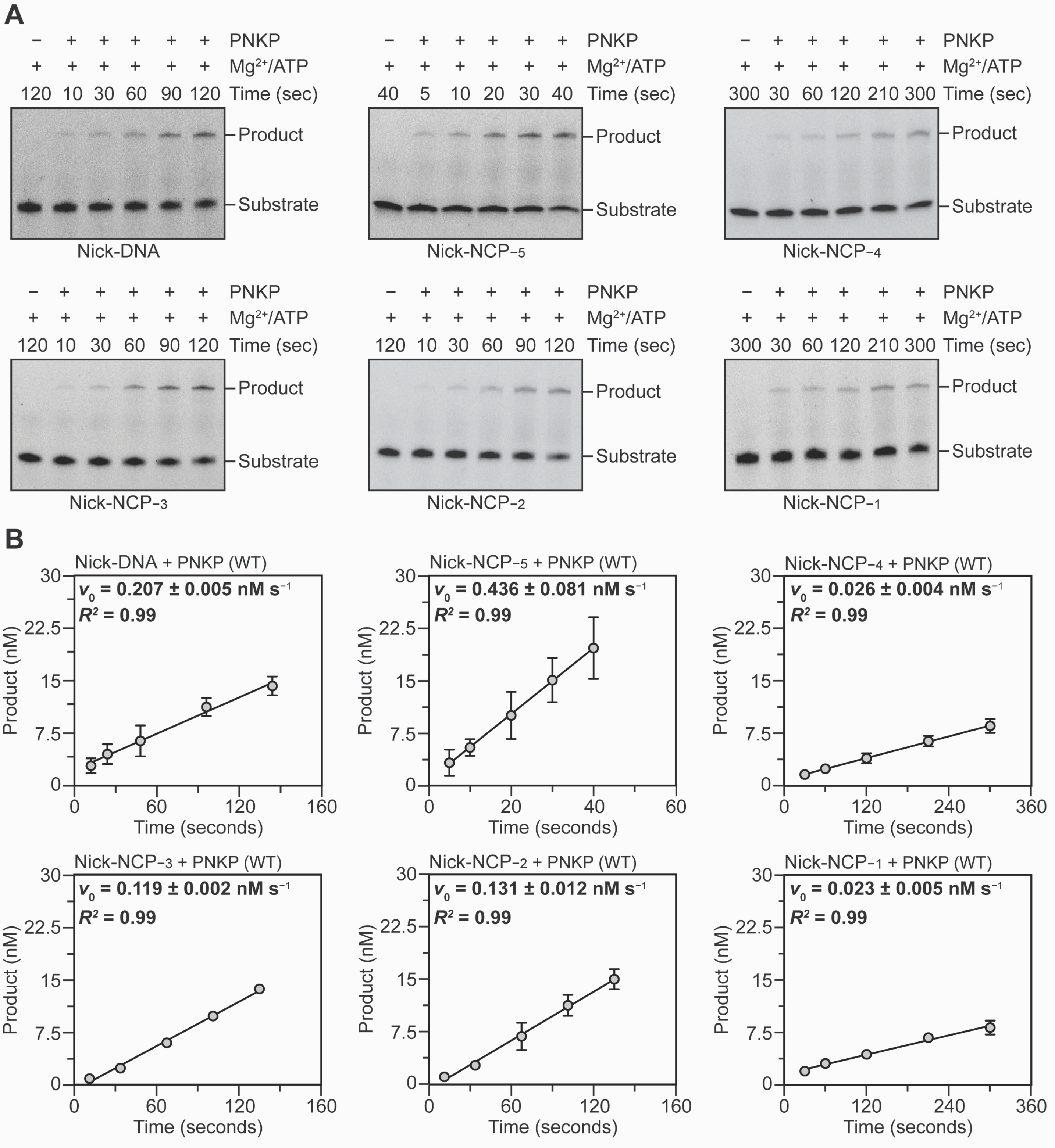

**Fig. S3: Kinetic analysis of PNKP end processing in the nucleosome**

**(A)** Representative denaturing Urea-PAGE gels from the PNKP enzyme kinetics on Nick-DNA, Nick-NCP$_{-5}$, Nick-NCP$_{-4}$, Nick-NCP$_{-3}$, Nick-NCP$_{-2}$, and Nick-NCP$_{-1}$. The substrate and product were detected using the 6-FAM label on the nucleosomal DNA. The gels are representative of three independent replicate experiments performed for PNKP and Nick-DNA, Nick-NCP$_{-5}$, Nick-NCP$_{-4}$, Nick-NCP$_{-3}$, Nick-NCP$_{-2}$, and Nick-NCP$_{-1}$. **(B)** Progress curves for the PNKP enzyme kinetics on Nick-DNA, Nick-NCP$_{-5}$, Nick-NCP$_{-4}$, Nick-NCP$_{-3}$, Nick-NCP$_{-2}$, and Nick-NCP$_{-1}$. The data points represent the mean ± standard deviation from the three independent replicate experiments. The error bars are included for all experimental data points, but some error bars are smaller than the circles used to represent data points. The initial velocity ($v_0$, nM $s^{-1}$) obtained from the linear regression of product formation as a function of time is shown as an inset for each experiment and represents the mean ± standard deviation from the three independent replicate experiments.

**Fig. S4: Cryo-EM processing workflow for the PNKP-Nick-NCP-3 complex**

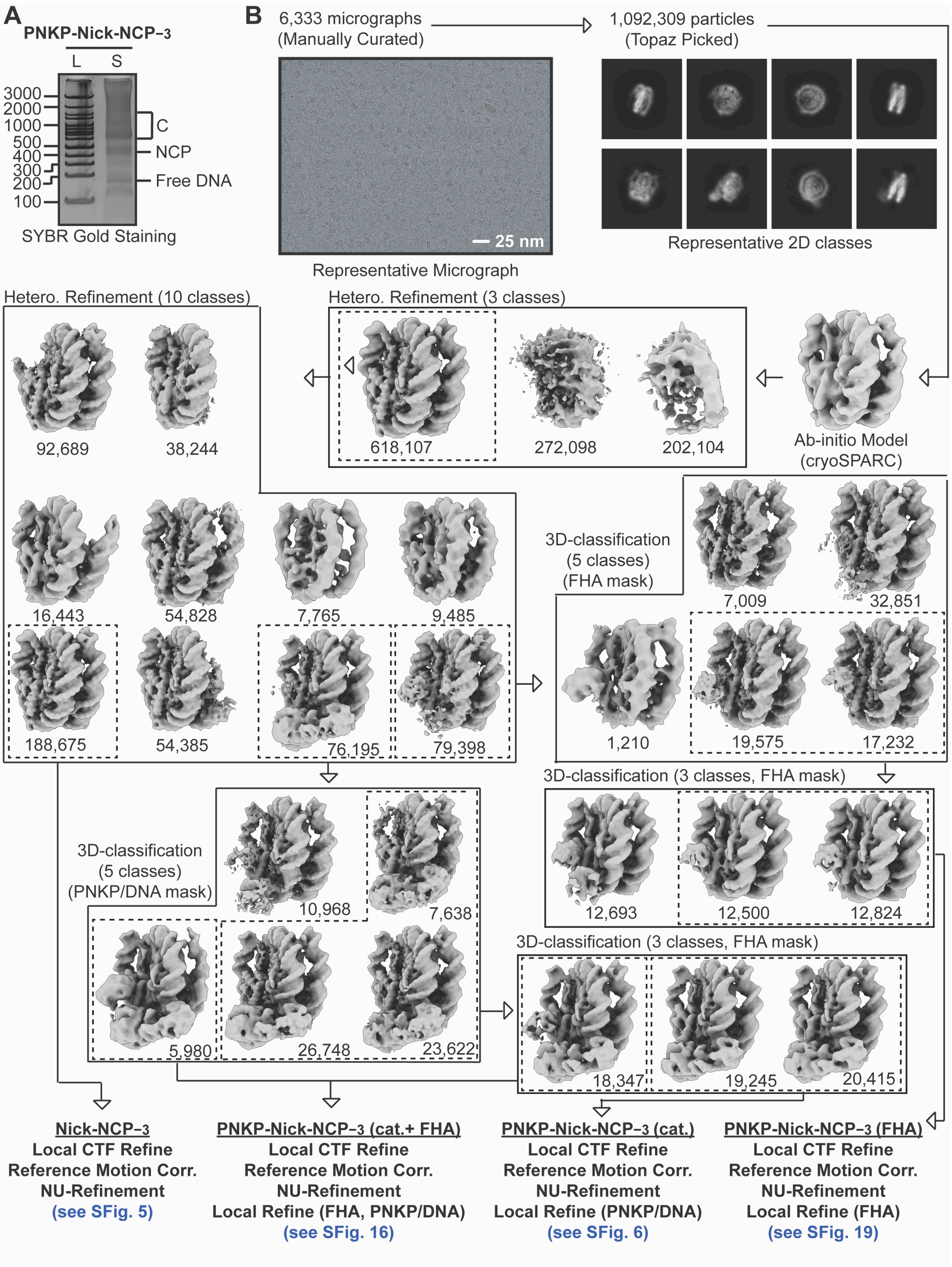

**Fig. S4: Cryo-EM processing workflow for the PNKP-Nick-NCP−3 complex**

**(A)** Native PAGE gel of a 100 bp DNA ladder (L) and the PNKP-Nick-NCP−3 cryo-EM sample (S). The PNKP-Nick-NCP−3 complexes were visualized with SYBR Gold staining. **(B)** Cryo-EM data processing workflow for the PNKP-Nick-NCP−3 dataset. A representative micrograph (n=6,333) and 2D classes from the PNKP-Nick-NCP−3 dataset are shown. The maps chosen for further classification and refinement throughout the cryo-EM data processing workflow are boxed. The final map, final model, and quality assessment metrics for the Nick-NCP−3, PNKP-Nick-NCP−3 (catalytic), PNKP-Nick-NCP−3 (catalytic-FHA), and PNKP-Nick-NCP−3 (FHA) can be found in Supplementary Figs. S5, S6, S16, and S19, respectively.

**Fig. S5: Nick-NCP-3 map and model quality assessment**

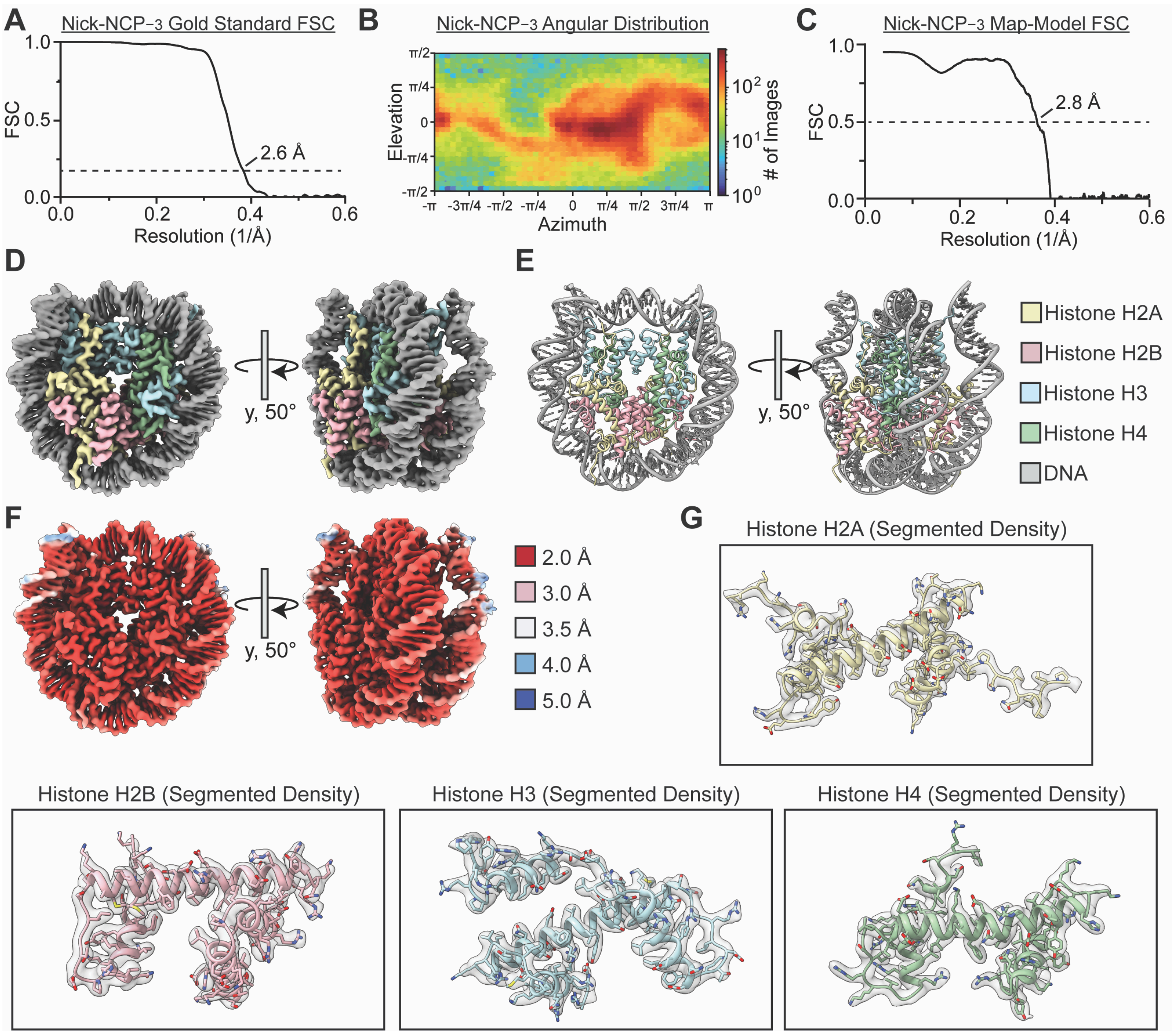

**Fig. S5: Nick-NCP−3 map and model quality assessment**

**(A**) Gold-standard Fourier shell correlation (GS-FSC) for the Nick-NCP−3 cryo-EM map. The dashed line corresponds to GS-FSC - 0.143. **(B)** Angular distribution heatmap for the Nick-NCP−3 cryo-EM map**. (C)** Map-to-model FSC for the Nick-NCP−3 model and cryo-EM map. The dashed line corresponds to FSC - 0.5. **(D)** The final 2.6 Å Nick-NCP−3 cryo-EM map shown in two different orientations. **(E)** The final Nick-NCP−3 model shown in two different orientations. **(F)** Local resolution estimation for the Nick-NCP−3 cryo-EM map shown in two different orientations. **(G)** Representative segmented densities for histones H2A, H2B, H3, and H4 from the Nick-NCP−3 cryo-EM map. The representative segmented densities from the cryo-EM map are shown as transparent gray surfaces.

**Fig. S6: PNKP-Nick-NCP-3 (catalytic) map and model quality assessment**

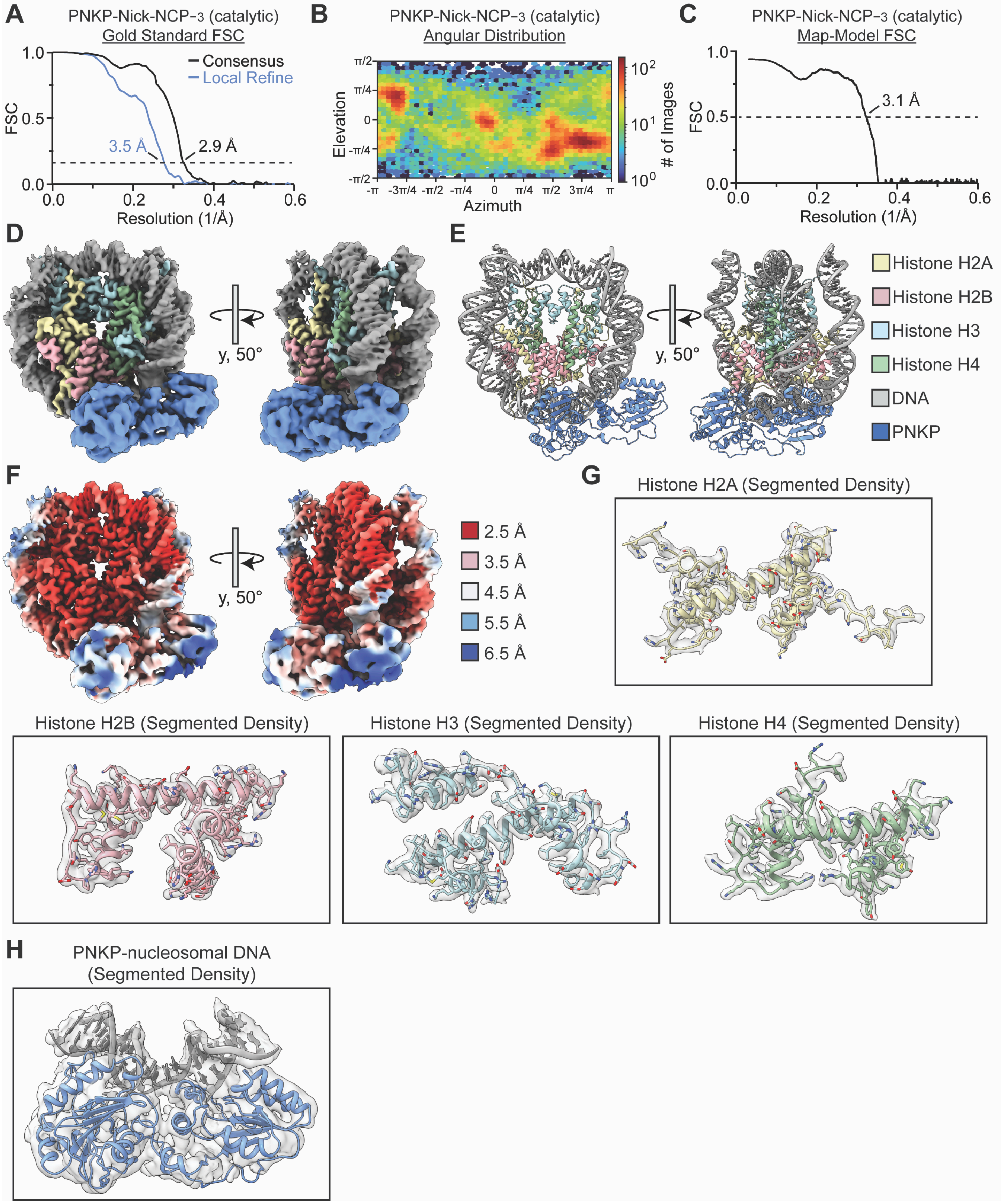

**Fig. S6: PNKP-Nick-NCP$_{-3}$ (catalytic) map and model quality assessment**

**(A**) Gold-standard Fourier shell correlation (GS-FSC) for the PNKP-Nick-NCP$_{-3}$ (catalytic) consensus cryo-EM map (solid black line) and the PNKP/nucleosomal DNA focus cryo-EM map (solid blue line). The dashed line corresponds to GS-FSC - 0.143. **(B)** Angular distribution heatmap for the PNKP-Nick-NCP$_{-3}$ (catalytic) cryo-EM map**. (C)** Map-to-model FSC for the PNKP-Nick-NCP$_{-3}$ (catalytic) model and composite cryo-EM map. The dashed line corresponds to FSC - 0.5. **(D)** The final 2.9 Å PNKP-Nick-NCP$_{-3}$ (catalytic) composite cryo-EM map shown in two different orientations. **(E)** The final PNKP-Nick-NCP$_{-3}$ (catalytic) model shown in two different orientations. **(F)** Local resolution estimation for the PNKP-Nick-NCP$_{-3}$ (catalytic) composite cryo-EM map shown in two different orientations. **(G)** Representative segmented densities for histones H2A, H2B, H3, and H4 from the PNKP-Nick-NCP$_{-3}$ (catalytic) cryo-EM map. The representative segmented densities from the cryo-EM map are shown as transparent gray surfaces. **(H)** Representative segmented densities for the PNKP catalytic domains and the surrounding nucleosomal DNA from the PNKP-Nick-NCP$_{-3}$ (catalytic) cryo-EM map. The representative segmented densities from the cryo-EM map are shown as transparent gray surfaces.

**Fig. S7: Cryo-EM processing workflow for the PNKP-Nick-NCP-4 complex**

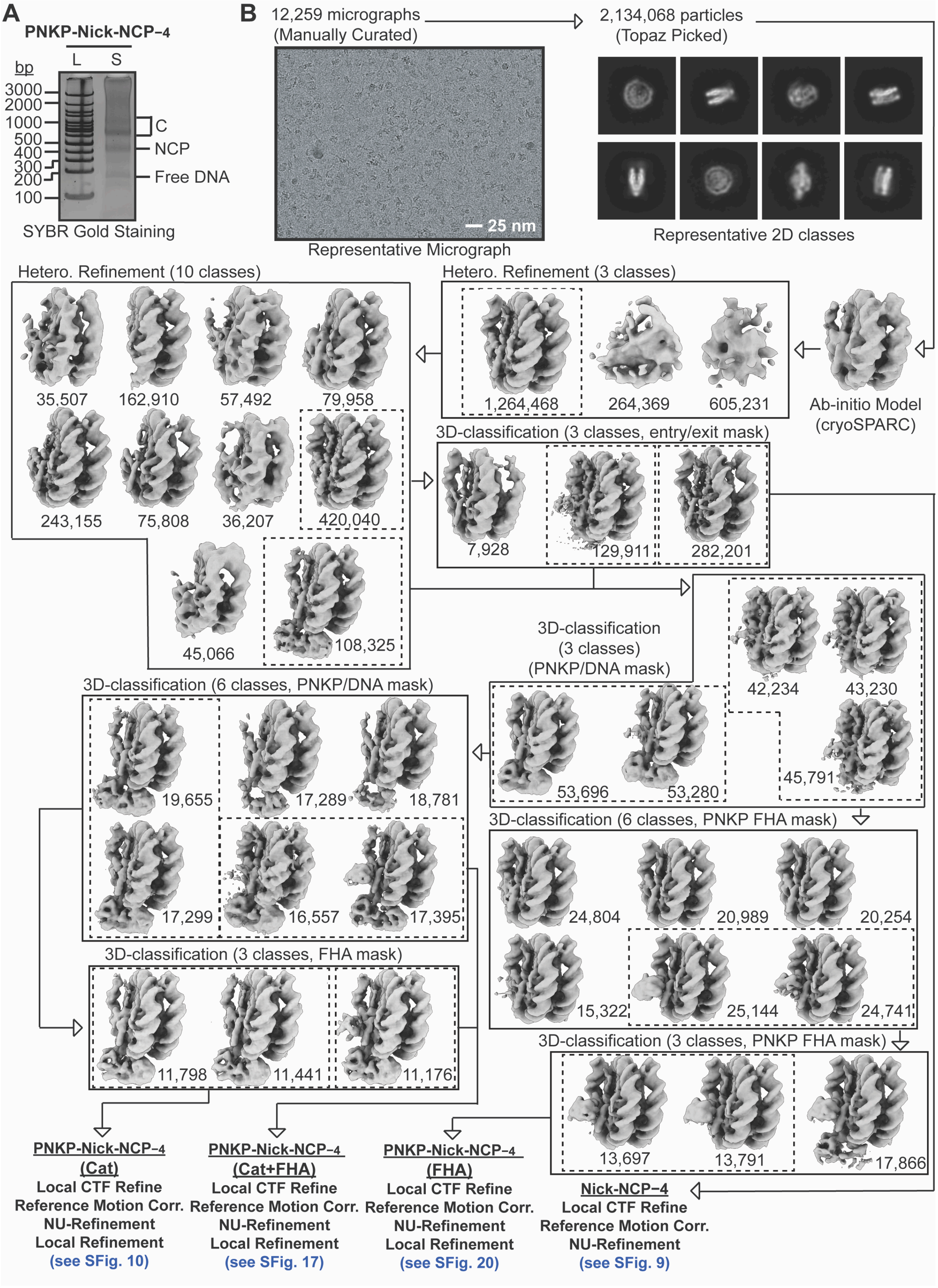

**Fig. S7: Cryo-EM processing workflow for the PNKP-Nick-NCP$_{-4}$ complex**

**(A)** Native PAGE gel of a 100 bp DNA ladder (L) and the PNKP-Nick-NCP$_{-4}$ cryo-EM sample (S). The PNKP-Nick-NCP$_{-4}$ complexes were visualized with SYBR Gold staining. **(B)** Cryo-EM data processing workflow for the PNKP-Nick-NCP$_{-4}$ dataset. A representative micrograph (n=12,259) and 2D classes from the PNKP-Nick-NCP$_{-4}$ dataset are shown. The maps chosen for further classification and refinement throughout the cryo-EM data processing workflow are boxed. The final map, final model, and quality assessment metrics for the Nick-NCP$_{-4}$, PNKP-Nick-NCP$_{-4}$ (catalytic), PNKP-Nick-NCP$_{-4}$ (catalytic-FHA), and PNKP-Nick-NCP$_{-4}$ (FHA) can be found in Supplementary Figs. S9, S10, S17, and S20, respectively.

**Fig. S8: Cryo-EM processing workflow for the PNKP-Nick-NCP-5 complex**

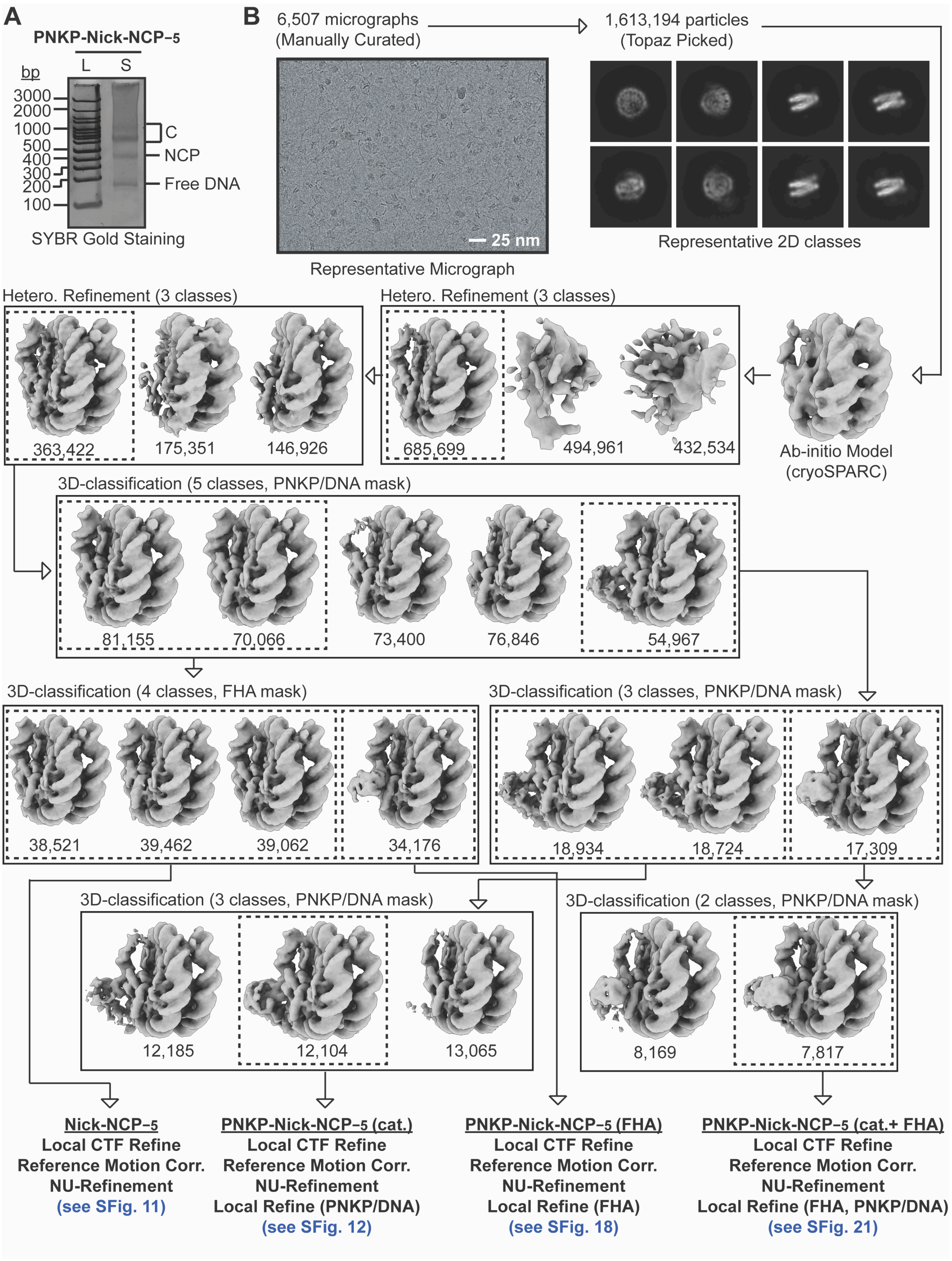

**Fig. S8: Cryo-EM processing workflow for the PNKP-Nick-NCP$_{-5}$ complex**

**(A)** Native PAGE gel of a 100 bp DNA ladder (L) and the PNKP-Nick-NCP$_{-5}$ cryo-EM sample (S). The PNKP-Nick-NCP$_{-5}$ complexes were visualized with SYBR Gold staining. **(B)** Cryo-EM data processing workflow for the PNKP-Nick-NCP$_{-5}$ dataset. A representative micrograph (n=6,507) and 2D classes from the PNKP-Nick-NCP$_{-5}$ dataset are shown. The maps chosen for further classification and refinement throughout the cryo-EM data processing workflow are boxed. The final map, final model, and quality assessment metrics for the Nick-NCP$_{-5}$, PNKP-Nick-NCP$_{-5}$ (catalytic), PNKP-Nick-NCP$_{-5}$ (catalytic-FHA), and PNKP-Nick-NCP$_{-5}$ (FHA) can be found in Supplementary Figs. S11, S12, S18, and S21, respectively.

**Fig. S9: Nick-NCP-4 map and model quality assessment**

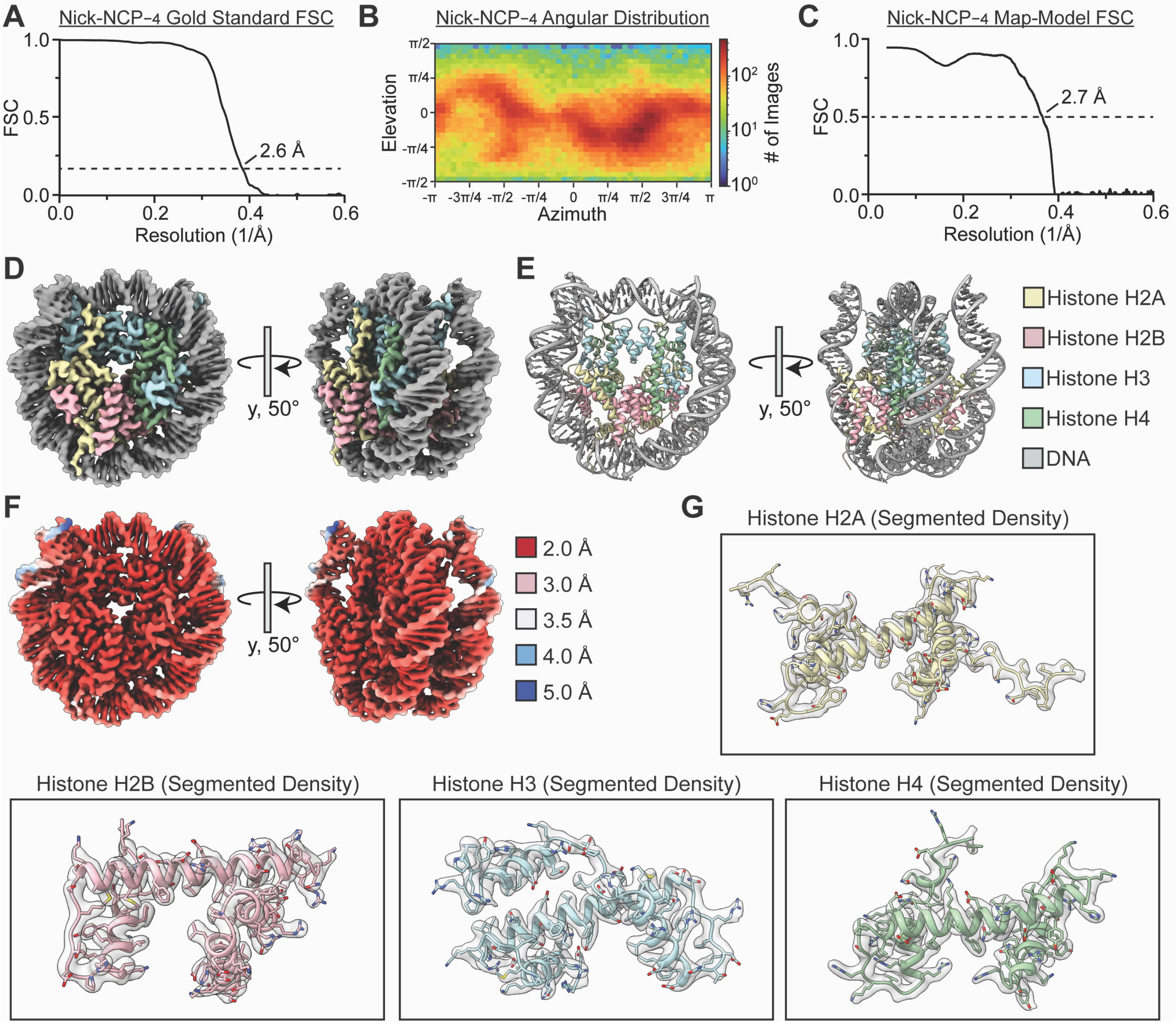

**Fig. S9: Nick-NCP−4 map and model quality assessment**

(**A**) Gold-standard Fourier shell correlation (GS-FSC) for the Nick-NCP−4 cryo-EM map. The dashed line corresponds to GS-FSC - 0.143. **(B)** Angular distribution heatmap for the Nick-NCP−4 cryo-EM map**. (C)** Map-to-model FSC for the Nick-NCP−4 model and cryo-EM map. The dashed line corresponds to FSC - 0.5. **(D)** The final 2.6 Å Nick-NCP−4 cryo-EM map shown in two different orientations. **(E)** The final Nick-NCP−4 model shown in two different orientations. **(F)** Local resolution estimation for the Nick-NCP−4 cryo-EM map shown in two different orientations. **(G)** Representative segmented densities for histones H2A, H2B, H3, and H4 from the Nick-NCP−4 cryo-EM map. The representative segmented densities from the cryo-EM map are shown as transparent gray surfaces.

**Fig. S10: PNKP-Nick-NCP−4 (catalytic) map and model quality assessment**

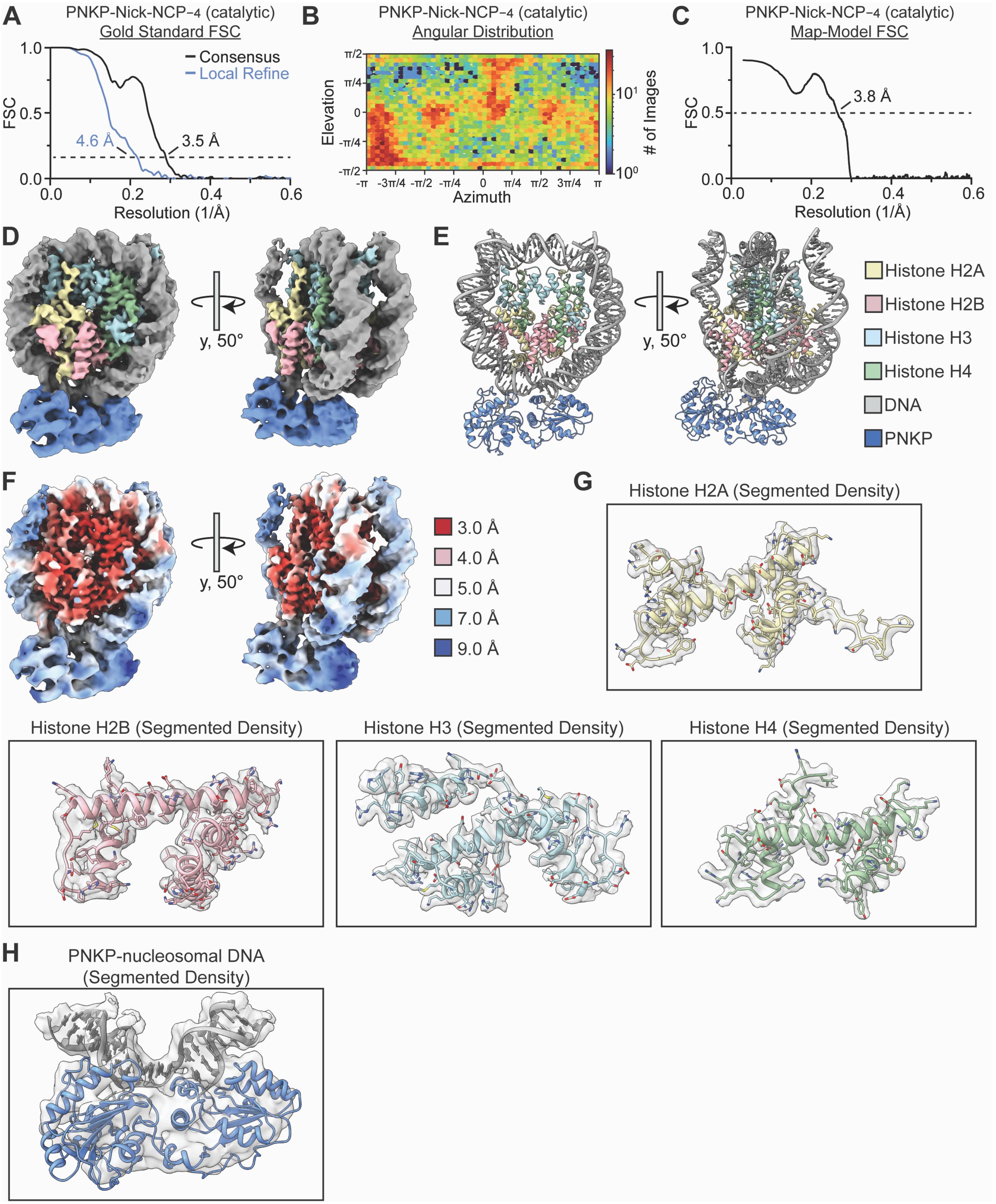

**Fig. S10: PNKP-Nick-NCP−4 (catalytic) map and model quality assessment**

**(A**) Gold-standard Fourier shell correlation (GS-FSC) for the PNKP-Nick-NCP−4 (catalytic) consensus cryo-EM map (solid black line) and the PNKP/nucleosomal DNA focus cryo-EM map (solid blue line). The dashed line corresponds to GS-FSC - 0.143. **(B)** Angular distribution heatmap for the PNKP-Nick-NCP−4 (catalytic) cryo-EM map**.** **(C)** Map-to-model FSC for the PNKP-Nick-NCP−4 (catalytic) model and composite cryo-EM map. The dashed line corresponds to FSC - 0.5. **(D)** The final 3.5 Å PNKP-Nick-NCP−4 (catalytic) composite cryo-EM map shown in two different orientations. **(E)** The final PNKP-Nick-NCP−4 (catalytic) model shown in two different orientations. **(F)** Local resolution estimation for the PNKP-Nick-NCP−4 (catalytic) composite cryo-EM map shown in two different orientations. **(G)** Representative segmented densities for histones H2A, H2B, H3, and H4 from the PNKP-Nick-NCP−4 (catalytic) cryo-EM map. The representative segmented densities from the cryo-EM map are shown as transparent gray surfaces. **(H)** Representative segmented densities for the PNKP catalytic domains and the surrounding nucleosomal DNA from the PNKP-Nick-NCP−4 (catalytic) cryo-EM map. The representative segmented densities from the cryo-EM map are shown as transparent gray surfaces.

**Fig. S11: Nick-NCP−5 map and model quality assessment**

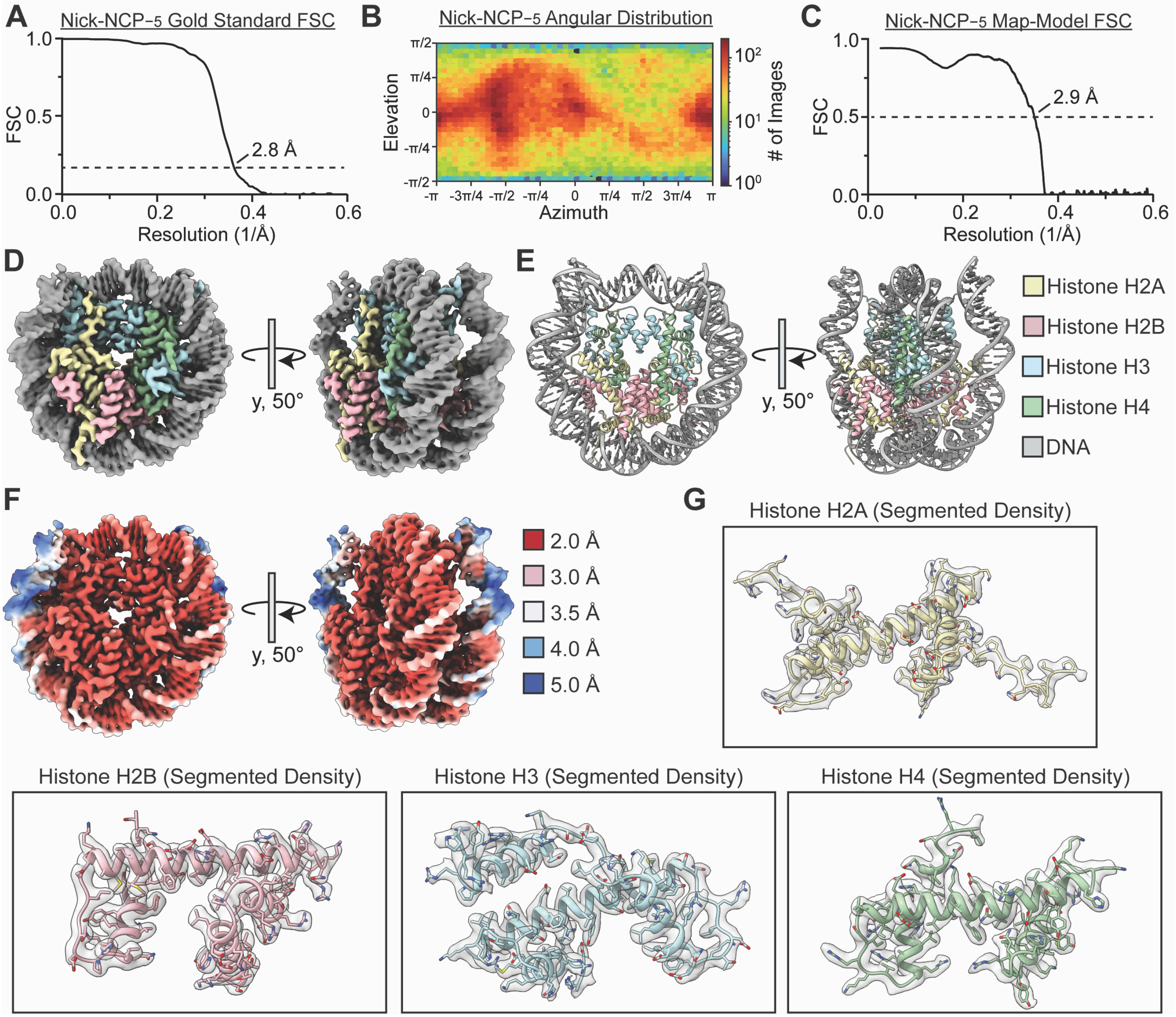

**Fig. S11: Nick-NCP−5 map and model quality assessment**

(**A**) Gold-standard Fourier shell correlation (GS-FSC) for the Nick-NCP−5 cryo-EM map. The dashed line corresponds to GS-FSC - 0.143. **(B)** Angular distribution heatmap for the Nick-NCP−5 cryo-EM map**. (C)** Map-to-model FSC for the Nick-NCP−5 model and cryo-EM map. The dashed line corresponds to FSC - 0.5. **(D)** The final 2.8 Å Nick-NCP−5 cryo-EM map shown in two different orientations. **(E)** The final Nick-NCP−5 model shown in two different orientations. **(F)** Local resolution estimation for the Nick-NCP−5 cryo-EM map shown in two different orientations. **(G)** Representative segmented densities for histones H2A, H2B, H3, and H4 from the Nick-NCP−5 cryo-EM map. The representative segmented densities from the cryo-EM map are shown as transparent gray surfaces.

**Fig. S12: PNKP-Nick-NCP−5 (catalytic) map and model quality assessment**

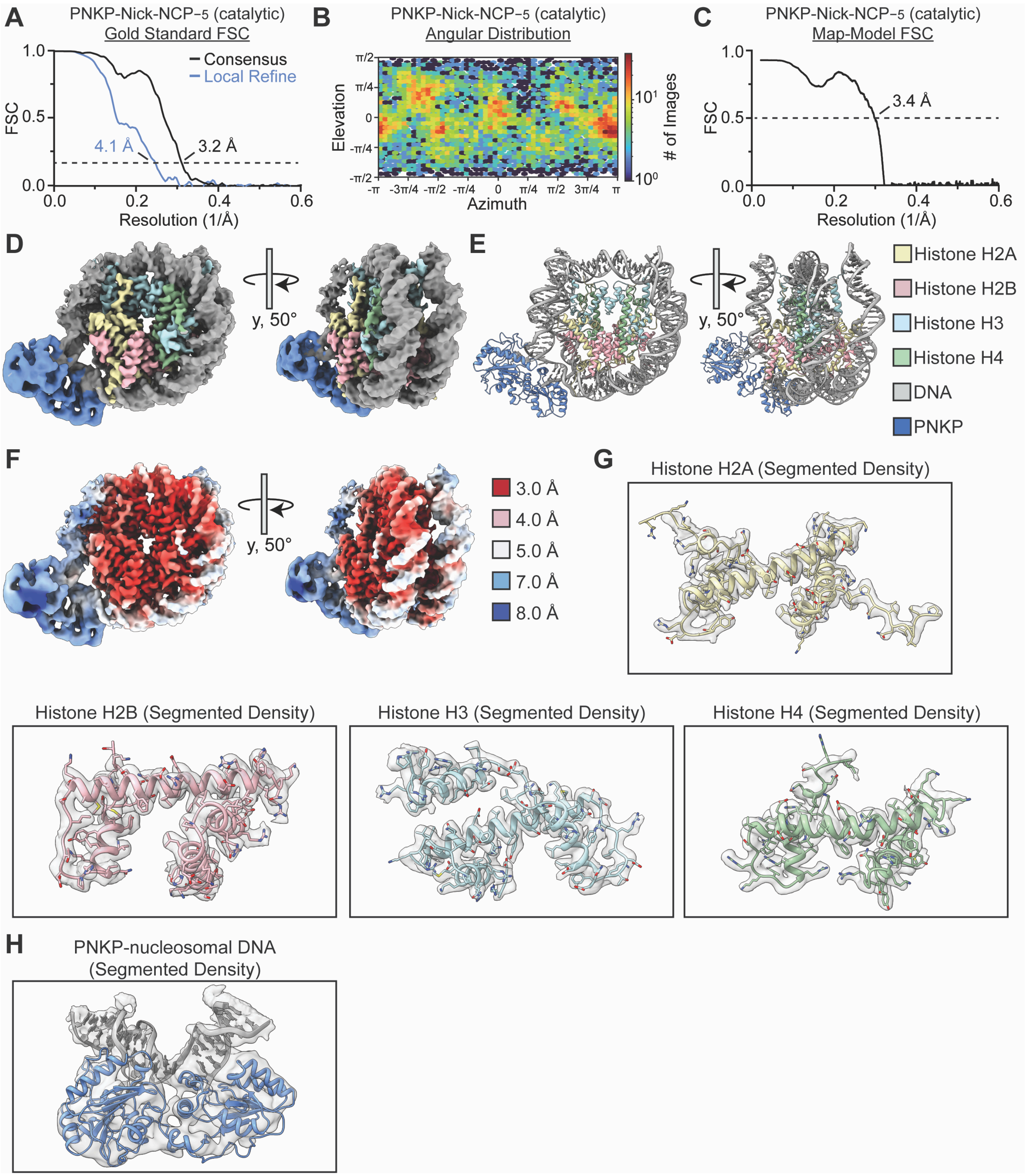

**Fig. S12: PNKP-Nick-NCP$_{-5}$ (catalytic) map and model quality assessment**

**(A**) Gold-standard Fourier shell correlation (GS-FSC) for the PNKP-Nick-NCP$_{-5}$ (catalytic) consensus cryo-EM map (solid black line) and the PNKP/nucleosomal DNA focus cryo-EM map (solid blue line). The dashed line corresponds to GS-FSC - 0.143. **(B)** Angular distribution heatmap for the PNKP-Nick-NCP$_{-5}$ (catalytic) cryo-EM map**.** **(C)** Map-to-model FSC for the PNKP-Nick-NCP$_{-5}$ (catalytic) model and composite cryo-EM map. The dashed line corresponds to FSC - 0.5. **(D)** The final 3.2 Å PNKP-Nick-NCP$_{-5}$ (catalytic) composite cryo-EM map shown in two different orientations. **(E)** The final PNKP-Nick-NCP$_{-5}$ (catalytic) model shown in two different orientations. **(F)** Local resolution estimation for the PNKP-Nick-NCP$_{-5}$ (catalytic) composite cryo-EM map shown in two different orientations. **(G)** Representative segmented densities for histones H2A, H2B, H3, and H4 from the PNKP-Nick-NCP$_{-5}$ (catalytic) cryo-EM map. The representative segmented densities from the cryo-EM map are shown as transparent gray surfaces. **(H)** Representative segmented densities for the PNKP catalytic domains and the surrounding nucleosomal DNA from the PNKP-Nick-NCP$_{-5}$ (catalytic) cryo-EM map. The representative segmented densities from the cryo-EM map are shown as transparent gray surfaces.

**Fig. S13: PNKP deforms the nucleosomal DNA during non-ligatable nick recognition at SHL−4**

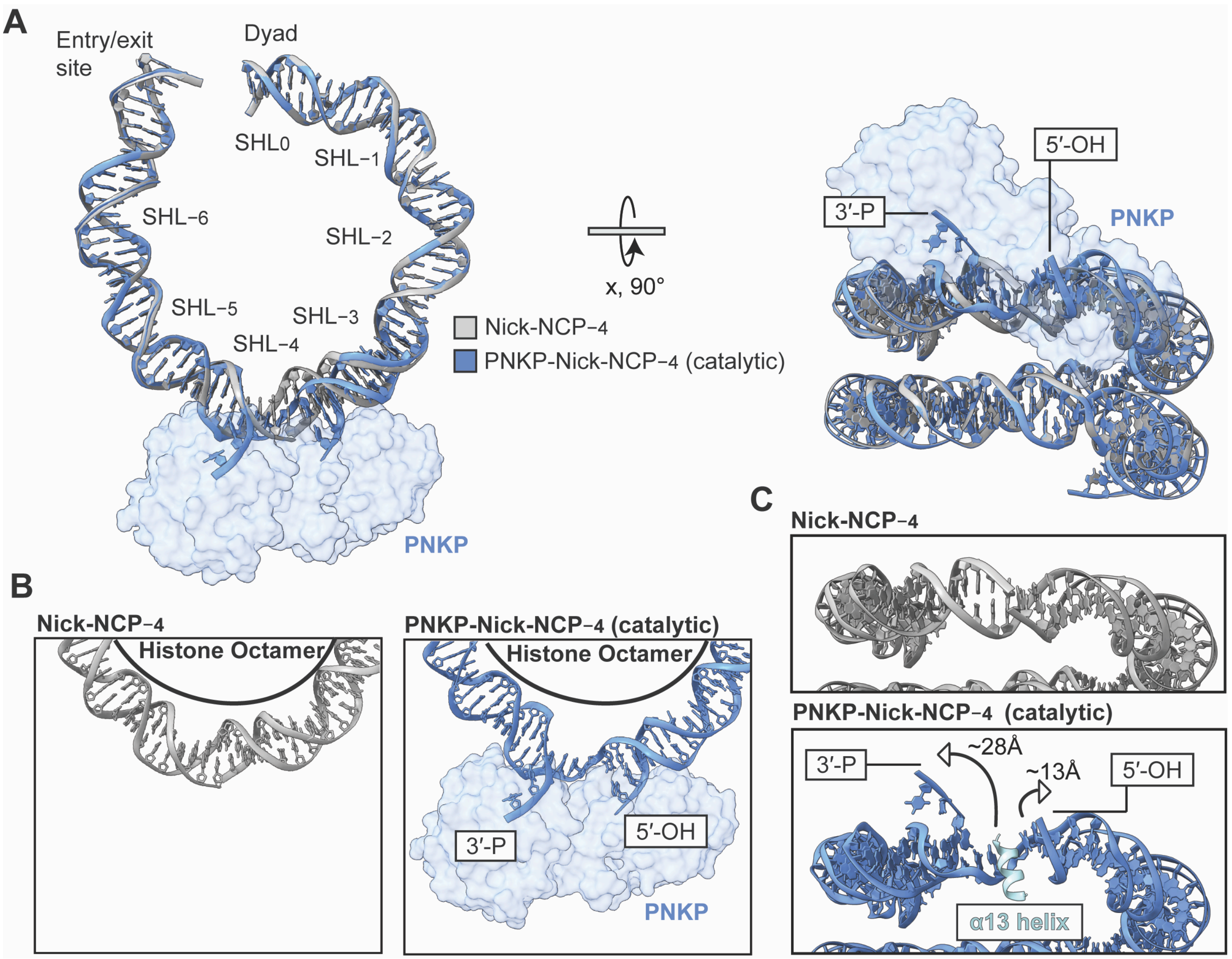

**Fig. S13: PNKP deforms the nucleosomal DNA during non-ligatable nick recognition at SHL−4**

**(A)** Structural comparison of the nucleosomal DNA in the Nick-NCP−4 and the PNKP-Nick-NCP−4 (catalytic) complex shown in two different orientations. The Nick-NCP−4 and PNKP-Nick-NCP−4 (catalytic) complex are shown in gray and blue, respectively. PNKP is shown as a transparent blue surface representation for improved clarity. **(B)** Focused views of the nucleosomal DNA from SHL−2.5 to SHL−5 in the Nick-NCP−4 (left) and PNKP-Nick-NCP−4 (catalytic) complex (right). **(C)** Focused views of the nucleosomal DNA from the inter-gyres perspective in the Nick-NCP−4 (top) and PNKP-Nick-NCP−4 (catalytic) complex (bottom). The α13 helix of the PNKP kinase domain is shown as a cartoon in light blue.

**Fig. S14: PNKP deforms the nucleosomal DNA during non-ligatable nick recognition at SHL−5**

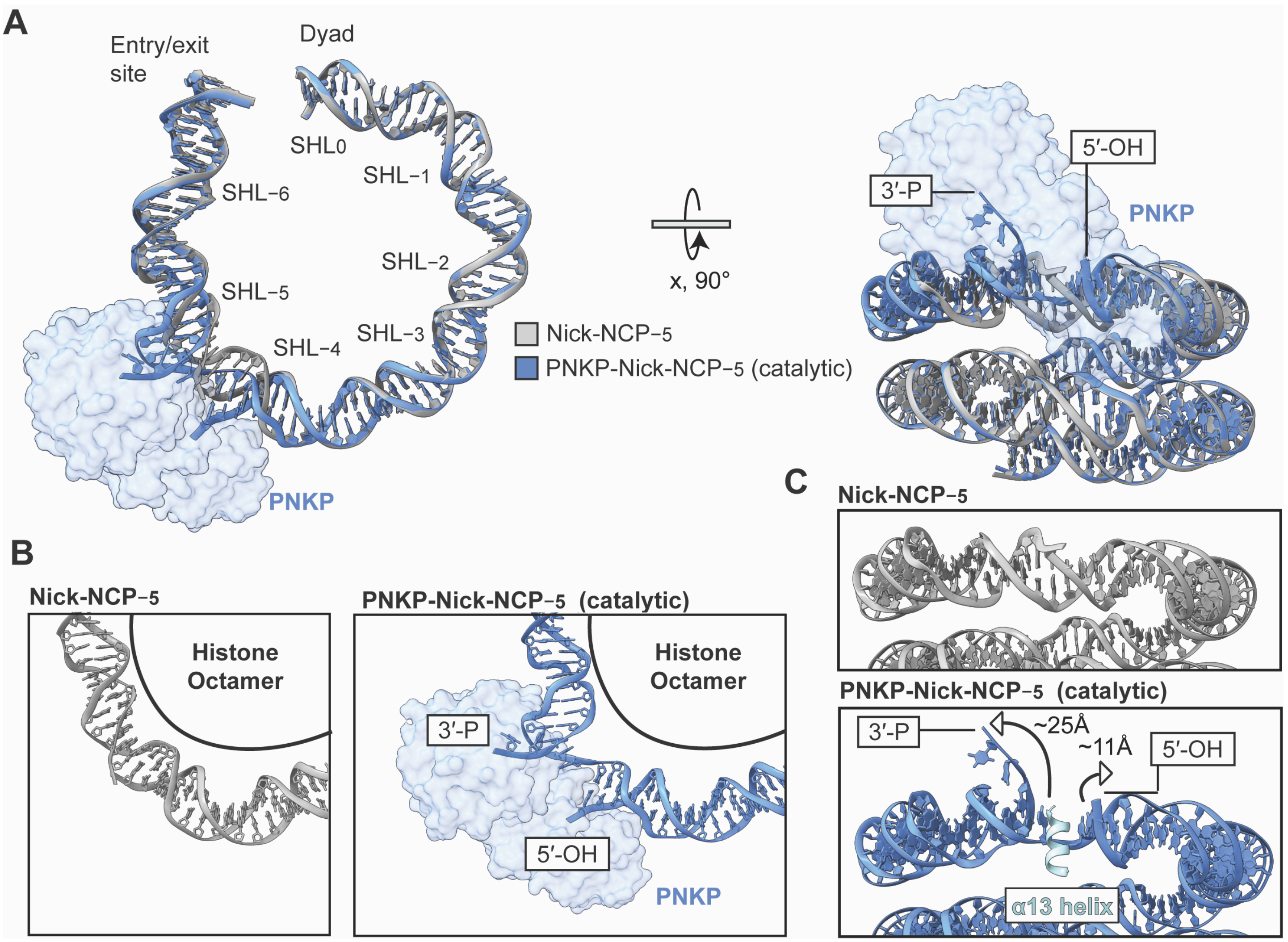

**Fig. S14: PNKP deforms the nucleosomal DNA during non-ligatable nick recognition at SHL−5**

**(A)** Structural comparison of the nucleosomal DNA in the Nick-NCP−5 and the PNKP-Nick-NCP−5 (catalytic) complex shown in two different orientations. The Nick-NCP−5 and PNKP-Nick-NCP−5 (catalytic) complex are shown in gray and blue, respectively. PNKP is shown as a transparent blue surface representation for improved clarity. **(B)** Focused views of the nucleosomal DNA from SHL−3.5 to SHL−6 in the Nick-NCP−5 (left) and PNKP-Nick-NCP−5 (catalytic) complex (right). **(C)** Focused views of the nucleosomal DNA from the inter-gyres perspective in the Nick-NCP−5 (top) and PNKP-Nick-NCP−5 (catalytic) complex (bottom). The α13 helix of the PNKP kinase domain is shown as a cartoon in light blue.

**Fig. S15: Structural comparison of the PNKP-Nick-NCP (catalytic) complex at SHL−3, SHL−4, and SHL−5**

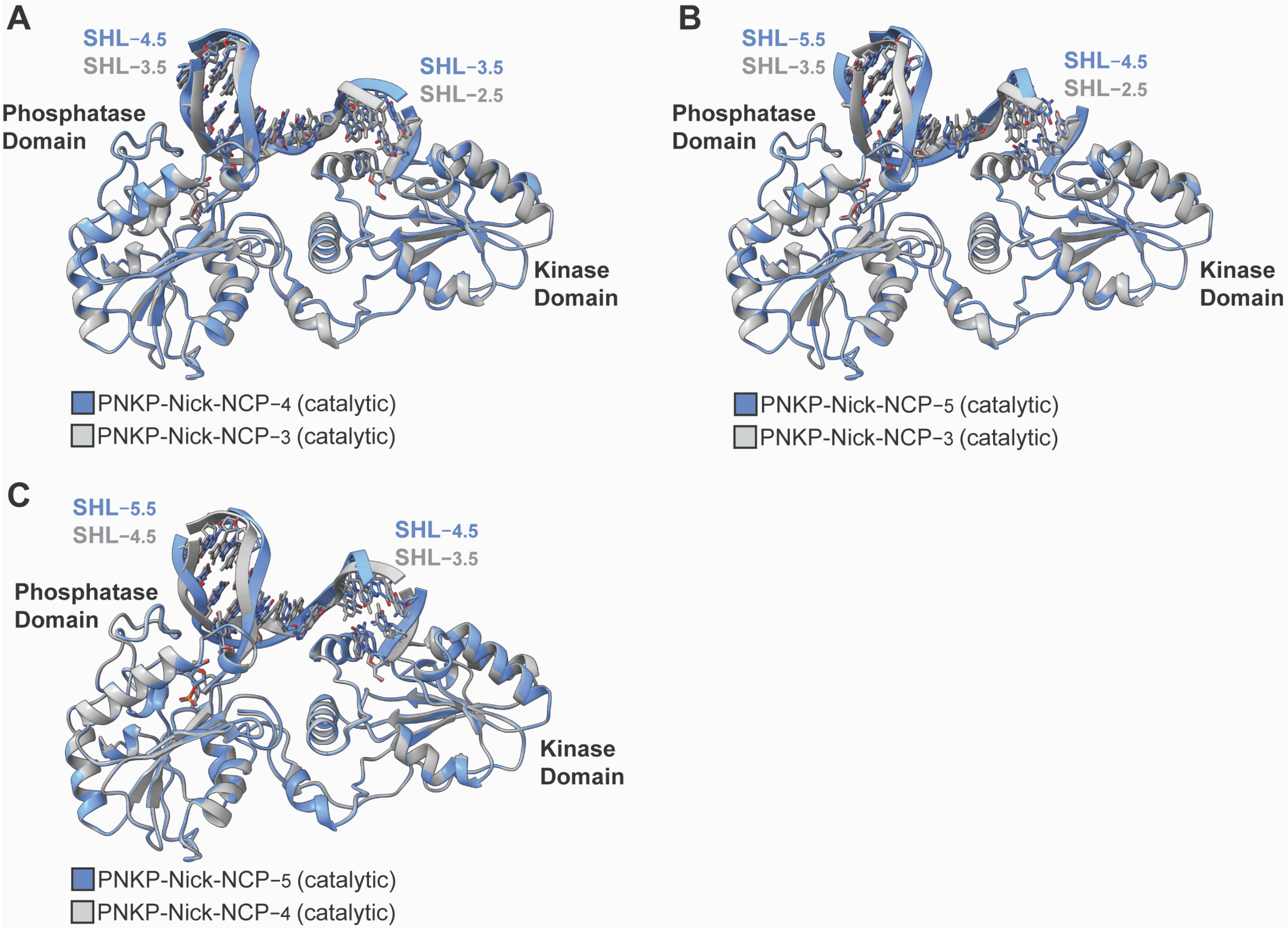

**Fig. S15: Structural comparison of the PNKP-Nick-NCP (catalytic) complex at SHL−3, SHL−4, and SHL−5**

**(A)** Structural overlay of the PNKP catalytic domains bound to the nucleosomal DNA in the PNKP-Nick-NCP−3 (catalytic) and PNKP-Nick-NCP−4 (catalytic) complexes. The PNKP-Nick-NCP−3 (catalytic) and PNKP-Nick-NCP−4 (catalytic) complexes are colored gray and blue, respectively. **(B)** Structural overlay of the PNKP catalytic domains bound to the nucleosomal DNA in the PNKP-Nick-NCP−3 (catalytic) and PNKP-Nick-NCP−5 (catalytic) complexes. The PNKP-Nick-NCP−3 (catalytic) and PNKP-Nick-NCP−5 (catalytic) complexes are colored gray and blue, respectively. **(C)** Structural overlay of the PNKP catalytic domains bound to the nucleosomal DNA in the PNKP-Nick-NCP−4 (catalytic) and PNKP-Nick-NCP−5 (catalytic) complexes. The PNKP-Nick-NCP−4 (catalytic) and PNKP-Nick-NCP−5 (catalytic) complexes are colored gray and blue, respectively.

**Fig. S16: PNKP-Nick-NCP–3 (catalytic-FHA) map and model quality assessment**

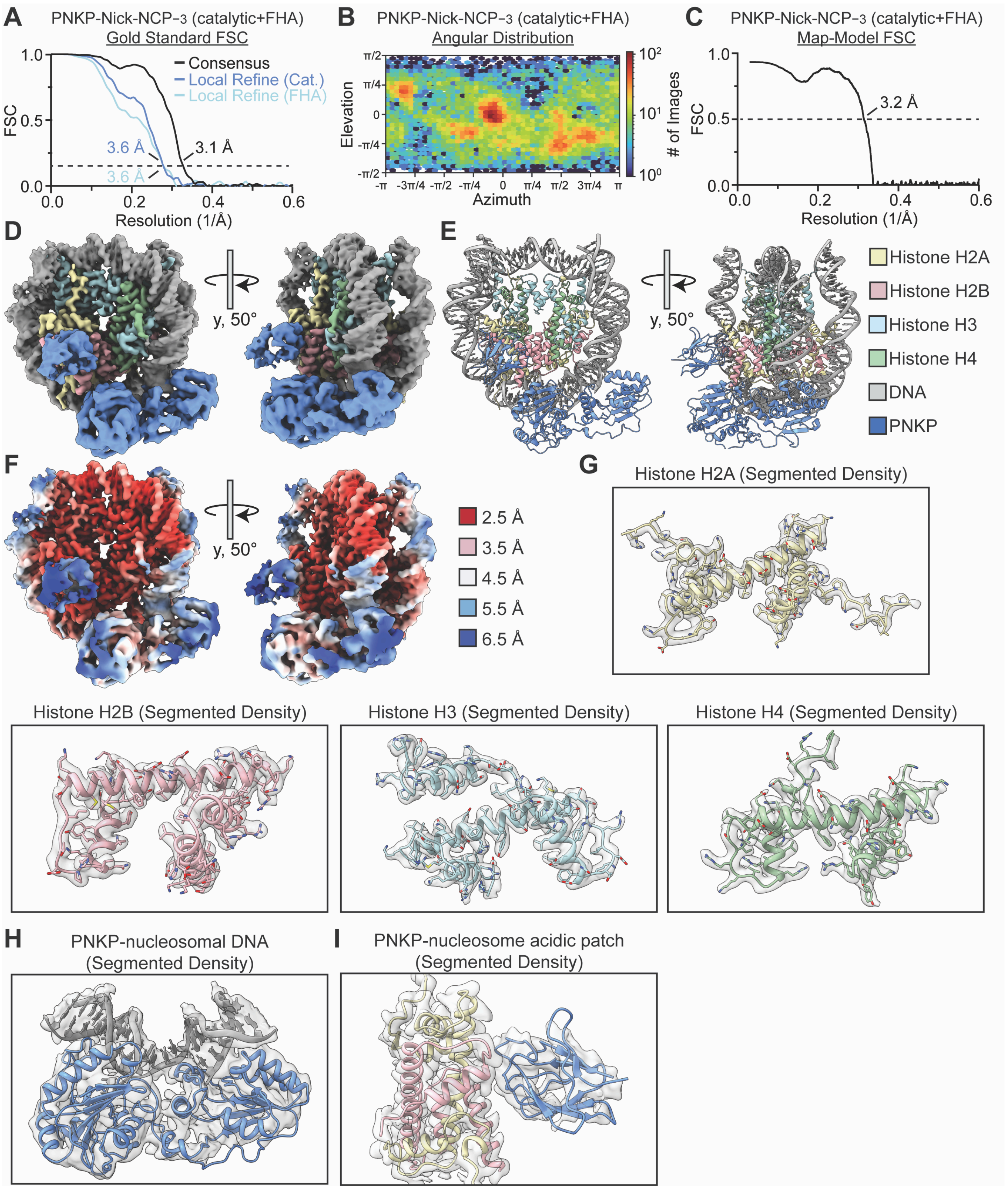

**Fig. S16: PNKP-Nick-NCP$_{-3}$ (catalytic-FHA) map and model quality assessment**

**(A**) Gold-standard Fourier shell correlation (GS-FSC) for the PNKP-Nick-NCP$_{-3}$ (catalytic-FHA) consensus cryo-EM map (solid black line), the PNKP/nucleosomal DNA focus cryo-EM map (solid blue line), and the PNKP FHA focus cryo-EM map (solid light blue line). The dashed line corresponds to GS-FSC - 0.143. **(B)** Angular distribution heatmap for the PNKP-Nick-NCP$_{-3}$ (catalytic-FHA) cryo-EM map**. (C)** Map-to-model FSC for the PNKP-Nick-NCP$_{-3}$ (catalytic-FHA) model and composite cryo-EM map. The dashed line corresponds to FSC - 0.5. **(D)** The final 3.1 Å PNKP-Nick-NCP$_{-3}$ (catalytic-FHA) composite cryo-EM map shown in two different orientations. **(E)** The final PNKP-Nick-NCP$_{-3}$ (catalytic-FHA) model shown in two different orientations. **(F)** Local resolution estimation for the PNKP-Nick-NCP$_{-3}$ (catalytic-FHA) composite cryo-EM map shown in two different orientations. **(G)** Representative segmented densities for histones H2A, H2B, H3, and H4 from the PNKP-Nick-NCP$_{-3}$ (catalytic-FHA) cryo-EM map. The representative segmented densities from the cryo-EM map are shown as transparent gray surfaces. **(H)** Representative segmented densities for the PNKP catalytic domains and the surrounding nucleosomal DNA from the PNKP-Nick-NCP$_{-3}$ (catalytic-FHA) cryo-EM map. The representative segmented densities from the cryo-EM map are shown as transparent gray surfaces. **(I)** Representative segmented densities for the PNKP FHA domain and the H2A/H2B dimer from the PNKP-Nick-NCP$_{-3}$ (catalytic-FHA) cryo-EM map. The representative segmented densities from the cryo-EM map are shown as transparent gray surfaces.

**Fig. S17: PNKP-Nick-NCP-4 (catalytic-FHA) map and model quality assessment**

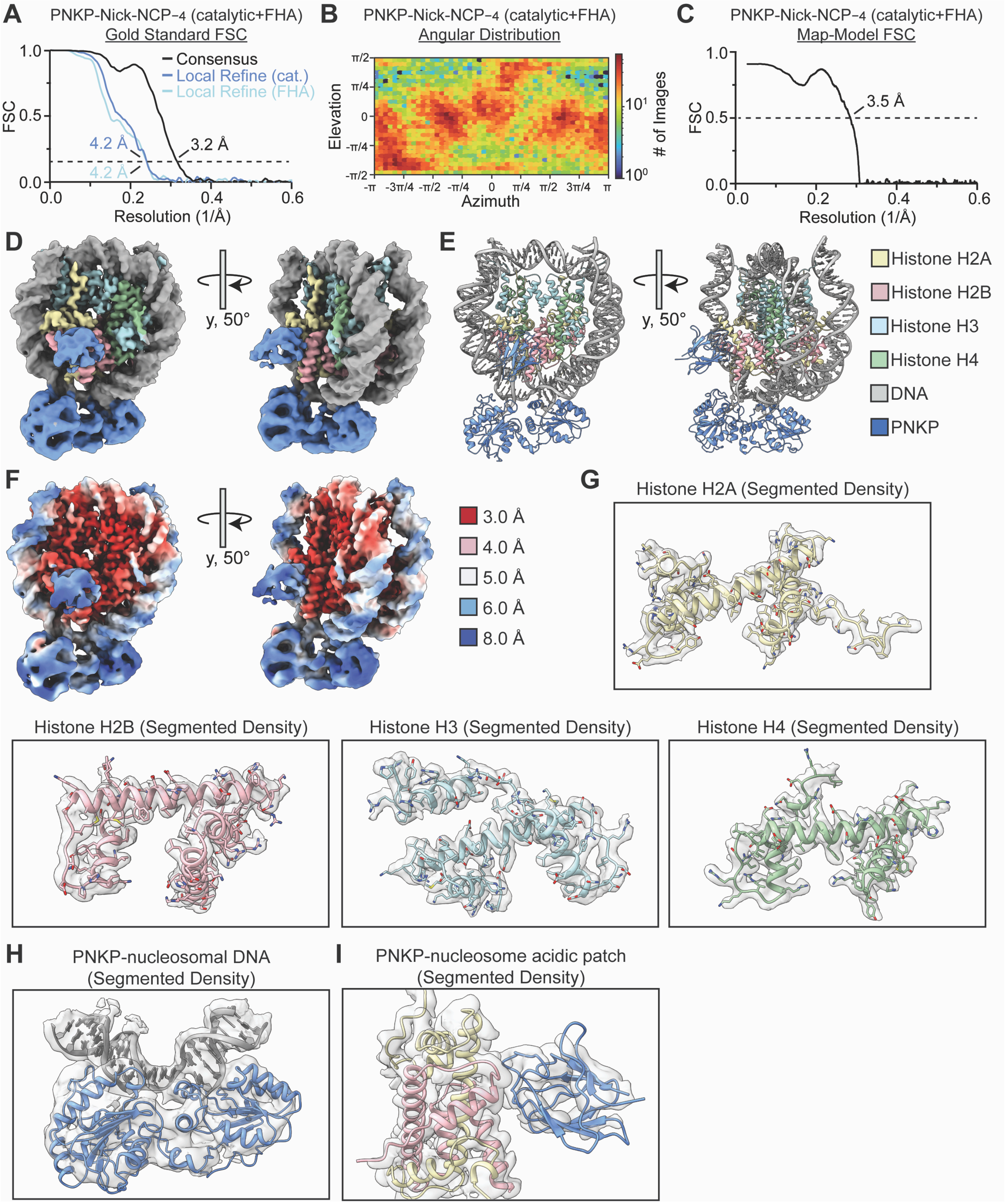

**Fig. S17: PNKP-Nick-NCP$_{-4}$ (catalytic-FHA) map and model quality assessment**

(**A**) Gold-standard Fourier shell correlation (GS-FSC) for the PNKP-Nick-NCP$_{-4}$ (catalytic-FHA) consensus cryo-EM map (solid black line), the PNKP/nucleosomal DNA focus cryo-EM map (solid blue line), and the PNKP FHA focus cryo-EM map (solid light blue line). The dashed line corresponds to GS-FSC - 0.143. **(B)** Angular distribution heatmap for the PNKP-Nick-NCP$_{-4}$ (catalytic-FHA) cryo-EM map**.** **(C)** Map-to-model FSC for the PNKP-Nick-NCP$_{-4}$ (catalytic-FHA) model and composite cryo-EM map. The dashed line corresponds to FSC - 0.5. **(D)** The final 3.2 Å PNKP-Nick-NCP$_{-4}$ (catalytic-FHA) composite cryo-EM map shown in two different orientations. **(E)** The final PNKP-Nick-NCP$_{-4}$ (catalytic-FHA) model shown in two different orientations. **(F)** Local resolution estimation for the PNKP-Nick-NCP$_{-4}$ (catalytic-FHA) composite cryo-EM map shown in two different orientations. **(G)** Representative segmented densities for histones H2A, H2B, H3, and H4 from the PNKP-Nick-NCP$_{-4}$ (catalytic-FHA) cryo-EM map. The representative segmented densities from the cryo-EM map are shown as transparent gray surfaces. **(H)** Representative segmented densities for the PNKP catalytic domains and the surrounding nucleosomal DNA from the PNKP-Nick-NCP$_{-4}$ (catalytic-FHA) cryo-EM map. The representative segmented densities from the cryo-EM map are shown as transparent gray surfaces. **(I)** Representative segmented densities for the PNKP FHA domain and the H2A/H2B dimer from the PNKP-Nick-NCP$_{-4}$ (catalytic-FHA) cryo-EM map. The representative segmented densities from the cryo-EM map are shown as transparent gray surfaces.

**Fig. S18: PNKP-Nick-NCP−5 (catalytic-FHA) map and model quality assessment**

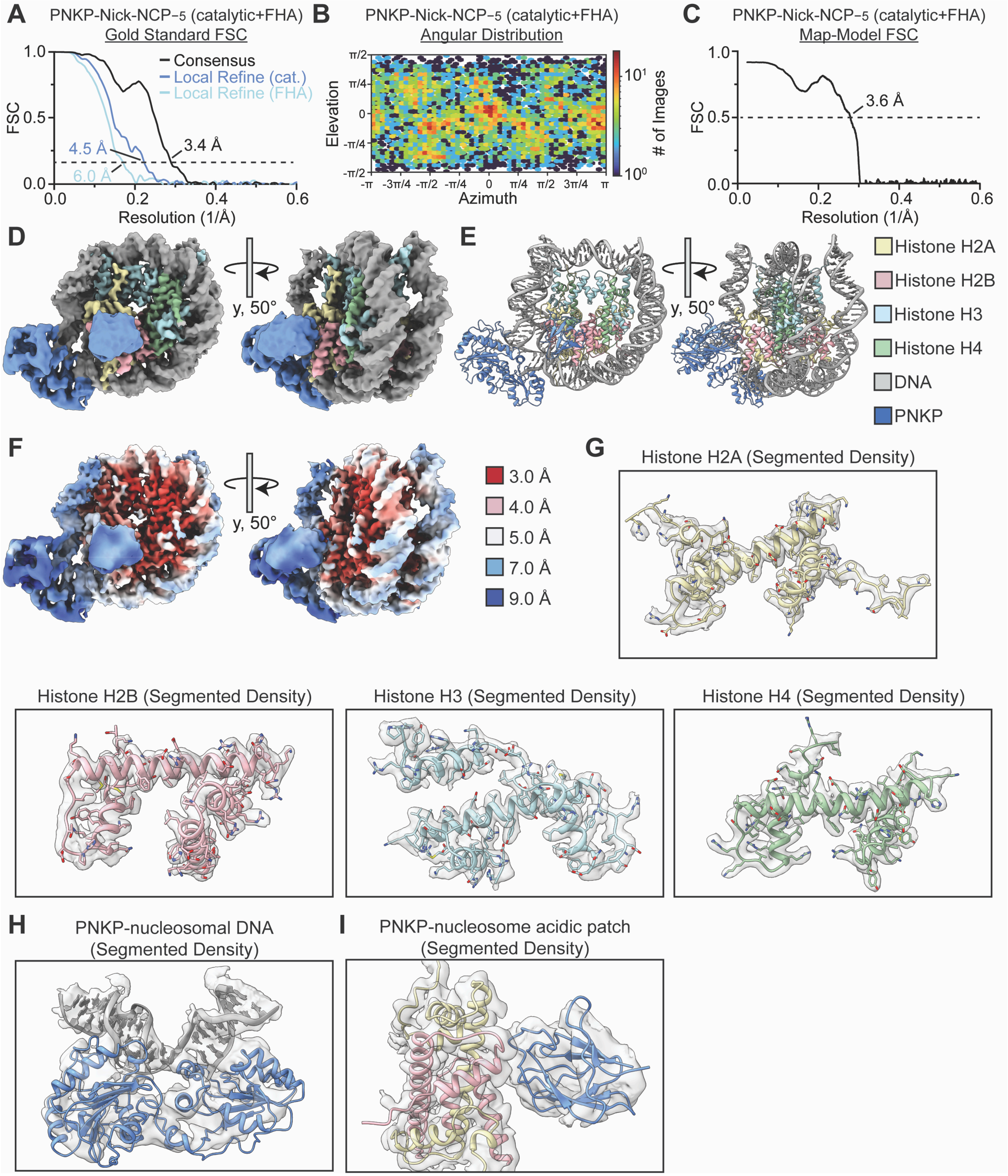

**Fig. S18: PNKP-Nick-NCP$_{-5}$ (catalytic-FHA) map and model quality assessment**

(**A**) Gold-standard Fourier shell correlation (GS-FSC) for the PNKP-Nick-NCP$_{-5}$ (catalytic-FHA) consensus cryo-EM map (solid black line), the PNKP/nucleosomal DNA focus cryo-EM map (solid blue line), and the PNKP FHA focus cryo-EM map (solid light blue line). The dashed line corresponds to GS-FSC - 0.143. **(B)** Angular distribution heatmap for the PNKP-Nick-NCP$_{-5}$ (catalytic-FHA) cryo-EM map**.** **(C)** Map-to-model FSC for the PNKP-Nick-NCP$_{-5}$ (catalytic-FHA) model and composite cryo-EM map. The dashed line corresponds to FSC - 0.5. **(D)** The final 3.4 Å PNKP-Nick-NCP$_{-5}$ (catalytic-FHA) composite cryo-EM map shown in two different orientations. **(E)** The final PNKP-Nick-NCP$_{-5}$ (catalytic-FHA) model shown in two different orientations. **(F)** Local resolution estimation for the PNKP-Nick-NCP$_{-5}$ (catalytic-FHA) composite cryo-EM map shown in two different orientations. **(G)** Representative segmented densities for histones H2A, H2B, H3, and H4 from the PNKP-Nick-NCP$_{-5}$ (catalytic-FHA) cryo-EM map. The representative segmented densities from the cryo-EM map are shown as transparent gray surfaces. **(H)** Representative segmented densities for the PNKP catalytic domains and the surrounding nucleosomal DNA from the PNKP-Nick-NCP$_{-5}$ (catalytic-FHA) cryo-EM map. The representative segmented densities from the cryo-EM map are shown as transparent gray surfaces. **(I)** Representative segmented densities for the PNKP FHA domain and the H2A/H2B dimer from the PNKP-Nick-NCP$_{-5}$ (catalytic-FHA) cryo-EM map. The representative segmented densities from the cryo-EM map are shown as transparent gray surfaces.

**Fig. S19: PNKP-Nick-NCP−3 (FHA) map and model quality assessment**

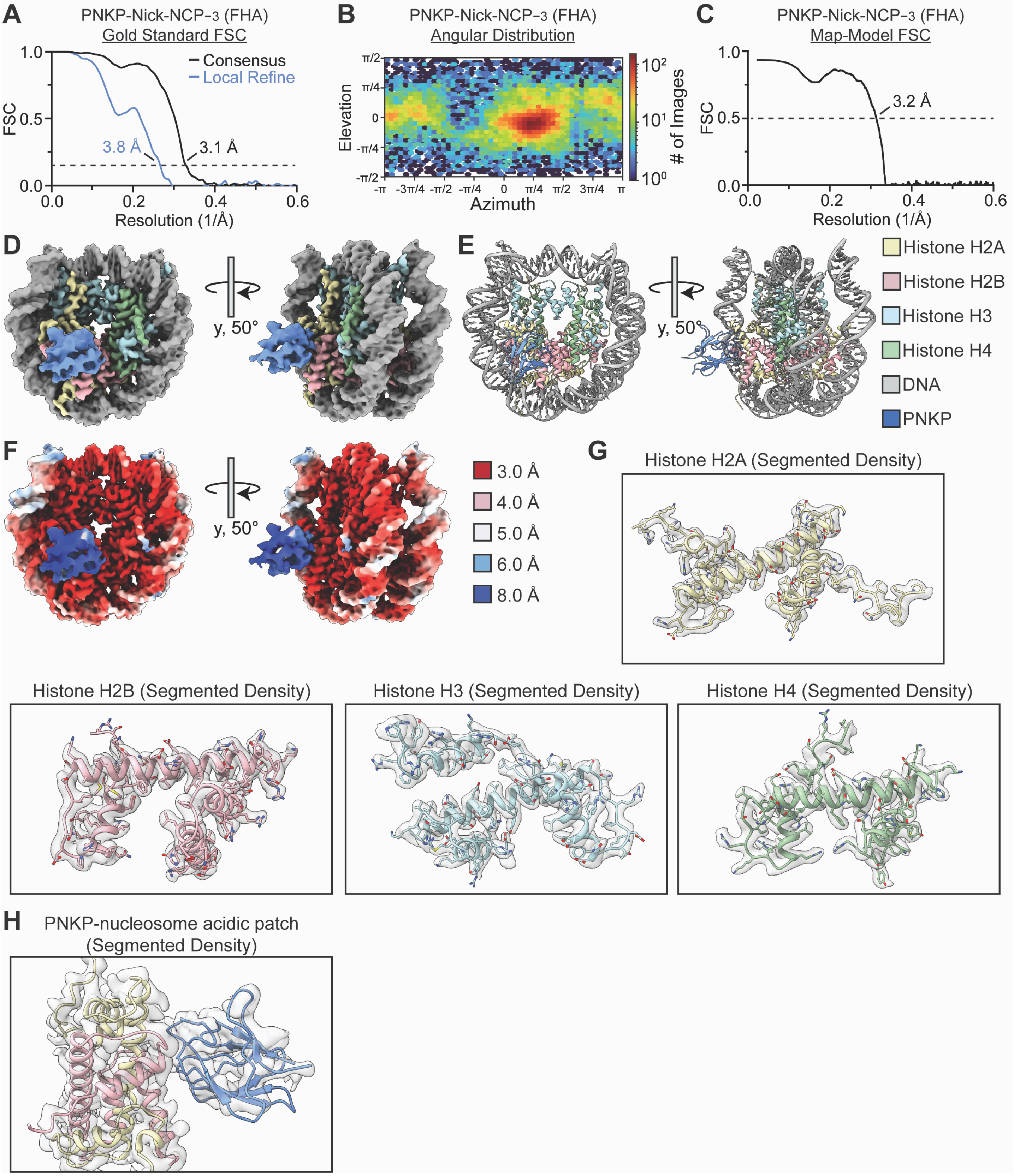

**Fig. S19: PNKP-Nick-NCP$_{-3}$ (FHA) map and model quality assessment**

**(A**) Gold-standard Fourier shell correlation (GS-FSC) for the PNKP-Nick-NCP$_{-3}$ (FHA) consensus cryo-EM map (solid black line) and the PNKP FHA focus cryo-EM map (solid blue line). The dashed line corresponds to GS-FSC - 0.143. **(B)** Angular distribution heatmap for the PNKP-Nick-NCP$_{-3}$ (FHA) cryo-EM map**.** **(C)** Map-to-model FSC for the PNKP-Nick-NCP$_{-3}$ (FHA) model and composite cryo-EM map. The dashed line corresponds to FSC - 0.5. **(D)** The final 3.1 Å PNKP-Nick-NCP$_{-3}$ (FHA) composite cryo-EM map shown in two different orientations. **(E)** The final PNKP-Nick-NCP$_{-3}$ (FHA) model shown in two different orientations. **(F)** Local resolution estimation for the PNKP-Nick-NCP$_{-3}$ (FHA) composite cryo-EM map shown in two different orientations. **(G)** Representative segmented densities for histones H2A, H2B, H3, and H4 from the PNKP-Nick-NCP$_{-3}$ (FHA) cryo-EM map. The representative segmented densities from the cryo-EM map are shown as transparent gray surfaces. **(H)** Representative segmented densities for the PNKP FHA domain and the H2A/H2B dimer from the PNKP-Nick-NCP$_{-3}$ (FHA) cryo-EM map. The representative segmented densities from the cryo-EM map are shown as transparent gray surfaces.

**Fig. S20: PNKP-Nick-NCP−4 (FHA) map and model quality assessment**

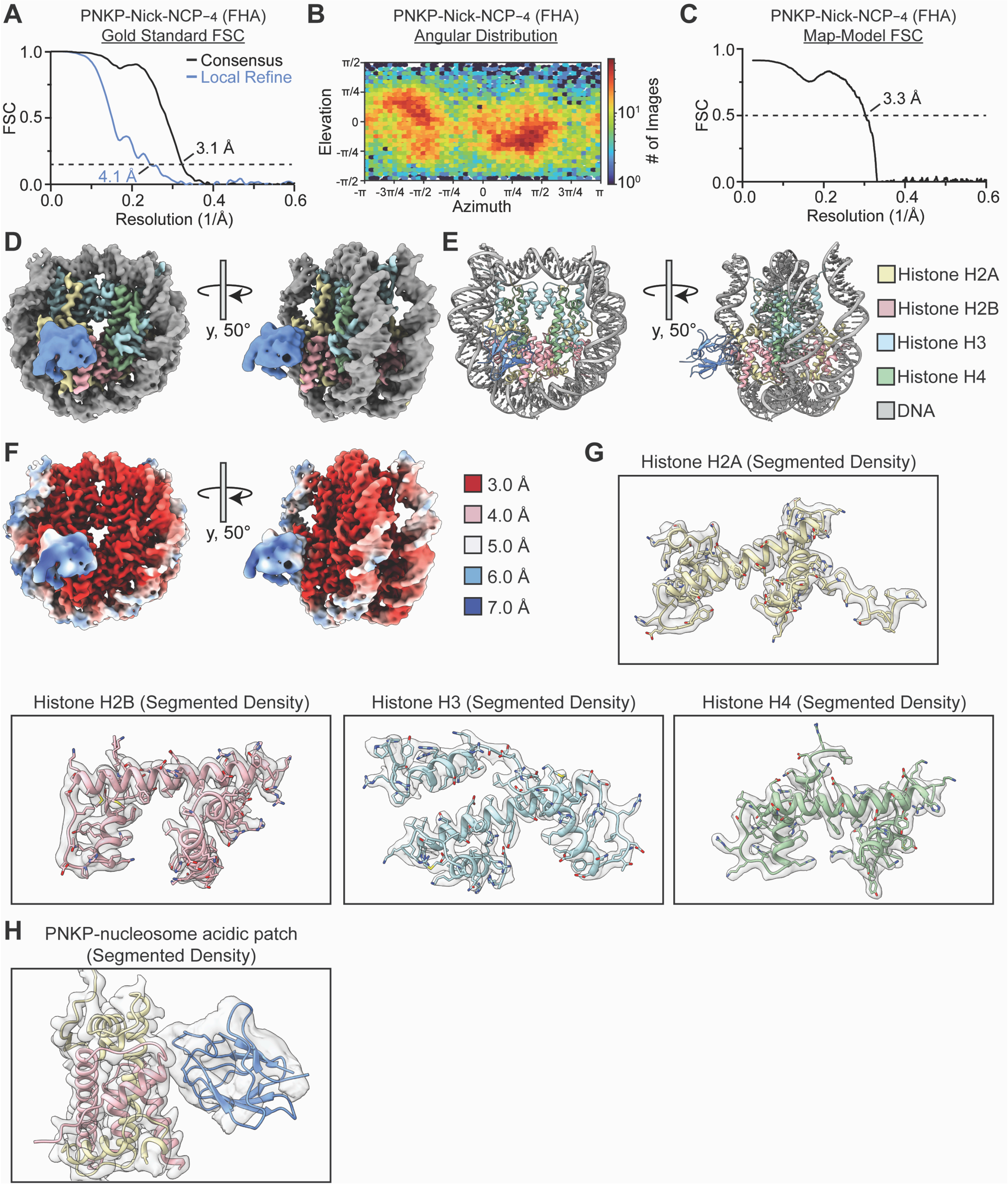

**Fig. S20: PNKP-Nick-NCP$_{-4}$ (FHA) map and model quality assessment**

**(A**) Gold-standard Fourier shell correlation (GS-FSC) for the PNKP-Nick-NCP$_{-4}$ (FHA) consensus cryo-EM map (solid black line) and the PNKP FHA focus cryo-EM map (solid blue line). The dashed line corresponds to GS-FSC - 0.143. **(B)** Angular distribution heatmap for the PNKP-Nick-NCP$_{-4}$ (FHA) cryo-EM map**.** **(C)** Map-to-model FSC for the PNKP-Nick-NCP$_{-4}$ (FHA) model and composite cryo-EM map. The dashed line corresponds to FSC - 0.5. **(D)** The final 3.1 Å PNKP-Nick-NCP$_{-4}$ (FHA) composite cryo-EM map shown in two different orientations. **(E)** The final PNKP-Nick-NCP$_{-4}$ (FHA) model shown in two different orientations. **(F)** Local resolution estimation for the PNKP-Nick-NCP$_{-4}$ (FHA) composite cryo-EM map shown in two different orientations. **(G)** Representative segmented densities for histones H2A, H2B, H3, and H4 from the PNKP-Nick-NCP$_{-4}$ (FHA) cryo-EM map. The representative segmented densities from the cryo-EM map are shown as transparent gray surfaces. **(H)** Representative segmented densities for the PNKP FHA domain and the H2A/H2B dimer from the PNKP-Nick-NCP$_{-4}$ (FHA) cryo-EM map. The representative segmented densities from the cryo-EM map are shown as transparent gray surfaces.

**Fig. S21: PNKP-Nick-NCP−5 (FHA) map and model quality assessment**

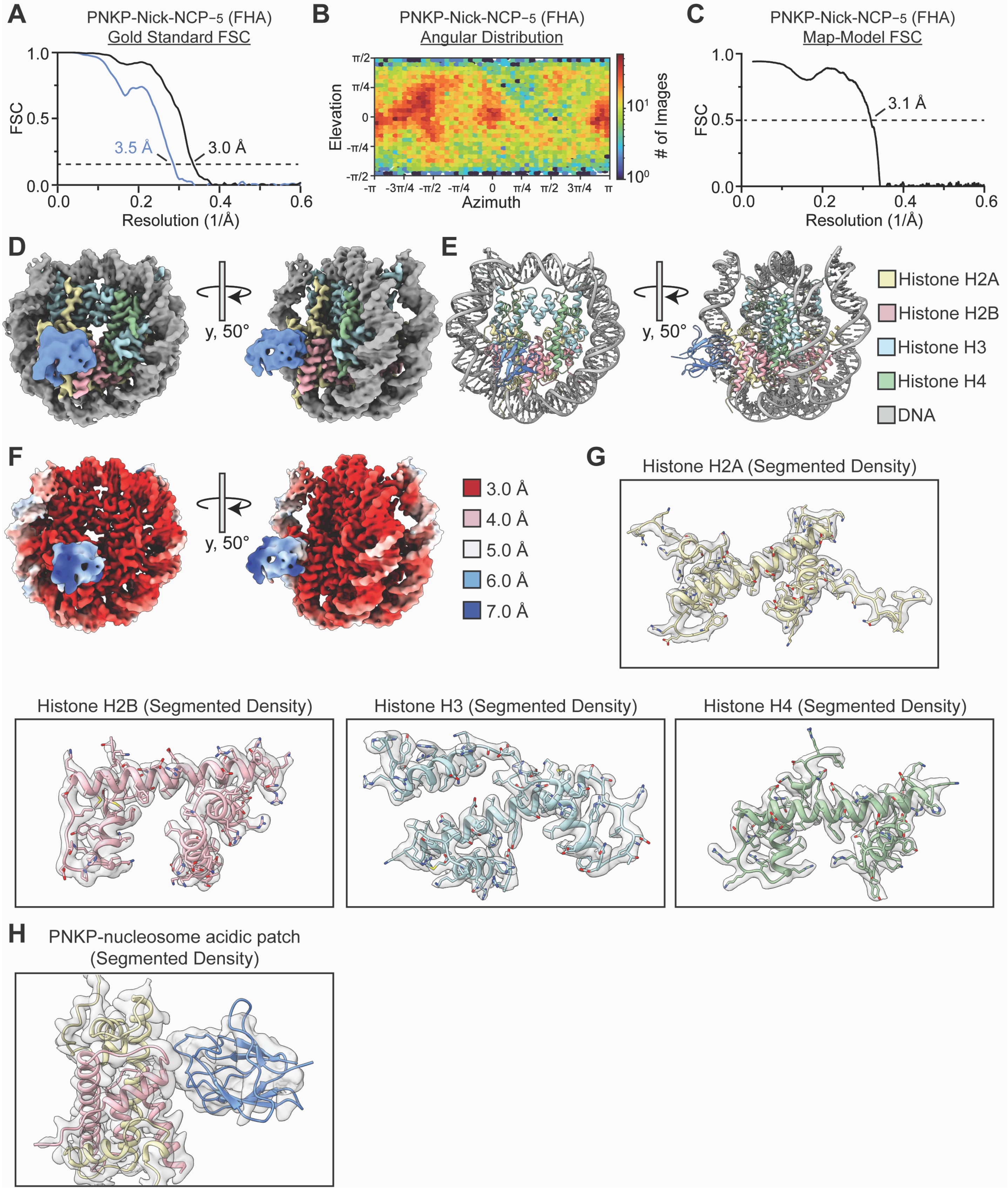

**Fig. S21: PNKP-Nick-NCP$_{-5}$ (FHA) map and model quality assessment**

**(A**) Gold-standard Fourier shell correlation (GS-FSC) for the PNKP-Nick-NCP$_{-5}$ (FHA) consensus cryo-EM map (solid black line) and the PNKP FHA focus cryo-EM map (solid blue line). The dashed line corresponds to GS-FSC - 0.143. **(B)** Angular distribution heatmap for the PNKP-Nick-NCP$_{-5}$ (FHA) cryo-EM map**. (C)** Map-to-model FSC for the PNKP-Nick-NCP$_{-5}$ (FHA) model and composite cryo-EM map. The dashed line corresponds to FSC - 0.5. **(D)** The final 3.0 Å PNKP-Nick-NCP$_{-5}$ (FHA) composite cryo-EM map shown in two different orientations. **(E)** The final PNKP-Nick-NCP$_{-5}$ (FHA) model shown in two different orientations. **(F)** Local resolution estimation for the PNKP-Nick-NCP$_{-5}$ (FHA) composite cryo-EM map shown in two different orientations. **(G)** Representative segmented densities for histones H2A, H2B, H3, and H4 from the PNKP-Nick-NCP$_{-5}$ (FHA) cryo-EM map. The representative segmented densities from the cryo-EM map are shown as transparent gray surfaces. **(H)** Representative segmented densities for the PNKP FHA domain and the H2A/H2B dimer from the PNKP-Nick-NCP$_{-5}$ (FHA) cryo-EM map. The representative segmented densities from the cryo-EM map are shown as transparent gray surfaces.

**Fig. S22: The PNKP FHA domain engages the nucleosome acidic patch with the catalytic domains disengaged from the non-ligatable nick**

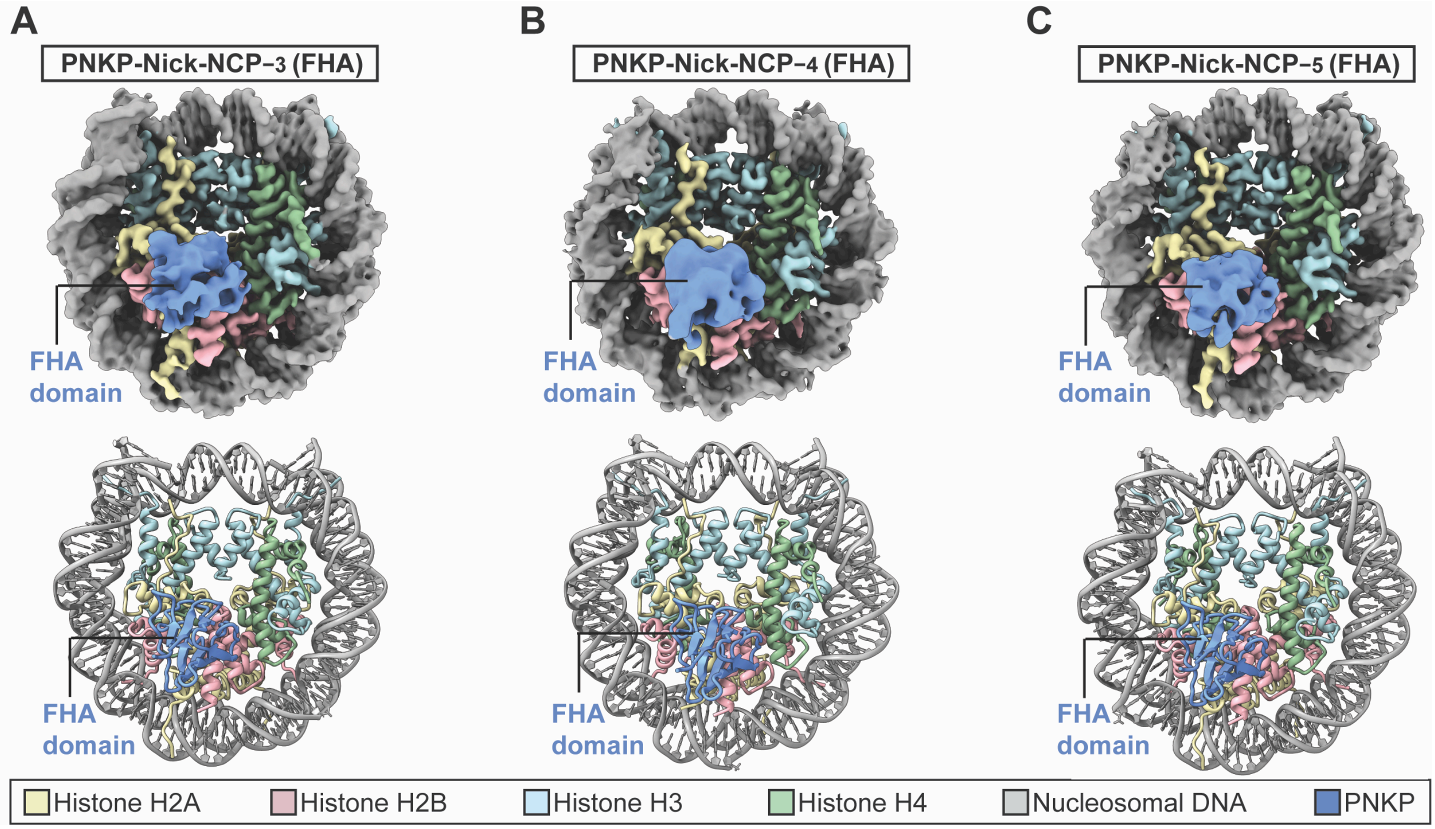

**Fig. S22: The PNKP FHA domain engages the nucleosome acidic patch with the catalytic domains disengaged from the non-ligatable nick**

**(A)** Composite cryo-EM map and model of the PNKP-Nick-NCP$_{-3}$ (FHA) complex**.** **(B)** Composite cryo-EM map and model of the PNKP-Nick-NCP$_{-4}$ (FHA) complex**.** **(C)** Composite cryo-EM map and model of the PNKP-Nick-NCP$_{-5}$ (FHA) complex**.**

**Fig. S23: The PNKP FHA domain engages the nucleosome acidic patch**

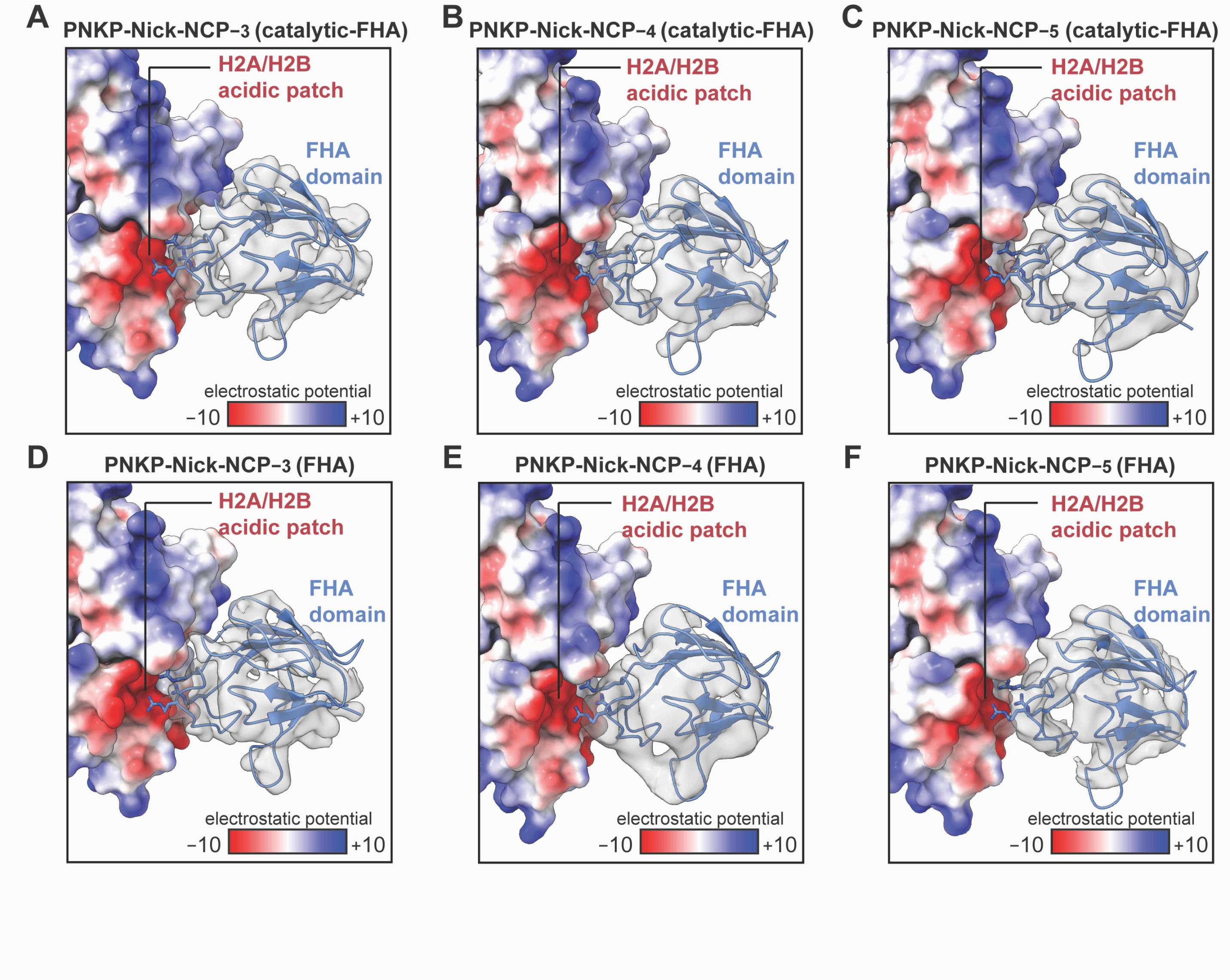

**Fig. S23: The PNKP FHA domain engages the nucleosome acidic patch**

**(A)** Focused view of the PNKP-Nick-NCP−3 (catalytic-FHA) complex highlighting the FHA-nucleosome acidic patch interaction. **(B)** Focused view of the PNKP-Nick-NCP−4 (catalytic-FHA) complex highlighting the FHA-nucleosome acidic patch interaction. **(C)** Focused view of the PNKP-Nick-NCP−5 (catalytic-FHA) complex highlighting the FHA-nucleosome acidic patch interaction. **(D)** Focused view of the PNKP-Nick-NCP−3 (FHA) complex highlighting the FHA-nucleosome acidic patch interaction. **(E)** Focused view of the PNKP-Nick-NCP−4 (FHA) complex highlighting the FHA-nucleosome acidic patch interaction. **(F)** Focused view of the PNKP-Nick-NCP−3 (FHA) complex highlighting the FHA-nucleosome acidic patch interaction. In **A-F**, segmented cryo-EM density for the FHA is shown as a transparent gray surface, FHA residues R35, R44, and R48 are shown as sticks, H2A/H2B are depicted as a surface representation with electrostatic surface potential.

**Fig. S24: Structural comparison of the PNKP-Nick-NCP (catalytic-FHA) and PNKP-Nick-NCP (FHA) complexes at SHL−3, SHL−4, and SHL−5**

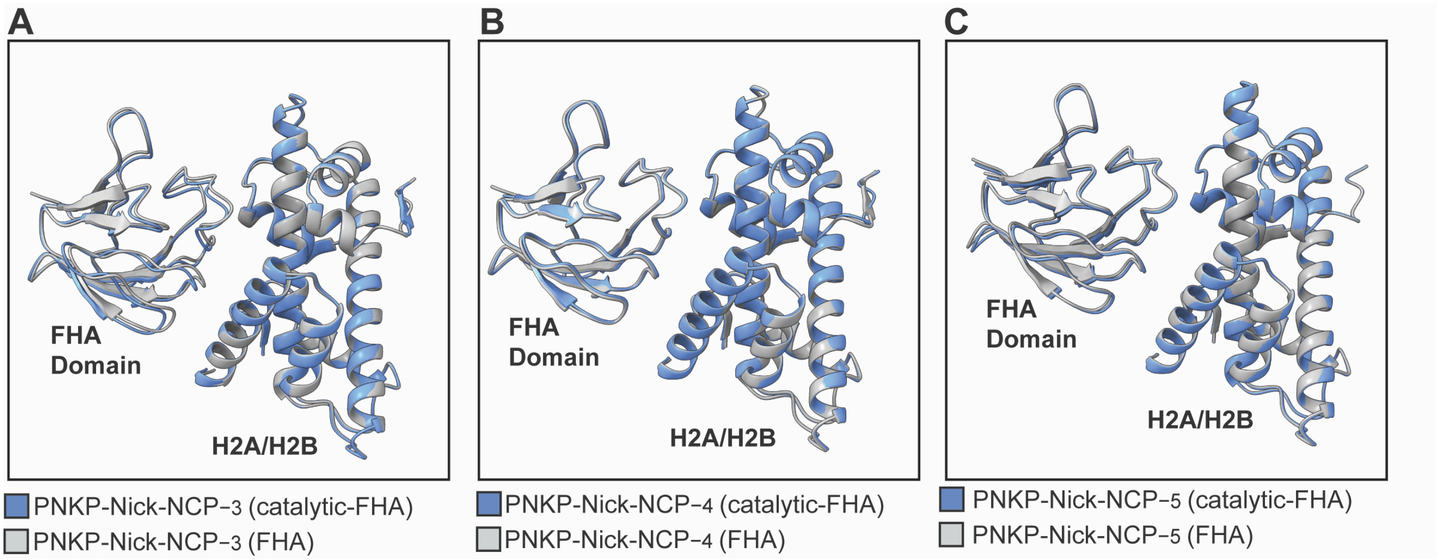

**Fig. S24: Structural comparison of the PNKP-Nick-NCP (catalytic-FHA) and PNKP-Nick-NCP (FHA) complexes at SHL$_{-3}$, SHL$_{-4}$, and SHL$_{-5}$**

**(A**) Structural overlay of the PNKP FHA domain bound to the nucleosome acidic patch in the PNKP-Nick-NCP$_{-3}$ (catalytic-FHA) and PNKP-Nick-NCP$_{-3}$ (FHA) complexes. The PNKP-Nick-NCP$_{-3}$ (catalytic-FHA) and PNKP-Nick-NCP$_{-3}$ (FHA) complexes are colored blue and gray, respectively. **(B)** Structural overlay of the PNKP FHA domain bound to the nucleosome acidic patch in the PNKP-Nick-NCP$_{-4}$ (catalytic-FHA) and PNKP-Nick-NCP$_{-4}$ (FHA) complexes. The PNKP-Nick-NCP$_{-4}$ (catalytic-FHA) and PNKP-Nick-NCP$_{-4}$ (FHA) complexes are colored blue and gray, respectively. **(C)** Structural overlay of the PNKP FHA domain bound to the nucleosome acidic patch in the PNKP-Nick-NCP$_{-5}$ (catalytic-FHA) and PNKP-Nick-NCP$_{-5}$ (FHA) complexes. The PNKP-Nick-NCP$_{-5}$ (catalytic-FHA) and PNKP-Nick-NCP$_{-5}$ (FHA) complexes are colored blue and gray, respectively.

**Fig. S25: Structural comparison of the PNKP-Nick-NCP (catalytic) and PNKP-Nick-NCP (catalytic-FHA) complexes at SHL−3, SHL−4, and SHL−5**

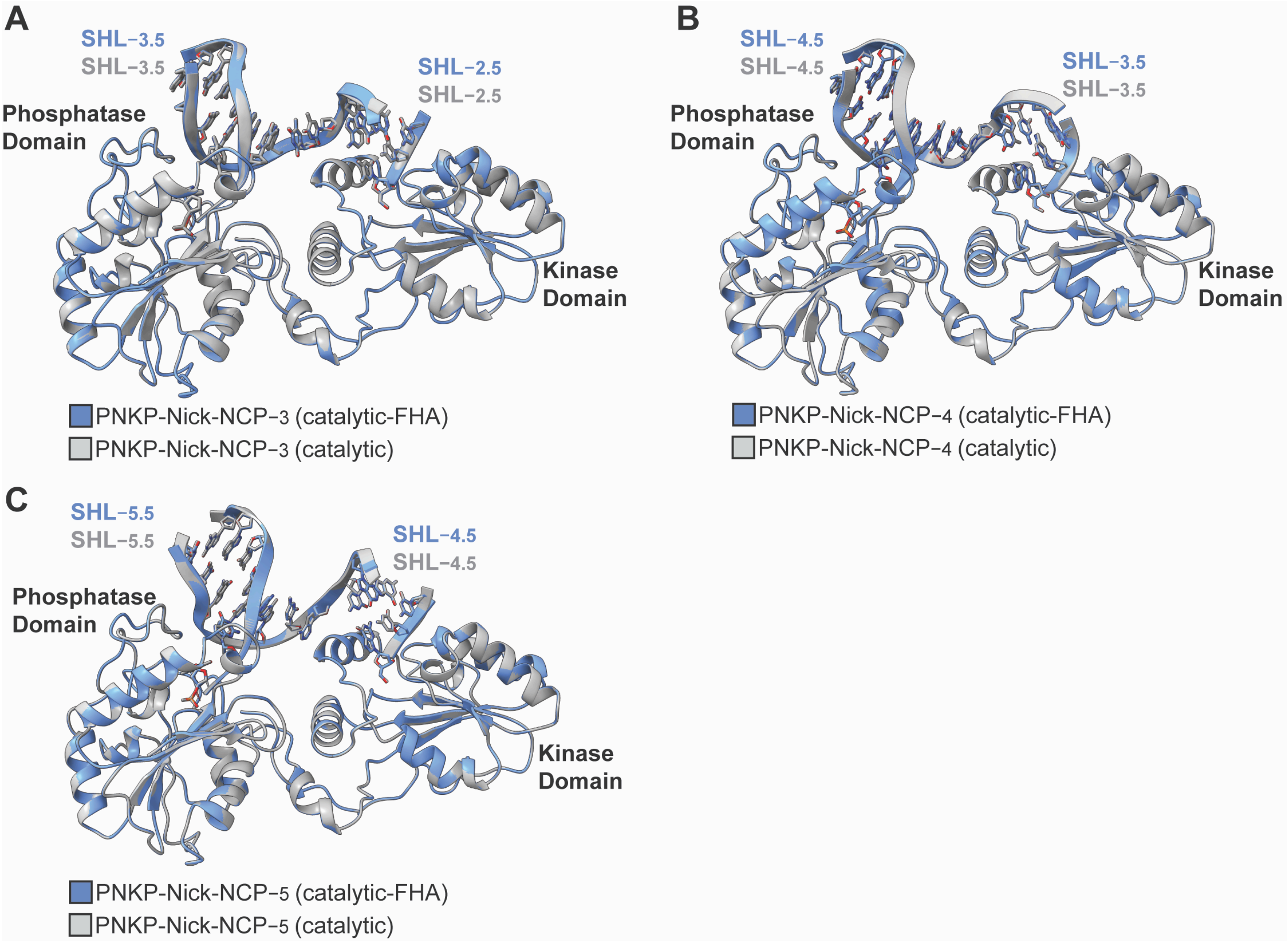

**Fig. S25: Structural comparison of the PNKP-Nick-NCP (catalytic) and PNKP-Nick-NCP (catalytic-FHA) complexes at SHL−3, SHL−4, and SHL−5**

**(A**) Structural overlay of the PNKP catalytic domains bound to the nucleosomal DNA in the PNKP-Nick-NCP−3 (catalytic) and PNKP-Nick-NCP−3 (catalytic-FHA) complexes. The PNKP-Nick-NCP−3 (catalytic) and PNKP-Nick-NCP−3 (catalytic-FHA) complexes are colored gray and blue, respectively. **(B)** Structural overlay of the PNKP catalytic domains bound to the nucleosomal DNA in the PNKP-Nick-NCP−4 (catalytic) and PNKP-Nick-NCP−4 (catalytic-FHA) complexes. The PNKP-Nick-NCP−4 (catalytic) and PNKP-Nick-NCP−4 (catalytic-FHA) complexes are colored gray and blue, respectively. **(C)** Structural overlay of the PNKP catalytic domains bound to the nucleosomal DNA in the PNKP-Nick-NCP−5 (catalytic) and PNKP-Nick-NCP−5 (catalytic-FHA) complexes. The PNKP-Nick-NCP−5 (catalytic) and PNKP-Nick-NCP−5 (catalytic-FHA) complexes are colored gray and blue, respectively.

**Fig. S26: Kinetic analysis of end processing in the nucleosome by the PNKP 3RA mutant**

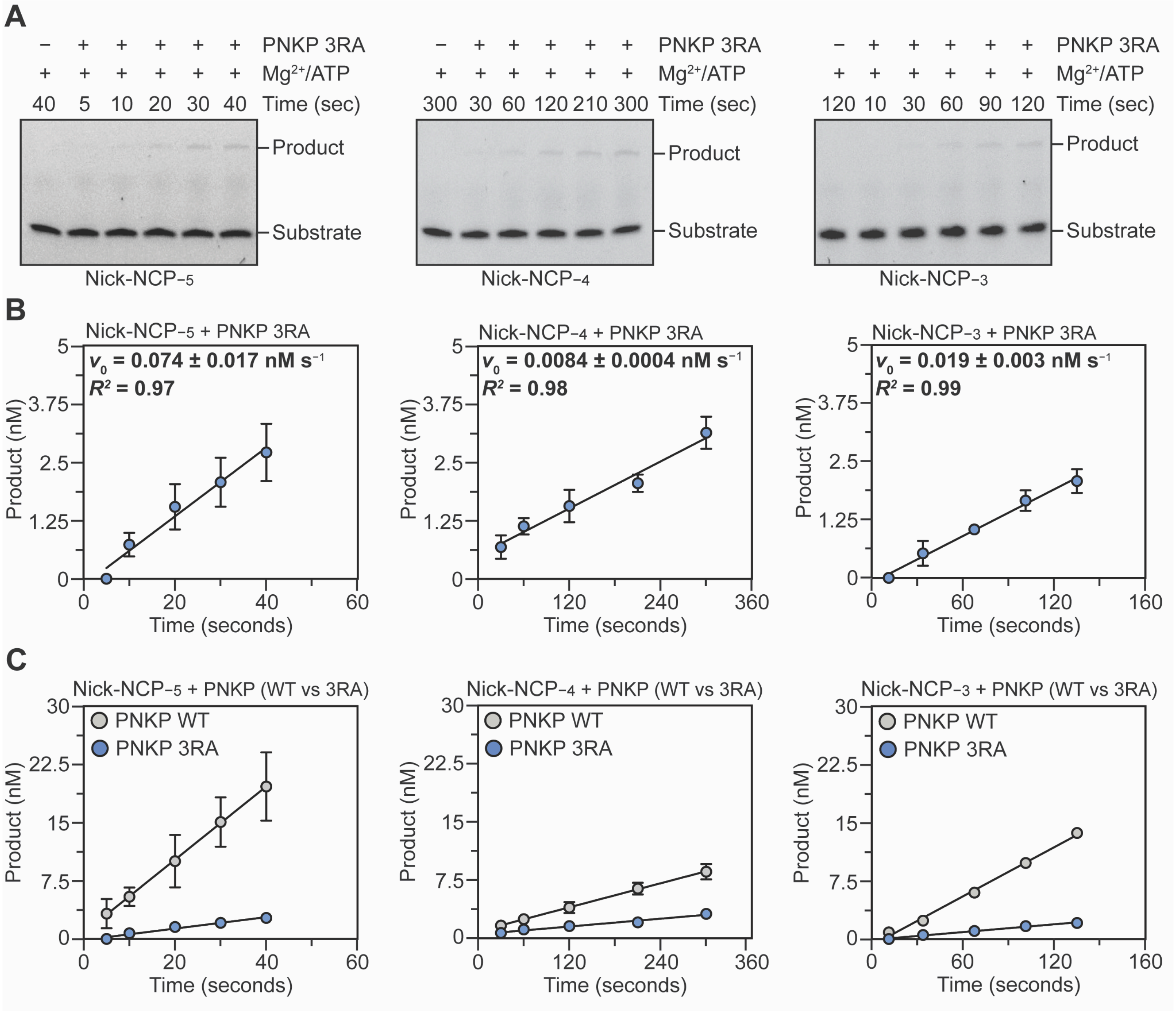

**Fig. S26: Kinetic analysis of end processing in the nucleosome by the PNKP 3RA mutant**

**(A)** Representative denaturing Urea-PAGE gels from the PNKP 3RA enzyme kinetics on Nick-NCP$_{-5}$, Nick-NCP$_{-4}$, and Nick-NCP$_{-3}$. The substrate and product were detected using the 6-FAM label on the nucleosomal DNA. The gels are representative of three independent replicate experiments performed for PNKP 3RA and Nick-NCP$_{-5}$, Nick-NCP$_{-4}$, and Nick-NCP$_{-3}$. **(B)** Progress curves for the PNKP 3RA enzyme kinetics on Nick-NCP$_{-5}$, Nick-NCP$_{-4}$, and Nick-NCP$_{-3}$. The data points represent the mean ± standard deviation from the three independent replicate experiments. The error bars are included for all experimental data points, but some error bars are smaller than the circles used to represent data points. The initial velocity ($v_0$, nM s$^{-1}$) obtained from the linear regression of product formation as a function of time is shown as an inset for each experiment and represents the mean ± standard deviation from the three independent replicate experiments. **(C)** Progress curves for the PNKP WT (gray) and PNKP 3RA (blue) enzyme kinetics on Nick-NCP$_{-5}$, Nick-NCP$_{-4}$, and Nick-NCP$_{-3}$ for direct comparison. The data points represent the mean ± standard deviation from the three independent replicate experiments. The error bars are included for all experimental data points, but some error bars are smaller than the circles used to represent data points. The initial velocity ($v_0$, nM s$^{-1}$) obtained from the linear regression of product formation as a function of time is shown as an inset for each experiment and represents the mean ± standard deviation from the three independent replicate experiments. The PNKP WT datapoints (gray) presented in **C** are the same data as presented in Fig. S3.

**Fig. S27: Validation of the DNA purification scheme after the PNKP kinetic time courses**

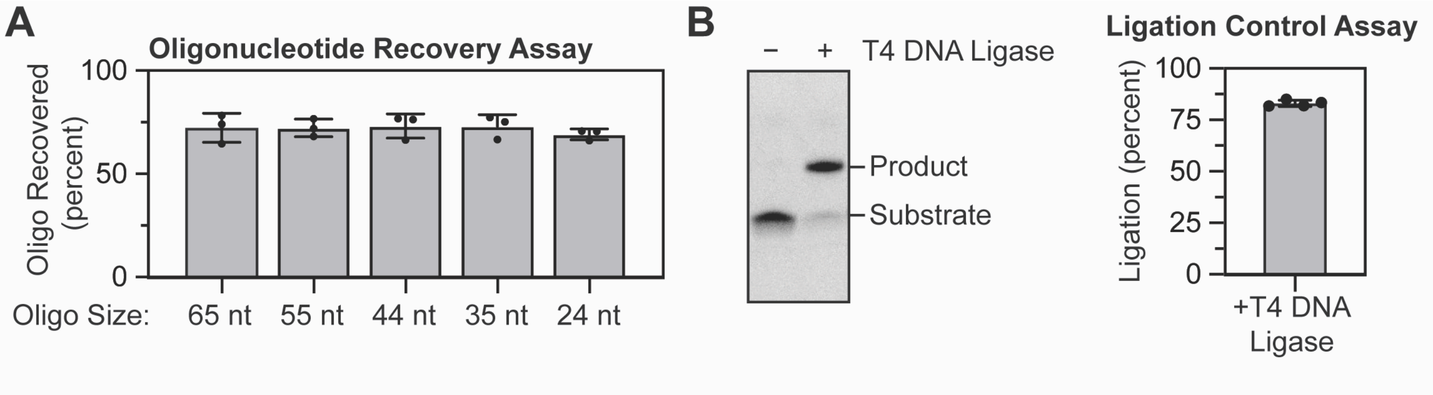

**Fig. S27: Validation of the DNA purification scheme after the PNKP kinetic time courses**

**(A)** Quantification of the oligonucleotide recovery assay using the Zymo Research oligonucleotide cleanup kit. The percent oligonucleotide recovered represents the mean ± standard deviation from three independent replicate experiments. **(B)** Representative denaturing Urea-PAGE gels from the T4 DNA ligase experiments showing robust ligation of a Nick-DNA containing a 5′-phosphate and 3′-OH following purification using the Zymo Research oligonucleotide cleanup kit (left). Quantification of the T4 DNA ligase experiments following purification of the Nick-DNA containing a 5′-phosphate and 3′-OH using the Zymo Research oligonucleotide cleanup kit. The percent of ligated DNA represents the mean ± standard deviation from four independent replicate experiments (right).

**Table S1: PNKP-Nick-NCP$_{-3}$ cryo-EM data collection and validation**

| **Data collection and processing** | | | | |
|---|---|---|---|---|
| **Dataset** | **Nick-NCP$_{-3}$** | | | |
| Magnification | 130,000x | | | |
| Voltage (kV) | 300 | | | |
| Electron exposure ($e^{-}/Å^{2}$) | 50 | | | |
| Defocus range (μm) | −1.2 to −1.8 | | | |
| Pixel size (Å) | 0.652 | | | |
| Symmetry imposed | C1 | | | |
| Initial particle images (no.) | 1,092,309 | | | |
| **Structure** | **Nick-NCP$_{-3}$** | **PNKP-Nick-NCP$_{-3}$ (catalytic)** | **PNKP-Nick-NCP$_{-3}$ (catalytic-FHA)** | **PNKP-Nick-NCP$_{-3}$ (FHA)** |
| Final particle images (no.) | 186,206 | 39,657 | 24,323 | 25,321 |
| Map resolution (Å) | 2.6 | 2.9 | 3.1 | 3.1 |
| FSC threshold | 0.143 | 0.143 | 0.143 | 0.143 |
| PDB accession | 37XL | 37XM | 37XO | 37XN |
| EMDB accession | EMD-78600 | EMD-78601<br>EMD-78612<br>EMD-78613 | EMD-78603<br>EMD-78614<br>EMD-78615<br>EMD-78616 | EMD-78602<br>EMD-78617<br>EMD-78618 |
| **Refinement** | | | | |
| Initial model used (PDB ID) | 10XZ | 10XZ, 3ZVN | 10XZ, 3ZVN, 2W3O | 10XZ, 2W3O |
| Model resolution (Å) | 2.8 | 3.1 | 3.2 | 3.2 |
| FSC threshold | 0.5 | 0.5 | 0.5 | 0.5 |
| **Model composition** | | | | |
| Nonhydrogen atoms | 12,013 | 14,883 | 15,651 | 12,759 |
| Protein residues | 755 | 1,128 | 1,231 | 855 |
| Nucleotide | 294 | 294 | 294 | 294 |
| **B factors ($Å^2$)** | | | | |
| Protein | 92.96 | 91.63 | 123.27 | 73.08 |
| Nucleotide | 135.46 | 141.88 | 154.21 | 123.01 |
| **r.m.s. deviations** | | | | |
| Bond Length (Å) (# > 4σ) | 0.007 (0) | 0.006 (0) | 0.004 (0) | 0.004 (1) |
| Bond Angles (°) (# > 4σ) | 0.829 (0) | 0.706 (1) | 0.612 (0) | 0.588 (0) |
| **Validation** | | | | |
| MolProbity score | 0.95 | 1.10 | 1.15 | 1.14 |
| Clashscore | 1.91 | 3.07 | 3.56 | 3.54 |
| Rotamer Outliers (%) | 0.32 | 0.33 | 0.40 | 0.28 |
| **Ramachandran plot** | | | | |
| Favored (%) | 98.11 | 98.02 | 98.51 | 98.09 |
| Allowed (%) | 1.89 | 1.98 | 1.49 | 1.91 |
| Disallowed (%) | 0.00 | 0.00 | 0.00 | 0.00 |

**Table S2: PNKP-Nick-NCP$_{-4}$ cryo-EM data collection and validation**

| **Data collection and processing** | | | | |
|---|---|---|---|---|
| **Dataset** | **Nick-NCP$_{-4}$** | | | |
| Magnification | 130,000x | | | |
| Voltage (kV) | 300 | | | |
| Electron exposure ($e^{-}/Å^{2}$) | 50 | | | |
| Defocus range (μm) | −1.2 to −1.8 | | | |
| Pixel size (Å) | 0.652 | | | |
| Symmetry imposed | C1 | | | |
| Initial particle images (no.) | 2,134,068 | | | |
| **Structure** | **Nick-NCP$_{-4}$** | **PNKP-Nick-NCP$_{-4}$ (catalytic)** | **PNKP-Nick-NCP$_{-4}$ (catalytic-FHA)** | **PNKP-Nick-NCP$_{-4}$ (FHA)** |
| Final particle images (no.) | 220,401 | 23,237 | 42,334 | 27,487 |
| Map resolution (Å) | 2.6 | 3.5 | 3.2 | 3.1 |
| FSC threshold | 0.143 | 0.143 | 0.143 | 0.143 |
| PDB accession | 37XP | 37XQ | 37XS | 37XT |
| EMDB accession | EMD-78604 | EMD-78605<br>EMD-78619<br>EMD-78620 | EMD-78606<br>EMD-78621<br>EMD-78623<br>EMD-78622 | EMD-78607<br>EMD-78624<br>EMD-78625 |
| **Refinement** | | | | |
| Initial model used (PDB ID) | 10XZ | 10XZ, 3ZVN | 10XZ, 3ZVN, 2W3O | 10XZ, 2W3O |
| Model resolution (Å) | 2.7 | 3.8 | 3.5 | 3.3 |
| FSC threshold | 0.5 | 0.5 | 0.5 | 0.5 |
| **Model composition** | | | | |
| Nonhydrogen atoms | 12,013 | 14,892 | 15,624 | 12,715 |
| Protein residues | 755 | 1,129 | 1,228 | 850 |
| Nucleotide | 294 | 294 | 294 | 294 |
| **B factors ($Å^{2}$)** | | | | |
| Protein | 93.89 | 183.03 | 203.14 | 80.79 |
| Nucleotide | 142.48 | 213.24 | 223.57 | 147.20 |
| **r.m.s. deviations** | | | | |
| Bond Length (Å) (# > 4σ) | 0.005 (0) | 0.004 (0) | 0.004 (0) | 0.003 (0) |
| Bond Angles (°) (# > 4σ) | 0.773 (0) | 0.582 (0) | 0.604 (0) | 0.609 (0) |
| **Validation** | | | | |
| MolProbity score | 0.89 | 1.11 | 1.17 | 1.00 |
| Clashscore | 1.53 | 3.18 | 3.78 | 2.21 |
| Rotamer Outliers (%) | 0.16 | 0.22 | 0.40 | 0.43 |
| **Ramachandran plot** | | | | |
| Favored (%) | 98.24 | 98.83 | 98.76 | 98.20 |
| Allowed (%) | 1.76 | 1.17 | 1.24 | 1.80 |
| Disallowed (%) | 0.00 | 0.00 | 0.00 | 0.00 |

**Table S3: PNKP-Nick-NCP$_{-5}$ cryo-EM data collection and validation**

| **Data collection and processing** | | | | |
|---|---|---|---|---|
| **Dataset** | **Nick-NCP$_{-5}$** | | | |
| Magnification | 130,000x | | | |
| Voltage (kV) | 300 | | | |
| Electron exposure ($e^{-}/Å^{2}$) | 50 | | | |
| Defocus range (μm) | −1.2 to −1.8 | | | |
| Pixel size (Å) | 0.652 | | | |
| Symmetry imposed | C1 | | | |
| Initial particle images (no.) | 1,613,194 | | | |
| **Structure** | **Nick-NCP$_{-5}$** | **PNKP-Nick-NCP$_{-5}$ (catalytic)** | **PNKP-Nick-NCP$_{-5}$ (catalytic-FHA)** | **PNKP-Nick-NCP$_{-5}$ (FHA)** |
| Final particle images (no.) | 116,841 | 12,103 | 7,817 | 33,821 |
| Map resolution (Å) | 2.8 | 3.2 | 3.4 | 3.0 |
| FSC threshold | 0.143 | 0.143 | 0.143 | 0.143 |
| PDB accession | 37XU | 37XV | 37XW | 37XX |
| EMDB accession | EMD-78608 | EMD-78609<br>EMD-78626<br>EMD-78627 | EMD-78610<br>EMD-78628<br>EMD-78629<br>EMD-78630 | EMD-78611<br>EMD-78631<br>EMD-78632 |
| **Refinement** | | | | |
| Initial model used (PDB ID) | 10XZ | 10XZ, 3ZVN | 10XZ, 3ZVN, 2W3O | 10XZ, 2W3O |
| Model resolution (Å) | 2.9 | 3.4 | 3.6 | 3.1 |
| FSC threshold | 0.5 | 0.5 | 0.5 | 0.5 |
| **Model composition** | | | | |
| Nonhydrogen atoms | 12,022 | 14,915 | 15,692 | 12,776 |
| Protein residues | 756 | 1,132 | 1,236 | 855 |
| Nucleotide | 294 | 294 | 294 | 294 |
| **B factors ($Å^2$)** | | | | |
| Protein | 87.54 | 135.91 | 136.92 | 81.92 |
| Nucleotide | 149.21 | 158.91 | 144.40 | 143.97 |
| **r.m.s. deviations** | | | | |
| Bond Length (Å) (# > 4σ) | 0.005 (0) | 0.004 (0) | 0.004 (0) | 0.006 (0) |
| Bond Angles (°) (# > 4σ) | 0.637 (0) | 0.602 (1) | 0.599 (0) | 0.629 (1) |
| **Validation** | | | | |
| MolProbity score | 0.82 | 0.90 | 0.87 | 1.01 |
| Clashscore | 1.14 | 1.45 | 1.36 | 2.29 |
| Rotamer Outliers (%) | 0.32 | 0.32 | 0.40 | 0.28 |
| **Ramachandran plot** | | | | |
| Favored (%) | 98.51 | 97.94 | 98.11 | 100.00 |
| Allowed (%) | 1.49 | 2.06 | 1.89 | 0.00 |
| Disallowed (%) | 0.00 | 0.00 | 0.00 | 0.00 |

**Table S4: Oligonucleotides for generating NCPs**

| Oligo | Sequence (5′ – 3′) |
|---|---|
| **ND-NCP** | |
| Oligo 1 | /5Phos/ATCGAGAATCCCGGTGCCGAGGCCGCTCAATTGGTCGTAGACAGCTCTAGCACCGCTTAAACGCACGTACGCGCTGTCCCCCGCGTTTTAACCGCCAAGGGGATTACTCCCTAGTCTCCAGGCACGTGTCAGATATATACATCCGAT/3OH/ |
| Oligo 2 | /6FAM/ATCGGATGTATATATCTGACACGTGCCTGGAGACTAGGGAGTAATCCCCTTGGCGGTTAAAACGCGGGGGACAGCGCGTACGTGCGTTTAAGCGGTGCTAGAGCTGTCTACGACCAATTGAGCGGCCTCGGCACCGGGATTCTCGAT/3OH/ |
| **Nick-NCP–5** | |
| Oligo 1 | /5Phos/ATCGAGAATCCCGGTGCCGAGGCCGCTCAATTGGTCGTAGACAGCTCTAGCACCGCTTAAACGCACGTACGCGCTGTCCCCCGCGTTTTAACCGCCAAGGGGATTACTCCCTAGTCTCCAGGCACGTGTCAGATATATACATCCGAT/3OH/ |
| Oligo 2 | /6FAM/ATCGGATGTATATATCTGACACGT/3Phos/ |
| Oligo 3 | /5OH/GCCTGGAGACTAGGGAGTAATCCCCTTGGCGGTTAAAACGCGGGGGACAGCGCGTACGTGCGTTTAAGCGGTGCTAGAGCTGTCTACGACCAATTGAGCGGCCTCGGCACCGGGATTCTCGAT/3OH/ |
| **Nick-NCP–4** | |
| Oligo 1 | /5Phos/ATCGAGAATCCCGGTGCCGAGGCCGCTCAATTGGTCGTAGACAGCTCTAGCACCGCTTAAACGCACGTACGCGCTGTCCCCCGCGTTTTAACCGCCAAGGGGATTACTCCCTAGTCTCCAGGCACGTGTCAGATATATACATCCGAT/3OH/ |
| Oligo 2 | /6FAM/ATCGGATGTATATATCTGACACGTGCCTGGAGACT/3Phos/ |
| Oligo 3 | /5OH/AGGGAGTAATCCCCTTGGCGGTTAAAACGCGGGGGACAGCGCGTACGTGCGTTTAAGCGGTGCTAGAGCTGTCTACGACCAATTGAGCGGCCTCGGCACCGGGATTCTCGAT/3OH/ |
| **Nick-NCP–3** | |
| Oligo 1 | /5Phos/ATCGAGAATCCCGGTGCCGAGGCCGCTCAATTGGTCGTAGACAGCTCTAGCACCGCTTAAACGCACGTACGCGCTGTCCCCCGCGTTTTAACCGCCAAGGGGATTACTCCCTAGTCTCCAGGCACGTGTCAGATATATACATCCGAT/3OH/ |
| Oligo 2 | /6FAM/ATCGGATGTATATATCTGACACGTGCCTGGAGACTAGGGAGTAA/3Phos/ |
| Oligo 3 | /5OH/TCCCCTTGGCGGTTAAAACGCGGGGGACAGCGCGTACGTGCGTTTAAGCGGTGCTAGAGCTGTCTACGACCAATTGAGCGGCCTCGGCACCGGGATTCTCGAT/3OH/ |
| **Nick-NCP–2** | |
| Oligo 1 | /5Phos/ATCGAGAATCCCGGTGCCGAGGCCGCTCAATTGGTCGTAGACAGCTCTAGCACCGCTTAAACGCACGTACGCGCTGTCCCCCGCGTTTTAACCGCCAAGGGGATTACTCCCTAGTCTCCAGGCACGTGTCAGATATATACATCCGAT/3OH/ |
| Oligo 2 | /6FAM/ATCGGATGTATATATCTGACACGTGCCTGGAGACTAGGGAGTAATCCCCTTGGCG/3Phos/ |
| Oligo 3 | /5OH/GTTAAAACGCGGGGGACAGCGCGTACGTGCGTTTAAGCGGTGCTAGAGCTGTCTACGACCAATTGAGCGGCCTCGGCACCGGGATTCTCGAT/3OH/ |

*Nucleosomes used for cryo-EM were generated with oligonucleotides containing a 5Phos instead of the 6FAM label.

**Table S5: Oligonucleotides for generating NCPs**

| Oligo | Sequence (5′ – 3′) |
|---|---|
| **Nick-NCP−1/Nick-DNA** | |
| Oligo 1 | **/5Phos/**ATCGAGAATCCCGGTGCCGAGGCCGCTCAATTGGTCGTAGACAGCTCTAGCACCGCTTAAACGCACGTACGCGCTGTCCCCCGCGTTTTAACCGCCAAGGGGATTACTCCCTAGTCTCCAGGCACGTGTCAGATATATACATCCGAT**/3OH/** |
| Oligo 2 | **/6FAM/**ATCGGATGTATATATCTGACACGTGCCTGGAGACTAGGGAGTAATCCCCTTGGCGGTTAAAACGC**/3Phos/** |
| Oligo 3 | **/5OH/**GGGGGACAGCGCGTACGTGCGTTTAAGCGGTGCTAGAGCTGTCTACGACCAATTGAGCGGCCTCGGCACCGGGATTCTCGAT**/3OH/** |
